# Biochemical characterisation of fungal bioluminescence enzymes reveals substrate inhibition and secondary turnover of 3-hydroxyhispidin by H3H

**DOI:** 10.64898/2026.09.09.750069

**Authors:** Jonathan F. Burnett, Johannes Lang, Anthony Tumber, Harold W. Mackenzie, Iryna Poplevicheva, Isaac X.R. Chan, Christopher Watson, Patrick Rabe

## Abstract

The fungal bioluminescence pathway has emerged as a versatile platform for bioimaging and synthetic biology. However, biochemical characterisation of its core enzymes remains limited. Here, we establish recombinant purification and quantitative *in vitro* characterisation of fungal luciferase (Luz) and hispidin-3-hydroxylase (H3H) homologues from *Neonothopanus nambi* and *Mycena chlorophos*. Complementary luminescence and mass spectrometry-based assays revealed pronounced substrate inhibition of H3H by hispidin, conserved across both homologues. Direct monitoring of substrate turnover further revealed an unexpected secondary H3H-catalysed conversion of 3-hydroxyhispidin that remained dependent on NADPH and FAD. Complementary NMR, isotope labelling and product characterisation studies identified multiple downstream products, including caffeic acid, while kinetic analysis demonstrated a strong preference of H3H for hispidin over 3-hydroxyhispidin. Together, these findings reveal previously unrecognised catalytic complexity within the fungal bioluminescence pathway and provide a biochemical framework for understanding and engineering pathway flux.

## Introduction

Bioluminescence has evolved independently across multiple kingdoms of life and is widely used in biological research, biotechnology, and bioimaging (1–5). Among known luminescent systems, fungal bioluminescence is unique in that light production arises from a metabolically integrated pathway derived from caffeic acid, a common intermediate of phenylpropanoid metabolism (6, 7). This enables autonomous light emission without continuous exogenous luciferin supplementation in heterologous hosts capable of endogenous caffeic acid biosynthesis, such as most plants (7–9).

The chemical basis of fungal bioluminescence remained unresolved for several decades despite early demonstrations of luminescence in cell free extracts (10–12). Identification of 3-hydroxyhispidin, the fungal luciferin, and hispidin, its biosynthetic precursor, established the biochemical framework of the fungal bioluminescence pathway (FBP) (13). Subsequent studies identified the core enzymes of the FBP in *Neonothopanus nambi*, including hispidin synthase (HispS), hispidin-3-hydroxylase (H3H), luciferase (Luz), and caffeoylpyruvate hydrolase (CPH), establishing a genetically encodable biosynthetic cycle linking luciferin production, light emission and metabolite recycling (Fig. 1) (6). Light emission occurs through oxidation of 3-hydroxyhispidin catalysed by Luz, involving a proposed high-energy endoperoxide intermediate that decomposes to form the oxyluciferin caffeoylpyruvate with concomitant photon emission (14).

**Figure 1:**
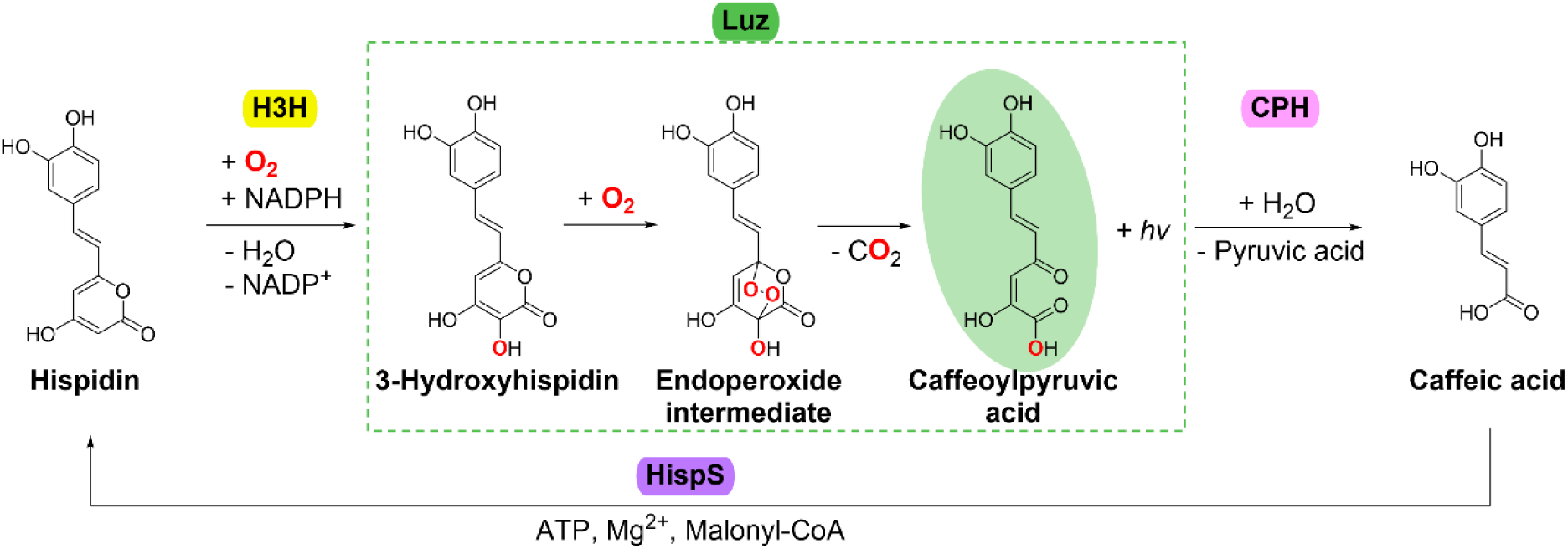
Fungal Bioluminescence Pathway. Schematic view of the fungal bioluminescence pathway. Hispidin is converted to 3-hydroxyhispidin by H3H. Oxidation by luciferase (Luz) produces light and caffeoylpyruvic acid, which is hydrolysed by caffeoylpyruvate hydrolase (CPH) to caffeic acid. Caffeic acid is then converted by hispidin synthase (HispS) back to hispidin, completing the fungal bioluminescence pathway (6).

The elucidation of the FBP enabled its deployment in heterologous hosts, with light emission demonstrated in plants, fungi, mammalian cells, and other model systems (6, 7, 15–24). The potential for autonomous light emission in suitable hosts represents a key advantage of the FBP over other systems requiring exogenous luciferin supplementation. Recent engineering efforts have improved light output through directed evolution and combinations of enzymes from different fungal species (8, 25).

Despite growing interest in its technological applications, the biochemical and structural properties of the FBP enzymes remain incompletely characterised. Recent biochemical studies on CPH have suggested an additional role, beyond metabolite recycling, in maintaining pathway flux by alleviating oxyluciferin-mediated inhibition of luciferase (26, 27). Biochemical characterisation of H3H, the flavin-dependent monooxygenase catalysing the conversion of hispidin to 3-hydroxyhispidin, remains limited. Previous studies have reported challenges in recombinant expression and purification of nnH3H from *N. nambi*, including aggregation, poor solubility, and dependence on fusion tags and molecular chaperones (26). In comparison, mcH3H from *Mycena chlorophos* showed more favourable recombinant expression and purification properties (27). To date, most functional H3H characterisation has relied upon oxygen consumption measurements or cofactor utilisation assays with little attention given to the conversion of hispidin and formation or subsequent turnover of 3-hydroxyhispidin (8, 25–29). Similarly, structural and functional aspects of the fungal luciferase Luz remain incompletely characterised. Bioinformatic analyses predict an N-terminal membrane-associated region, and recombinant enzyme has generally been recovered from membrane fractions (26, 30). However, the functional significance of this region remains unclear, and mutagenesis studies have so far provided only limited insight into residues important for enzymatic activity (8, 31).

These limitations complicate mechanistic investigation and rational engineering of the pathway and highlight the need for purified, well-behaved enzymes and quantitative assays capable of resolving individual catalytic steps. Furthermore, systematic biochemical comparisons of H3H and Luz using purified enzymes remain limited, while potential secondary catalytic activities and regulatory features of the FBP have received comparatively little attention.

In this study, we establish reliable purification strategies for recombinant Luz and H3H homologues from *N. nambi* and *M*. *chlorophos.* Using complementary luminescence-based, mass spectrometric and NMR analyses, we characterise their catalytic properties using direct and orthogonal assays. We further identify pronounced substrate inhibition during H3H catalysis and provide evidence for an additional H3H-catalysed conversion of 3-hydroxyhispidin. These findings contribute to our fundamental understanding of the biochemical properties of the fungal bioluminescence pathway enzymes and may inform future mechanistic, structural, and engineering studies.

## Results

### Expression and purification of fungal luciferase and H3H homologues

Full-length and truncated luciferase constructs from *N. nambi* (Luz_FL_ and Luz_Tr_ (Δ1-37), respectively) were recombinantly expressed in *E. coli*, together with an optimised truncated variant initially identified by directed evolution (Luz_Tr_ v4 with substitutions I63T, T99P, T192S and A199P) (Fig. S1) (8). Hispidin-3-hydroxylase (H3H) homologues from *N. nambi* (nnH3H) and *M. chlorophos* (mcH3H) were expressed in parallel (Fig. S2). Constructs were designed with N- or C-terminal affinity tags to facilitate purification (Figs. S1-S4).

Both Luz_Tr_ and Luz_Tr_ v4 were recovered as catalytically active enzymes but remained associated with the membrane fraction, requiring detergent solubilisation prior to affinity purification (Figs. S4A and B). N-terminally tagged Luz_Tr_ was retained on the nickel affinity column only following solubilisation with CHAPS but not DDM, whereas the C-terminally tagged Luz_Tr_ v4 could be purified using DDM or CHAPS, followed by reverse IMAC and size-exclusion chromatography, yielding a homogeneous and catalytically active enzyme (Figs. S3 and S4). The same C-terminal purification strategy was applied to full-length Luz, yielding active protein; however, purification of active full-length enzyme from the membrane fraction to a quality suitable for kinetic characterisation was unsuccessful, and the samples lost activity following freeze-thaw cycles.

Expression of nnH3H required an N-terminal SUMO tag together with co-expression of the molecular chaperones GroEL/ES or Trigger Factor to obtain soluble protein (Fig. S4C-E) (26). In contrast to previous reports describing limited or unstable activity during purification (26), purified nnH3H retained measurable catalytic activity under the conditions employed when supplemented with FAD. However, inefficient tag cleavage and partial protein heterogeneity consistent with oligomerisation limited recovery of cleaved protein, and nnH3H was therefore retained as the SUMO fusion (SUMO nnH3H) for subsequent biochemical characterisation. In contrast, mcH3H was readily expressed and purified in good yield without the need for chaperone co-expression (Fig. S4F) (27). Inclusion of reducing agent (TCEP) during purification improved protein stability, and mcH3H could be obtained as a cleaved, homogeneous preparation following affinity purification and size-exclusion chromatography.

All purified enzymes retained catalytic activity. Luz_Tr_ v4 exhibited substantially increased luminescent output relative to Luz_Tr_ upon addition of 3-hydroxyhispidin (Fig. 2). NADPH-dependent turnover of hispidin by both H3H homologues was subsequently characterised using coupled luminescence assays with Luz_Tr_ v4 and orthogonal mass spectrometry-based assays.

**Figure 2:**
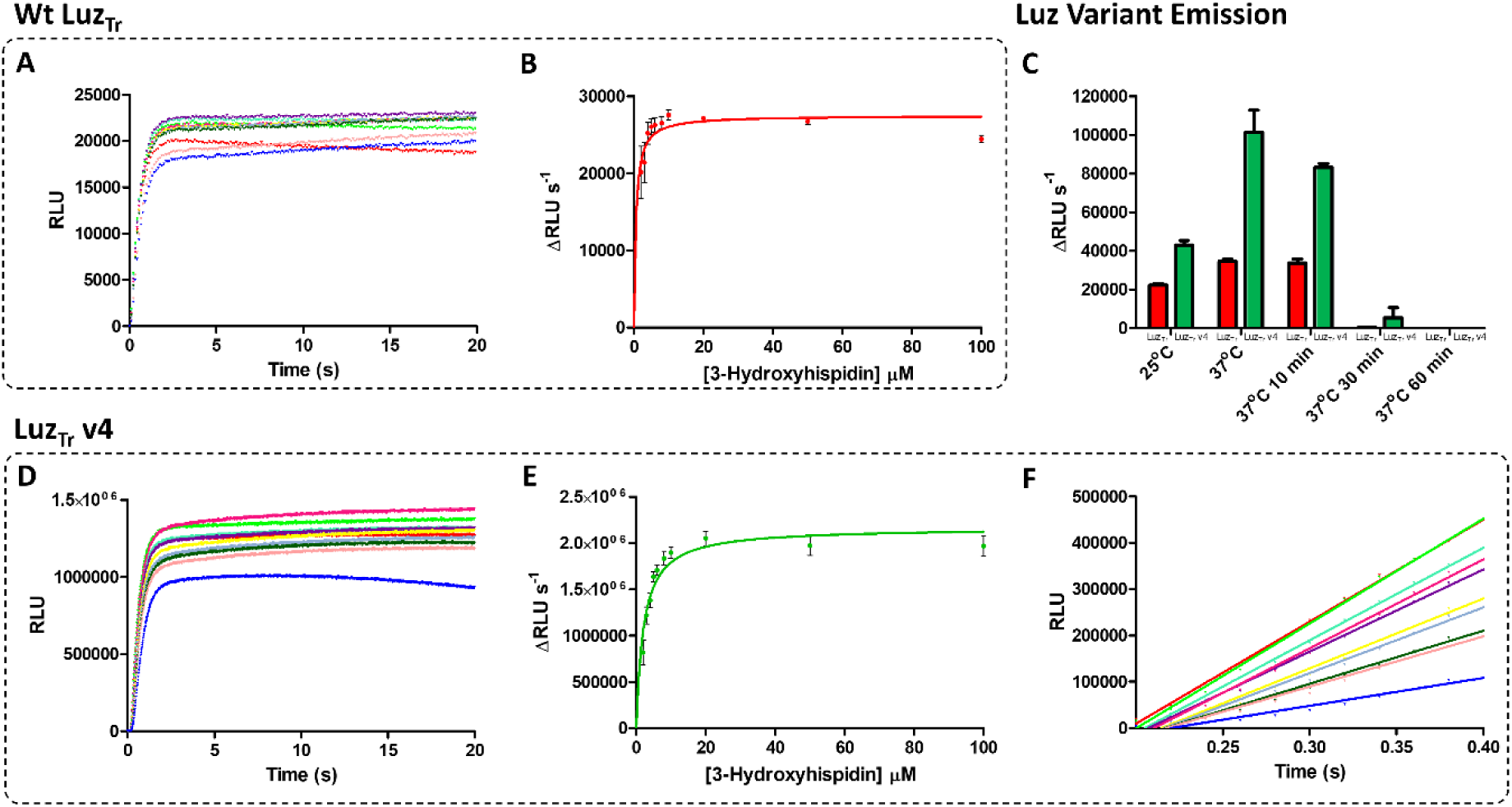
Luciferase steady-state kinetics and comparison of wildtype and v4 variants. (**A**) Progress curves describing the reaction between LuzTr (200 nM) and 3-hydroxyhispidin (red: 100, green: 50, teal: 20, purple: 10, pink: 8, yellow: 6, grey: 5, dark green: 4, coral: 3, blue: 2 µM). (**B**) Michaelis-Menten kinetic fit from the initial rate of RLU change (ΔRLU s^-1^) from LuzTr progress curves. (**C**) Comparison of the initial rates of luminescence generated by LuzTr (red, 200 nM) and LuzTr v4 (green, 200 nM) following addition of 3-hydroxyhispidin (100 µM). Enzymes were assayed at 25 °C and 37 °C and following incubation at 37 °C for 10, 30 and 60 min to assess temperature dependent loss of activity. (**D**) Progress curves describing the reaction between Luz_Tr_ v4 (200 nM) and 3-hydroxyhispidin (red: 100, green: 50, teal: 20, purple: 10, pink: 8, yellow: 6, grey: 5, dark green: 4, coral: 3, blue: 2 µM). (**E**) Michaelis-Menten kinetic fit from the initial rate of RLU change (ΔRLU s^-1^) from Luz_Tr_ v4 progress curves. (**F**) Linear regression fit of initial rate of reaction of purified Luz_Tr_ v4 (200 nM) with 3-hydroxyhispidin from (**D**). Luminescence-based plate reader assays were performed in sodium phosphate buffer (200 mM NaH₂PO₄/Na₂HPO₄, 500 mM Na_2_SO_4_, 0.1% DDM, pH 8.0). Results were collected in triplicate (n=3; mean ± SD). Luz_Tr_, (38-267) luciferase; RLU, relative luminescent units.

### Kinetic characterisation of truncated and full-length luciferase

The activities of Luz_Tr_ and Luz_Tr_ v4 were characterised *in vitro* to compare their luminescent properties and define conditions for subsequent coupled enzymatic assays. Addition of 3-hydroxyhispidin resulted in rapid light emission, with the luminescent signal approaching a plateau within the first seconds following substrate addition. The initial increase in luminescence was quantified over the first 0.2–0.4 s following simultaneous substrate injection and data acquisition using a CLARIOstar plate reader.

To assess the influence of substrate dispensing and mixing on the initial rate, reactions were initiated at dispensing speeds between 50 and 400 µL s^-1^ (Fig. S5D). The initial rate increased concomitantly with dispensing speed up to approximately 200 µL s^-1^ but remained essentially unchanged at higher speeds. All subsequent kinetic measurements were performed with dispensing at 400 µL s^-1^, ensuring that dispensing and mixing were not rate-limiting under the conditions used.

The dependence of the luminescence response on Luz_Tr_ v4 concentration was examined between 20 and 200 nM. Michaelis-Menten analysis of the initial luminescence increase gave apparent K_M_ values in the low micromolar range although the fitted K_M_ decreased with increasing enzyme concentration (Fig. S5). Analysis based on maximal luminescence showed a stronger enzyme-concentration dependence. We therefore interpret the luminescence-derived parameters as apparent kinetic values describing the rapid light-emitting reaction rather than rigorous steady-state kinetic constants. Previous studies on other luciferases have used both the initial light response and integrated luminescence to characterise luciferase activity (32–36). Here, we focused on the initial light response to characterise the rapid light-emitting reaction, while direct substrate turnover was assessed independently by SPE-MS.

Kinetic analysis revealed a low micromolar apparent Michaelis constant for 3-hydroxyhispidin for Luz_Tr_ (K_M_ = 0.57 µM) (Fig. 2, Table 1). The mutations introduced to generate Luz_Tr_ v4 had a modest effect on the apparent K_M_ (2.12 µM). Notably, under identical assay conditions, Luz_Tr_ v4 displayed enhanced luminescent output compared to Luz_Tr_ (Fig. 2C), which may reflect differences in catalytic turnover, quantum yield, or both. Purified Luz constructs exhibited robust activity over a broad pH range, with maximal rates observed between pH 8.0 and 9.0 (Fig. S6A), in agreement with previous studies on Luz_FL_ in cell lysates (30). Luminescent output remained stable throughout the assay (Fig. S7C), however gradual non-enzymatic degradation of 3-hydroxyhispidin in solution was observed over time, directly shown by NMR (Fig. S8B).

**Table 1.** Steady-state kinetic parameters of H3H and Luz variants determined using orthogonal assays. Kinetic parameters were calculated from initial rate measurements using nonlinear regression in GraphPad Prism. H3H kinetics were measured using coupled luminescence and SPE-MS assays, whereas Luz kinetics were determined using luminescence or SPE-MS as indicated. Coupled, coupled H3H-Luz luminescence assay; Lum, luciferase luminescence assay; SPE-MS, solid phase extraction coupled to mass spectrometry assay. H3H, Hispidin-3-hydroxylase; LuzTr, (38-267) luciferase.

| Enzyme |  | Hispidin-3-hydroxylase |  |  |  | Luciferase |  |  |
|---|---|---|---|---|---|---|---|---|
|  |  | nnH3H |  | mcH3H |  | Luz<sub>Tr</sub> | Luz<sub>Tr</sub> v4 | Luz Fl |
| Assay |  | Coupled | SPE-MS | Coupled | SPE-MS | Luminescence |  |  |
| OH-Hispidin<br>K<sub>M</sub> (µM) |  | - | 472 ± 144 | - | 109 ± 49.4 | 0.57 ± 0.19 | 2.12 ± 0.40 | N.D |
| Hispidin | K<sub>M</sub> (µM) | 0.69 ± 0.35 | 3.10 ± 1.17 | 2.80 ± 0.15 | 6.61 ± 1.84 | - | - | - |
|  | K<sub>I</sub> (µM) | 106.4 ± 32.9 | 19.06 ± 7.96 | 126.2 ± 7.84 | 26.6 ± 5.23 |  |  |  |
| NADPH K<sub>M</sub> (µM) |  | 34.9 ± 6.38 | 21.8 ± 6.6 | 114.6 ± 14.1 | 72.7 ± 23.7 | - | - | - |

While luminescence measurements provided a highly sensitive readout of luciferase activity, relative light units do not directly quantify substrate turnover. Solid-phase extraction mass spectrometry (SPE-MS) was therefore used to directly monitor 3-hydroxyhispidin consumption and provide an orthogonal measure of luciferase activity independent of photon production. DDM was omitted from the assay buffer to ensure compatibility with SPE-MS. Under these conditions, plate reader controls retained substantial activity, although the initial rate and maximal emission were reduced by 37% and 24%, respectively (Fig. S9D). SPE-MS analysis yielded an apparent K_M_ for 3-hydroxyhispidin of K_M_ = 118 µM (Fig. S9E), substantially higher than that determined from the luminescence assay. However, the limited temporal resolution of SPE-MS (approximately 14.5 s sampling interval) prevents capture of the rapid initial phase observed by luminescence. In addition to that, omission of DDM affected luciferase activity, further complicating direct quantitative comparison between the two methods.

Comparison of Luz_Tr_ and Luz_Tr_ v4 demonstrated that the increased luminescent output of the engineered variant was not associated with a lower apparent K_M_ for 3-hydroxyhispidin. Luz_Tr_ v4 also showed improved retention of activity following incubation at 37 °C (Fig. 2C), indicating increased stability relative to Luz_Tr_ (8). However, the relative contributions of catalytic turnover and quantum yield to the increased luminescent output could not be distinguished from these measurements. Direct comparison of substrate turnover by SPE-MS was not possible due to the limited stability and recovery of purified Luz_Tr_. Together, the increased luminescent output and improved stability of Luz_Tr_ v4 supported its selection as the reporter enzyme for subsequent coupled assays monitoring H3H-catalysed turnover of hispidin.

### H3H displays substrate inhibition by hispidin

Previous studies on H3H activity have primarily relied on monitoring NADPH consumption and oxygen depletion, rather than directly tracking the turnover of the caffeic acid-derived substrate hispidin and the formation of its corresponding product in real time (27). We therefore developed a coupled luminescence assay in which H3H-catalysed formation of 3-hydroxyhispidin was monitored through its subsequent conversion by Luz_Tr_ v4 (Fig. 3). Initial experiments were performed using 200 nM SUMO nnH3H and 2 µM Luz_Tr_ v4, which provided a robust luminescent readout of H3H activity. For subsequent kinetic measurements performed at lower H3H concentrations, the required Luz_Tr_ v4 concentration was experimentally assessed to ensure that reporter activity did not limit the measured H3H rate. Under these conditions, increasing Luz_Tr_ v4 concentrations reached a plateau in the measured rate, supporting the reporter concentrations selected for kinetic analysis (Fig. S10).

**Figure 3:**
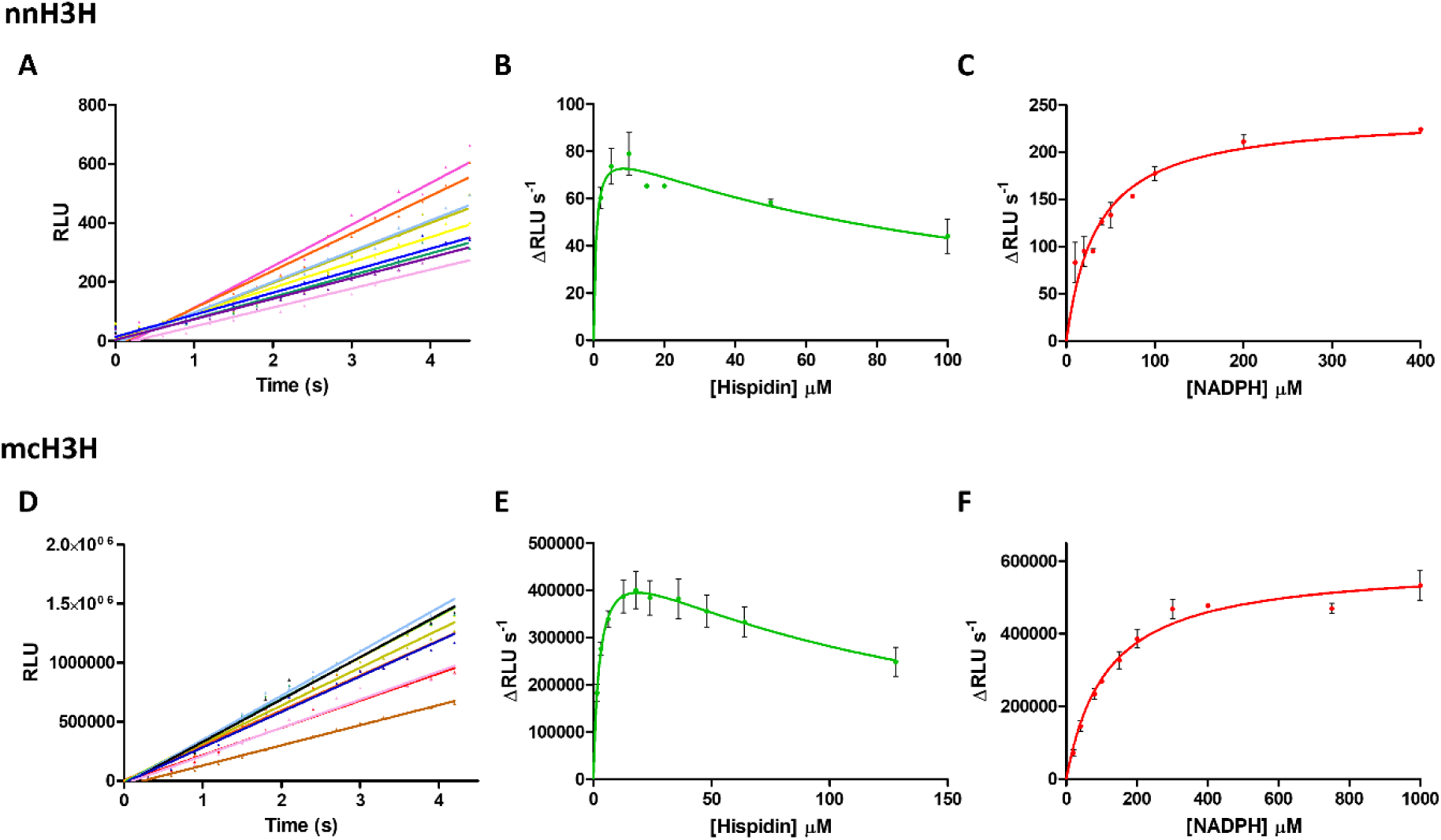
Coupled luciferase assay reveals hispidin substrate inhibition of H3H. Steady-state kinetic analysis of SUMO nnH3H (A -C) and mcH3H (D - F) using a coupled luminescence assay in which excess LuzTr v4 reports formation of 3-hydroxyhispidin. Initial rates (ΔRLU s^-1^) were determined from the early linear phase of the progress curves and fitted to Michaelis-Menten or substrate inhibition models, as appropriate. (**A**) Representative progress curves using SUMO nnH3H (50 nM), LuzTr v4 (10 µM) and FAD (0.5 µM) recorded at fixed hispidin (10 µM) with increasing NADPH concentrations: pale pink, 10 µM; purple, 20 µM; blue, 30 µM; light blue, 40 µM; green, 50 µM; yellow, 75 µM; sand, 100 µM; orange, 150 µM; pink, 200 µM. (**B**) SUMO nnH3H (50 nM) initial rate analysis at fixed NADPH (200 µM) with varying hispidin concentrations (2 - 100 µM), FAD (0.5 µM ) and LuzTr v4 (10 µM). (**C**) Initial rate analysis using SUMO nnH3H (50 nM), LuzTr v4 (10 µM) and FAD (0.5 µM) at fixed hispidin (10 µM) with varying NADPH concentrations (10 - 400 µM). (**D** - **F**) mcH3H (20 nM) was used to initiate the reaction with hispidin and NADPH in the presence of LuzTr v4 (10 µM). (**D**) Representative progress curves recorded at fixed NADPH (200 µM) with increasing hispidin concentrations: brown, 1.6 µM; pink, 3.2 µM; blue, 6.4 µM; black, 12.8 µM; light blue, 18 µM; green, 24 µM; yellow, 36 µM; sand, 48 µM; orange, 64 µM; red, 128 µM. (**E**) Initial rate analysis at fixed NADPH (200 µM) with varying hispidin concentrations (1.6 - 128 µM). (**F**) Initial rate analysis at fixed hispidin (6.4 µM) with varying NADPH concentrations (10 – 1000 µM). All reactions were performed in 50 mM Tris HCl, 150 mM NaCl, pH 8.0. Results were collected in triplicate (n=3; mean ± SD). H3H, hispidin-3-hydroxylase; Luz_Tr_, (38-267) luciferase; RLU, relative luminescent units.

H3H-dependent luminescence required both NADPH and FAD, consistent with the established cofactor requirements of H3H as a flavin-dependent monooxygenase (27, 37). Supplementation with FAD was required for optimal activity of purified SUMO nnH3H, consistent with partial cofactor loss during purification. Both H3H homologues displayed maximal apparent activity under the coupled assay conditions at a slightly higher pH than previously reported for mcH3H (Fig. S6B+C) (27). Unexpectedly, increasing hispidin concentrations beyond the low micromolar range resulted in a progressive decrease in luminescent output (Fig. 3). Analysis of initial rates revealed pronounced substrate inhibition of H3H, with maximal enzyme activity observed at low micromolar hispidin concentrations (Fig. 3B, Table 1). This behaviour was reproducible across independent experiments and was not attributable to hispidin instability (Fig. S8A). Although hispidin also inhibited Luz_Tr_ v4 at higher concentrations, the magnitude of this effect was insufficient to account for the inhibition observed in the coupled H3H assay (Fig. S9B).

Kinetic analysis of SUMO nnH3H yielded an apparent K_M_ = 0.69 µM for hispidin. At higher hispidin concentrations, activity decreased in a manner consistent with substrate inhibition, yielding an apparent K_I_ = 106.4 µM (Fig. 3A+B, Table 1). In contrast, variation of NADPH concentration at fixed hispidin concentrations followed classical Michaelis-Menten behaviour, with an apparent K_M_(NADPH) = 34.9 µM (Fig. 3C). Similar behaviour was observed for mcH3H, with the apparent K_M_ = 2.8 µM and K_I_= 126.2 µM for hispidin (Fig. 3D+E), while variation of NADPH again followed Michaelis-Menten behaviour, with an apparent K_M_(NADPH) = 114.6 µM (Fig. 3F). Consistent with previous reports for mcH3H and the broader cofactor flexibility of FAD-dependent monooxygenases (27), NADH could also serve as a cofactor for both H3H homologues (Fig. S9F), although with lower catalytic efficiency than NADPH under otherwise identical assay conditions (Table 1).

### Orthogonal mass spectrometry reveals turnover of 3-hydroxyhispidin

Having established SPE-MS for direct monitoring of 3-hydroxyhispidin turnover by Luz_Tr_ v4, we next extended this approach to characterise H3H activity independently of the coupled luminescence assay and directly monitor hispidin and 3-hydroxyhispidin turnover. Initial experiments focused on optimisation of cartridge usage and assay conditions to enable reliable detection of both hispidin and 3-hydroxyhispidin. Alternative stationary phases were screened; optimal retention, recovery, and signal stability for both hispidin and 3-hydroxyhispidin were achieved using a C18 cartridge, while a graphitic carbon cartridge provided improved retention of cofactors and putative side products. This setup enabled reproducible detection of all relevant species across the full reaction time course.

Using the optimised SPE-MS setup, hispidin consumption was monitored over time, confirming efficient substrate turnover by H3H. Apparent kinetic parameters derived from SPE-MS analysis were broadly consistent in magnitude with those obtained from the coupled luminescence assay, although differences were observed between the two approaches (Fig. 4A, B, D, E and Table 1), providing orthogonal support for the kinetic behaviour observed using the coupled assay. For SUMO nnH3H, observed K_M_ values for NADPH (K_M_ = 21.8 µM) and hispidin (K_M_ = 3.10 µM) were of a similar magnitude to those obtained using the coupled luciferase assay (K_M_(NADPH) = 34.9 µM and K_M_(hispidin) = 0.69 µM). Similarly, for mcH3H, observed K_M_ values for NADPH (K_M_ = 72.7 µM) and hispidin (K_M_ = 6.61 µM) were of a similar magnitude to those obtained using the coupled luciferase assay (K_M_(NADPH) = 114.6 µM and K_M_(hispidin) = 2.80 µM).

**Figure 4:**
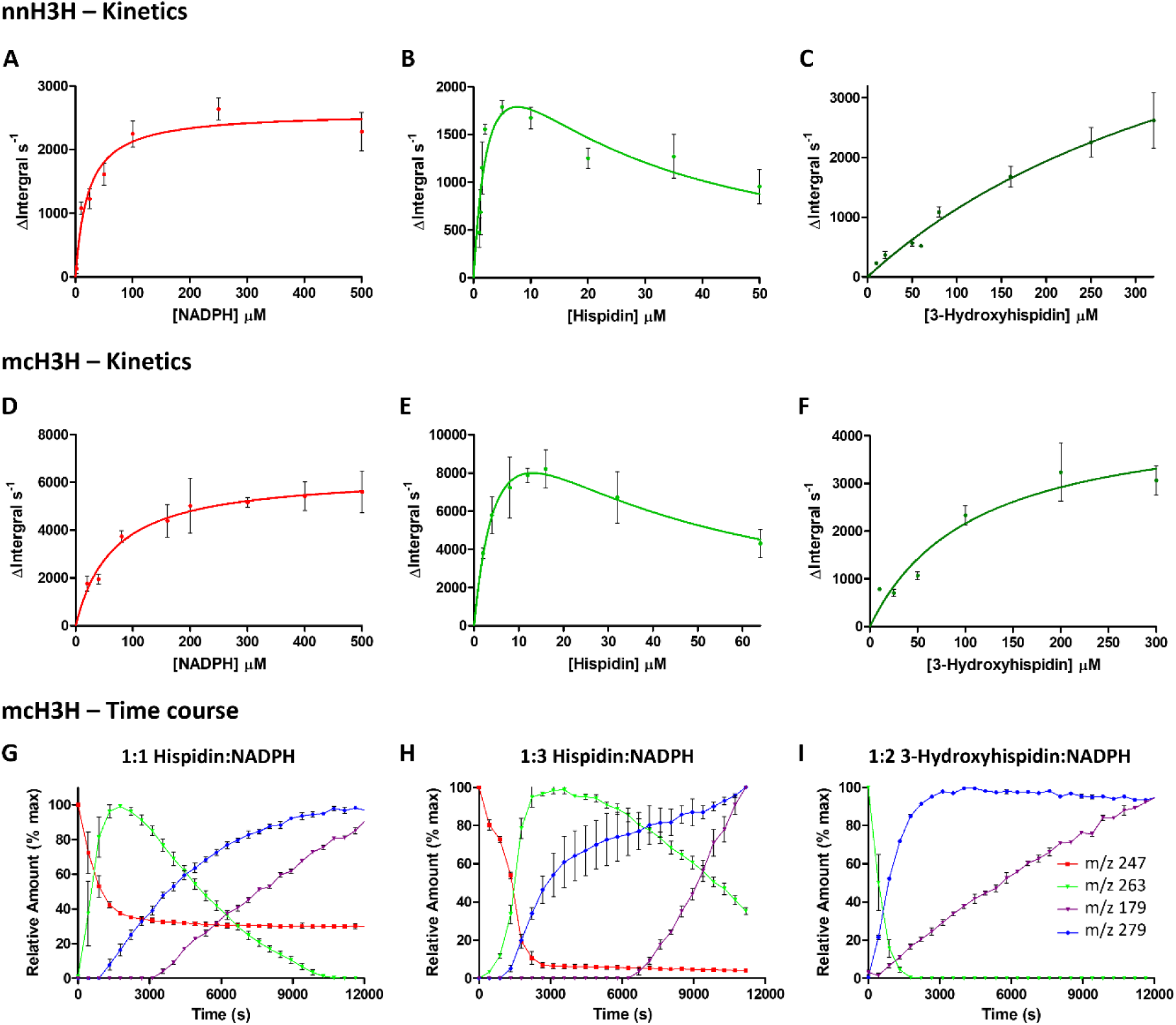
SPE-MS steady-state kinetics of SUMO nnH3H and mcH3H. SUMO nnH3H and mcH3H were analysed using SPE-MS. Steady-state kinetic parameters were determined from the initial decrease in ion integral corresponding to consumption of hispidin at m/z = 247.0359 ([M+H]^+^) or 3-hydroxyhispidin at m/z = 263.0281 ([M+H]^+^). **SUMO nnH3H kinetics** (**A** - **C**): SUMO nnH3H (200 nM) was premixed with FAD (5 µM) and used to initiate reactions containing: (**A**) hispidin (10 µM) with NADPH (2 – 500 µM); (**B**) NADPH (100 µM) with hispidin (1 – 50 µM); (**C**) NADPH (500 µM) with 3-hydroxyhispidin (2 – 320 µM). **mcH3H kinetics** (**D** - **F**): mcH3H (10 nM) used to initiate the reaction containing: (**D**) hispidin (10 µM) with NADPH (20 – 500 µM); (**E**) NADPH (200 µM) with hispidin (2 - 64 µM); (**F**) NADPH (500 µM) with 3-hydroxyhispidin (10 – 300 µM). **mcH3H time-courses** (**G** - **I**): Turnover of hispidin and 3- hydroxyhispidin by mcH3H (500 nM) was monitored over an extended period at different substrate:NADPH ratios: (**G**) hispidin (200 µM) and NADPH (200 µM; 1:1); (**H**) hispidin (200 µM) and NADPH (600 µM; 1:3); (**I**) 3-hydroxyhispidin (200 µM) and NADPH (400 µM; 1:2). Monitored [M+H]^+^ ions were m/z 247 (hispidin, red), 263 (3-hydroxyhispidin, green), 179 (putative side product, purple) and 279 (putative side product, blue). All reactions were performed in 50 mM Tris-HCl, 150 mM NaCl, pH 8.0. C18 cartridges were used for steady-state kinetic measurements and graphitic cartridges for extended time course experiments. Data represent the mean of three independent repeats (n=3, mean ± SD). H3H, Hispidin-3-hydroxylase; SPE-MS, solid phase extraction coupled to mass spectrometry.

Direct measurement by SPE-MS additionally enabled determination of apparent *k*_cat_ values of *k*_cat_(NADPH) = 4.5 s^-1^ and *k*_cat_(hispidin) = 13.1 s^-1^ for mcH3H. These values were obtained by independently varying NADPH or hispidin while maintaining the second substrate at a fixed concentration and therefore represent apparent turnover parameters under the respective assay conditions. The difference between these apparent *k*_cat_ values may partially reflect the different fixed substrate concentrations used, together with substrate inhibition by hispidin. The greater variation in K_M_ observed for hispidin may reflect its rapid consumption at low substrate concentrations, which, combined with the approximately 14.4 s SPE-MS sampling interval, limited the number of data points captured along the ascending portion of the kinetic curve. The kinetic constants describing NADPH turnover by mcH3H were also of a similar magnitude to previously reported values (27). In comparison, the reaction of SUMO nnH3H displayed lower apparent catalytic efficiencies under the conditions tested than mcH3H (Table 1), suggesting future biotechnological development of recombinant pathways may benefit from a multispecies approach.

Notably, and consistent with the behaviour observed in the coupled assay (Fig. S7C), accumulation of 3-hydroxyhispidin was not sustained (Fig. S7A). Instead, following an initial increase, product levels decreased over time. This behaviour was reproducible across independent experiments. In the absence of active enzyme, no significant depletion was observed over the time of the experiment (Fig. S7B) indicating that 3-hydroxyhispidin does not accumulate under catalytic conditions but is instead further processed enzymatically in the presence of H3H. Furthermore, limiting NADPH to stoichiometric concentrations (1 eq.) resulted in incomplete depletion of hispidin, alongside a decrease of 3-hydroxyhispidin (Fig. 4G). Together, these observations suggest that, in the absence of luciferase-mediated depletion, 3-hydroxyhispidin undergoes further NADPH-dependent turnover in the presence of H3H.

### H3H catalyses turnover of 3-hydroxyhispidin

To determine whether the observed decrease in 3-hydroxyhispidin arises from enzymatic turnover, 3-hydroxyhispidin was incubated with both H3H homologues under catalytic conditions (Fig. 4C and F). Both SUMO nnH3H and mcH3H catalysed turnover of 3-hydroxyhispidin, indicating that this activity is conserved across the two H3H homologues. In the presence of SUMO nnH3H, NADPH, and FAD, rapid depletion of 3-hydroxyhispidin was observed, whereas no significant change occurred in control reactions lacking enzyme or cofactors (Fig. S7B and D). Under SPE-MS conditions, two additional species with m/z = 179 and m/z = 279 accumulated over the course of the reaction. The formation of these species was more rapid when 3-hydroxyhispidin was supplied directly (Fig. 4I) than when it was formed through initial turnover of hispidin (Fig. 4G and H), particularly for the m/z = 179 species. This temporal relationship suggests that these species arise downstream of 3-hydroxyhispidin, although their formation may include both enzymatic and non-enzymatic processes.

To further probe 3-hydroxyhispidin turnover, 3-hydroxyhispidin was pre-incubated with SUMO nnH3H for extended periods prior to addition of luciferase. Under these conditions, luminescence decreased with increasing preincubation time, consistent with a reduction in the amount of 3-hydroxyhispidin available for subsequent luciferase turnover. Addition of fresh 3-hydroxyhispidin restored luminescence, confirming that the reduced signal was not caused by loss or inhibition of luciferase activity (Fig. S7C). Together, these results provide orthogonal evidence that 3-hydroxyhispidin undergoes further H3H-dependent turnover.

### NMR analysis reveals sequential turnover and formation of secondary products

Given its improved activity, stability and higher expression yield, mcH3H was used for further characterisation. To directly monitor reaction progression at the molecular level, NMR-based turnover assays were performed (Fig. 5). When hispidin was incubated with mcH3H in the presence of excess NADPH (3 eq.), complete depletion of the substrate was observed, as evidenced by the disappearance of the doublet corresponding to the proton at pyrone *C*-3 (δ 5.05 ppm). Concurrent formation of 3-hydroxyhispidin was shown by the loss of the resonances at δ 6.60 (vinylic proton adjacent to pyrone) and 5.92 ppm (proton at pyrone *C*-5) and the appearance of the corresponding product signals at δ 6.56 and 6.02 ppm, respectively. Upon prolonged incubation, 3-hydroxyhispidin was subsequently consumed as shown by disappearance of vinylic and pyrone proton resonances. This was accompanied by continued production of NADP^+^ (Fig. 5, Figs. S11-13), further indicating that the second transformation step is also NADPH-dependent, corroborated by coupled assay controls (Fig. S7C and D). NMR time-course experiments comparing different hispidin:NADPH ratios further supported this dependence: under excess NADPH, initial formation of 3-hydroxyhispidin was followed by its depletion, whereas near-stoichiometric NADPH resulted in accumulation of 3-hydroxyhispidin with limited subsequent turnover (Fig. S12). Notably, a transient intermediate species was detected during the course of the reaction, accumulating following 3-hydroxyhispidin formation and subsequently decreasing upon prolonged incubation (Fig. 5b.iv and 5b.v). While the precise origin of this species remains unclear, its temporal behaviour suggests that it may arise from a secondary transformation under the reaction conditions. Concomitant with its depletion, the appearance of additional resonances was observed, indicating formation of downstream products. Further time-course controls comparing reactions containing active and heat-inactivated H3H demonstrated accelerated 3-hydroxyhispidin depletion in the presence of active enzyme for both H3H homologues (Figs. S14 and S15), further supporting the requirement for catalytically active H3H.

**Figure 5:**
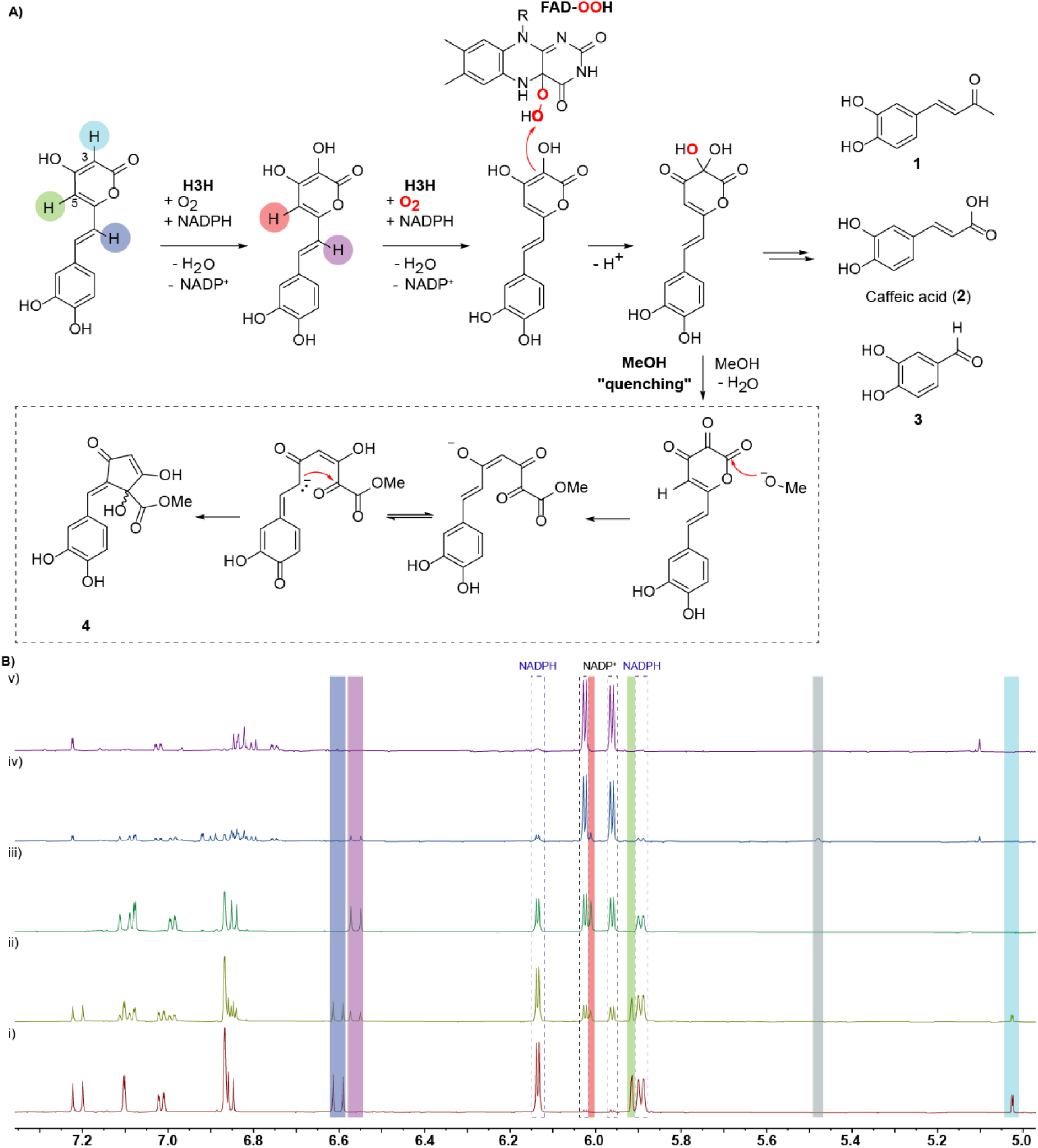
^1^H NMR analysis and proposed mechanism of secondary 3-hydroxyhispidin turnover by mcH3H. (**A**) Proposed model for sequential turnover of hispidin by mcH3H and subsequent product formation. The canonical mcH3H-catalysed hydroxylation produces 3-hydroxyhispidin through an NADPH, FAD and O2-dependent reaction. Secondary turnover involves an NADPH-dependent oxidative transformation of 3-hydroxyhispidin. One possible mechanism involving further flavin-dependent oxygenation at C-3 is shown, generating a transient geminal diol that may collapse to a C-3 carbonyl intermediate. Compounds **1**–**3** represent isolated downstream products; their routes of formation remain unresolved and may involve subsequent non-enzymatic transformations. Box: Methanol treatment resulted in formation of cyclopentenone **4**. One possible pathway starts from the proposed C3 carbonyl containing intermediate, with subsequent methanol addition promoting nucleophilic pyrone ring opening, rearrangement and cyclisation, ultimately yielding cyclopentenone 4. The proposed intermediates and individual reaction steps have not been experimentally established.(**B**) ^1^H NMR spectra monitoring mcH3H (0.5 µM) catalysed turnover of hispidin (250 µM) with NADPH (750 µM; 3 eq.) and FAD (0.5 µM) revealing secondary turnover of 3-hydroxyhispidin. (**i**) Prior to enzyme addition; (**ii**) 21 min after mcH3H addition; (**iii**) 36 min reaction in the NMR tube under limited O_2_ availability; (**iv**) 77 min following incubation in an Eppendorf tube with exposure to air; (**v**) following a further ∼2.5 h exposure to air in an Eppendorf tube. Highlighted resonances correspond to the proton positions indicated by matching colours in (a). The transient resonance highlighted in grey corresponds to an unidentified species formed during secondary turnover. Spectra were acquired in 25 mM Tris-d_11_, 100 mM NaCl, pD 8.0 in D_2_O using a Bruker AVIII 700 MHz spectrometer with water presaturation.

To characterise the products of secondary 3-hydroxyhispidin turnover, the mcH3H reaction was performed on a preparative scale in ammonium bicarbonate buffer using H_2_O or D_2_O as solvent under an O_2_ atmosphere. Reaction progress was monitored by ^1^H NMR spectroscopy and luminescence assay; following complete turnover, reactions were quenched with acetonitrile and products separated by HPLC for subsequent NMR and LCMS analysis.

Analysis of the HPLC fractions by NMR and LCMS revealed a complex mixture of products, several of which retained the characteristic catechol moiety of hispidin and 3-hydroxyhispidin. Compound **1** was identified as (3*E*)-4-(3,4-dihydroxyphenyl)-3-buten-2-one (Fig. 5), with its identity confirmed by comparison with a synthetic standard (Fig. S16). This compound is a naturally occurring fungal metabolite that has also been reported as a product of non-enzymatic oxyluciferin degradation (38, 39). Compound **2** was identified as caffeic acid (Fig. S17), an upstream intermediate of the fungal bioluminescence pathway. Compound **3** was identified as 3,4-dihydroxybenzaldehyde (Fig. S18), a previously reported degradation product of caffeic acid and related compounds (40). Additional products were also detected by LCMS, including a species consistent with dimerisation of downstream degradation products, highlighting the complexity of the reaction mixture, however the structures of these additional species were not determined.

To further investigate these downstream transformations, partially converted reaction mixtures were quenched and resuspended in methanol to limit enzymatic turnover. These experiments predominantly retained 3-hydroxyhispidin, consistent with incomplete turnover, but also revealed a distinct additional species that was subsequently characterised by NMR (Fig. S19) and identified as the previously unassigned cyclopentenone derivative **4**. The ^1^H NMR spectrum retained the characteristic catechol proton pattern, with resonances at δ_H_ 7.35 (d, *J* = 2.0 Hz), 7.15 (dd, *J* = 8.3, 2.0 Hz) and 6.94 (d, *J* = 8.3 Hz), together with a singlet at δ_H_ 7.03 assigned to a vinylic proton with its ^13^C signal at δ_C_ 127.7 ppm (Table S1). A methyl ester was supported by the singlet at δ_H_ 3.60 (3H) and corresponding δ_C_ 53.1 ppm. An additional singlet at δ_H_ 5.16 (^1^H) with a corresponding δ_C_ 102.5 ppm, exchanged in D_2_O and disappeared during HMBC acquisition (Fig. S19). Re-analysis of an isolated fraction containing this compound alongside an unidentified species in 90% H_2_O/10% D_2_O enabled acquisition of diagnostic HMBC correlations, including correlations from the δ_H_ 5.16 signal to δ_C_ 79.6, 136.4 and 198.1 ppm. The vinylic proton at δ_H_ 7.03 showed HMBC correlations to δ_C_ 79.3, 117.3, 127.7, 135.4 and 195.6 ppm (Table S1). Together with the high-resolution mass spectrometry data (m/z = 293.0681 Da) supporting the molecular formula C_14_H_12_O_7_, these data support assignment of the five-membered cyclopentenone **4** containing a methyl ester and exocyclic alkene (Fig. S19). Cyclopentenones are established secondary metabolites in the fungal metabolome, and can arise through transformations of highly conjugated systems (41, 42).

### Labelling experiments

To investigate the origin of oxygen incorporated during H3H catalysis, reactions were performed under an ^18^O_2_ atmosphere using a Schlenk-line setup previously established for oxygen isotope labelling experiments (43–45). Reactions were performed with either hispidin or 3-hydroxyhispidin as substrate at different substrate-to-NADPH ratios, with corresponding H_218_O controls used to assess incorporation of solvent-derived oxygen.

In reactions containing hispidin and a 1:1 substrate-to-NADPH ratio, formation of 3-hydroxyhispidin was observed at m/z 263.054 under ^16^O_2_, with a corresponding shift to m/z 265.058 under ^18^O_2_ (Fig. S20), consistent with incorporation of one oxygen atom from molecular oxygen during the canonical H3H-catalysed hydroxylation. This shift was not observed in the corresponding H ^18^O control using ^16^O_2_ (Fig. S21).

In contrast, ^18^O was not retained in lower molecular mass products observed following prolonged secondary turnover, including compound **1** (Figs. S21–S23). As these products arise following substantial fragmentation or rearrangement of the 3-hydroxyhispidin scaffold, loss of the portion containing an incorporated oxygen cannot be excluded. The absence of ^18^O in these products is therefore not diagnostic of the mechanism of secondary turnover.

An additional species at m/z 382.113 was reproducibly observed in Tris-containing buffer (Fig. S23). Substitution of Tris with Tris-d_11_ shifted this signal from m/z 382.113 to m/z 388.150, consistent with retention of the six non-exchangeable deuterium atoms and supporting incorporation of a Tris-derived component into this species (Fig. S24). Under ^18^O_2_, isotope incorporation into this species was observed when hispidin was used as the starting substrate, whereas it remained unlabelled when 3-hydroxyhispidin was supplied directly. H ^18^O experiments additionally showed solvent-derived oxygen incorporation or exchange (Figs. S21, S25, S26). These condition-dependent labelling patterns do not permit an unambiguous assignment of the structure or route of formation of this species and do not establish whether additional oxygen incorporation occurs during secondary 3-hydroxyhispidin turnover.

### Proposed mechanism for secondary turnover

The observed secondary turnover is unlikely to arise from simple oxidation of the C3 hydroxy group to the corresponding carbonyl, as this reaction would not account for the continued consumption of NADPH observed during 3-hydroxyhispidin turnover (46). Consistent with this, NMR analysis demonstrated continued conversion of NADPH to NADP⁺ during 3-hydroxyhispidin turnover, even after depletion of the initial hispidin substrate (Fig. S12). These findings support an NADPH-dependent secondary turnover, which may involve further flavin-dependent oxygenation of 3-hydroxyhispidin. One possible mechanism involves hydroxylation at an already oxygenated carbon, generating a transient geminal diol (Fig. 5), which subsequently collapses to a carbonyl-containing intermediate with concomitant elimination of water. This pathway would be consistent with the observed NADPH dependence of the secondary turnover and would generate an oxidised intermediate susceptible to subsequent rearrangement and fragmentation.

The proposed oxygenation step could proceed through the C4a-hydroperoxyflavin intermediate characteristic of flavin-dependent monooxygenases (Fig. 5), with the highly oxygenated geminal diol intermediate potentially facilitating subsequent pyrone ring opening, rearrangement or fragmentation. However, the isotope-labelling experiments do not establish additional oxygen incorporation during secondary turnover. The absence of ^18^O in the smaller products is not mechanistically conclusive, as isotope loss during their extensive fragmentation cannot be excluded, while the structure and route of formation of the Tris-associated species remain unresolved.

The formation of cyclopentenone **4** following methanol treatment further supports the presence of reactive oxidized intermediates that can undergo nucleophilic trapping. A plausible pathway involves initial oxidation of 3-hydroxyhispidin, potentially at C-3, generating a reactive carbonyl-containing intermediate. Subsequent methanol addition could promote nucleophilic pyrone ring opening, rearrangement and cyclisation ultimately yielding cyclopentenone **4** (Fig. 5). Together, these observations support a model in which H3H catalyses an NADPH-dependent oxidative transformation of 3-hydroxyhispidin, generating reactive intermediates that can undergo a variety of further chemical transformations, with the reaction environment influencing the products ultimately formed.

## Discussion

Following elucidation of the fungal bioluminescence pathway (FBP), significant progress has been made towards its application in plants and other heterologous systems (7, 8, 23, 25, 47–49). However, fundamental biochemical characterisation of the individual pathway enzymes remains limited, particularly with respect to factors governing pathway flux. The conservation of substrate inhibition across both H3H homologues suggests that this behaviour may be an intrinsic feature of H3H catalysis, constraining pathway flux at elevated hispidin concentrations. Such behaviour could potentially limit excessive consumption of cellular resources, including NADPH, while preserving hispidin for other biosynthetic pathways. Since hispidin serves as a precursor for a diverse family of fungal styrylpyrones, restricting its conversion to 3-hydroxyhispidin may help maintain metabolite availability for competing biosynthetic processes (38, 50–52). In addition to that, limiting 3-hydroxyhispidin formation may also reduce accumulation of this intrinsically unstable biosynthetic pathway intermediate. From an applied perspective, substrate inhibition may also be an important consideration for optimisation of the FBP in heterologous expression systems. Engineering H3H variants with reduced susceptibility to substrate inhibition could increase pathway throughput and enhance autonomous light production in heterologous systems, provided downstream enzymes remain capable of processing the increased metabolite load.

Previous biochemical characterisation of H3H has primarily focused on monitoring oxygen or NADPH consumption (27). Monitoring of hispidin consumption and 3-hydroxyhispidin formation by SPE-MS revealed an unexpected secondary turnover of 3-hydroxyhispidin (Fig. 4), which was subsequently corroborated using complementary luminescence and NMR analyses (Fig. 5 and Fig. S7). Continued conversion of NADPH to NADP^+^ following hispidin depletion supports the involvement of an additional NADPH-dependent reaction rather than solely spontaneous degradation (Fig. 5). This secondary activity was observed for both H3H homologues, suggesting that it represents a conserved catalytic capability. Nevertheless, kinetic analysis demonstrated a clear preference for hispidin over 3-hydroxyhispidin, with a catalytic efficiency approximately 50-fold higher for hispidin in mcH3H, (Table 2), indicating that secondary turnover remains substantially less efficient than the canonical reaction. Whether this activity contributes to pathway behaviour *in vivo* or reflects conserved catalytic flexibility remains to be established.

**Table 2.** Catalytic efficiencies of H3H homologues and LuzTr v4 determined by SPE-MS. Catalytic efficiencies (kcat/KM) were determined from direct measurement of substrate depletion by SPE-MS. Values are reported as mean ± SD. SPE-MS, solid phase extraction coupled to mass spectrometry; H3H, hispidin-3-hydroxylase; LuzTr v4, truncated *N. nambi* luciferase variant containing I63T, T99P, T192S and A199P substitutions.

| **Enzyme** | **Hispidin-3-hydroxylase** |  |
|---|---|---|
|  | nnH3H | mcH3H |
| Assay | SPE-MS | SPE-MS |
| OH-Hispidin<br>*k<sub>cat</sub>*/K<sub>M</sub> (M<sup>-1</sup> s<sup>-1</sup>) | 4.85 ± 1.79 x 10<sup>3</sup> | 4.02 ± 1.97 x 10<sup>4</sup> |
| Hispidin<br>*k<sub>cat</sub>*/K<sub>M</sub> (M<sup>-1</sup> s<sup>-1</sup>) | 3.26 ± 1.43 x 10<sup>4</sup> | 1.98 ± 0.44 x 10<sup>6</sup> |
| NADPH *k<sub>cat</sub>*/K<sub>M</sub> (M<sup>-1</sup> s<sup>-1</sup>) | 1.13 ± 0.35 x 10<sup>3</sup> | 6.89 ± 1.11 x 10<sup>4</sup> |

Consistent with previous reports (26, 30), full-length Luz remained challenging to purify and handle in an active form, necessitating the use of truncated constructs for detailed biochemical characterisation. Characterisation of purified Luz_Tr_ and Luz_Tr_ v4 revealed comparable low micromolar apparent Michaelis constants for 3-hydroxyhispidin in the luminescence assay (K_M_ = 0.57 and 2.12 µM, respectively), despite the substantially higher luminescent output and stability of Luz_Tr_ v4 (Fig. 2 and Table 1). This suggests that the improved luminescent properties of Luz_Tr_ v4 do not arise from decreased K_M_, but may instead reflect differences in catalytic turnover or quantum yield. Direct measurement of substrate depletion by SPE-MS yielded a substantially higher apparent K_M_ for Luz_Tr_ v4 (118 µM), reflecting differences in assay conditions, readout and temporal resolution, while enabling determination of an apparent k_cat_ of 1.23 s^-1^ and direct quantification of substrate turnover. Although Luz_Tr_ v4 provided a robust reporter for coupled analysis of H3H activity, reporter concentrations substantially exceeding those of H3H were required to ensure that luciferase activity did not limit the measured rate. The coupled assay therefore provides a sensitive approach for monitoring H3H activity, while direct substrate-based approaches such as SPE-MS provide a complementary method for quantitative kinetic characterisation.

The products generated during the secondary turnover by H3H further highlight the chemical complexity associated with this system. Preparative turnover and subsequent NMR and LCMS analysis identified several smaller catechol-containing products, including compound **1**, caffeic acid (**2**) and 3,4-dihydroxybenzaldehyde (**3**). The formation of caffeic acid is particularly notable because it returns material from secondary turnover to an upstream intermediate of the FBP, potentially providing a route for recycling rather than complete loss of pathway intermediates. The diversity of products observed suggests that not all downstream products are formed directly through enzyme catalysis alone. Instead, H3H may initiate an oxidative transformation of 3-hydroxyhispidin from which reactive species can undergo subsequent rearrangement, hydrolysis and/or fragmentation. The m/z 279 species observed by SPE-MS may represent a component of this downstream chemistry, although its structure remains unresolved. Notably, formation of cyclopentenone **4** following methanol treatment supports the formation of reactive oxidised species during secondary turnover and is consistent with the proposed pathway involving a C-3 carbonyl intermediate, nucleophilic interception, pyrone ring opening and subsequent rearrangement (Fig. 5). The identification of compound **1**, a metabolite previously associated with fungal secondary metabolism and non-enzymatic oxyluciferin degradation (26, 34), also suggests a connection between the FBP and broader fungal styrylpyrone metabolism (38), although its physiological relevance remains uncertain.

The luciferase from *N. nambi* has previously been shown to accept a range of 3-hydroxyhispidin analogues (14, 47) with structural modifications focusing on the catechol moiety, demonstrating considerable substrate tolerance. Substrate flexibility of H3H has received less attention (27). The secondary H3H activity identified here demonstrates that H3H can recognise and further process its own hydroxylated product. Future substrate analogue studies could therefore explore the tolerance of H3H and Luz towards modifications of the pyrone moiety, extending previous investigations on the catechol moiety. Defining the substrate scope will help elucidate the overall chemical flexibility of the FBP and its potential for future engineering efforts.

Collectively, this work establishes robust methodologies for recombinant production of fungal bioluminescence enzymes (Fig. S29). Comparison of the two H3H homologues revealed substantial differences in recombinant behaviour, with mcH3H showing improved expression and recovery without chaperone coexpression or retention of a solubility tag. The compatibility of mcH3H with *N. nambi* Luz demonstrates that enzymes from different fungal species can be functionally combined to produce visible light, highlighting the potential value of exploiting natural enzyme diversity when constructing and optimising the FBP in heterologous systems. Purification of active truncated Luz additionally provides a foundation for future biochemical and structural studies, although its continued association with membrane fractions indicates that further optimisation will be required to improve protein recovery, stability and biochemical tractability. More broadly, further biochemical and structural characterisation of individual FBP enzymes will help elucidate their functional interplay within the native pathway and identify properties suitable for engineered systems. Combining favourable properties from enzymes across fungal species with a deeper understanding of their catalytic scope will facilitate the rational development of more efficient and versatile autonomous bioluminescent systems for applications in synthetic biology, bioimaging and biomedical research.

## Materials and Methods

### Chemicals, reagents and strains

NADPH was purchased from Melford (UK) and Biomol (Germany). 3-Hydroxyhispidin was purchased from Enamine. Hispidin was purchased from BLDpharm (China). Flavin adenine dinucleotide (FAD) was obtained from Cayman Chemical (USA). Restriction enzymes, DNA ligase and miniprep and gel extraction kits were purchased from New England Biolabs. Chemically competent *Escherichia coli* cells were purchased from Agilent (USA) or prepared in-house. All other chemicals were purchased from Sigma-Aldrich (USA).

### Plasmid construction and transformation

Genes encoding for *Neonothopanus nambi* luciferase (Luz_Fl_), truncated luciferase (Luz_Tr_), hispidin-3-hydroxylase (nnH3H), and *Mycena chlorophos* H3H (mcH3H), were codon-optimised for expression in *Escherichia coli* and synthesised in pMK RQ shuttle vectors (GeneArt, Thermo Fisher Scientific). Genes were subcloned into custom pCold vectors with Kanamycin resistance using SacI/DraIII and NotI restriction sites to generate N- or C-terminally tagged constructs. N-terminal His_6_-SUMO tagged nnH3H, mcH3H, full length Luz_Fl_ and truncated Luz_Tr_ with a 3C protease cleavage site were created, whereas full-length Luz_Fl_ and truncated Luz_Tr_ v4 were also prepared with a 3C cleavable C-terminal SUMO-His_6_ tag (Fig. S1 and 2, Table. S2). All sequences were verified by sequencing (Eurofins, Germany).

Luz and mcH3H expression plasmids were transformed into *E. coli* C43(DE3) cells, whereas nnH3H constructs were transformed into *E. coli* BL21(DE3) cells carrying either pTf16 or pGro7 (Takara Bio, Japan), encoding Trigger Factor or GroEL/ES, respectively. Transformants were generated by heat shock (42 *°*C, 30 seconds) and selected on LB agar supplemented with kanamycin (50 µg mL^-1^). Chaperone plasmids were maintained by supplementation with chloramphenicol (35 µg mL^-1^).

### Recombinant Protein Expression, Purification and Storage

#### Expression and purification of nnH3H

A single colony was used to inoculate a starter culture of Terrific Broth supplemented with 60 mM KCl, kanamycin and chloramphenicol, and the culture was grown overnight at 37 *°*C. A 6 L culture of Terrific Broth supplemented with 60 mM KCl and antibiotic was then inoculated and grown at 37 *°*C, 180 rpm, until an OD_600_ of 0.4, at which point arabinose was added to a final concentration of 0.1 mg mL^-1^. After approximately 20 min when the OD_600_ reached >0.6, the culture was cooled to 15 *°*C and induced with 0.1 mM IPTG. Cells were harvested after overnight expression by centrifugation at 4800 × g for 15 min at 4 *°*C. The cell pellets were resuspended in lysis buffer (100 mM potassium phosphate, 400 mM NaCl, 300 mM L-Arginine, pH 8.0 supplemented with phenylmethylsulfonyl fluoride (PMSF, 0.2 mM PMSF), lysozyme (1.0 mg mL^-1^) and DNase (20 μg mL^-1^) and lysed by three passes through a high-pressure cell disruptor at 25 kPSI at 4 °C (Continuous flow cell disruptor, Constant Systems, UK). Insoluble debris was removed by centrifugation at 18000 x g for 20 min at 4 *°*C. The clarified lysate was loaded onto a 5 mL HisTrap HP column (Cytiva, UK) equilibrated in buffer A (50 mM potassium phosphate, 300 mM NaCl, 25 mM imidazole, 150 mM L-arginine, 100 mM KCl, 5% (w/v) glycerol, pH 8). The column was washed with 15 column volumes (CV) buffer A, followed by 10 CV 5% buffer B (50 mM potassium phosphate, 300 mM NaCl, 1 M imidazole, 150 mM L-arginine, 100 mM KCl, 5% (w/v) glycerol, pH 8), a high salt wash (50 mM potassium phosphate, 2 M NaCl, 25 mM imidazole, 150 mM L-arginine, 100 mM KCl, 5% (w/v) glycerol, pH 8) and 8 CV 10% buffer B. The protein was eluted using a linear gradient from 100 mM to 1 M imidazole. Fractions containing nnH3H were pooled and concentrated for either 3C cleavage and dialysis overnight (50 mM potassium phosphate, 300 mM NaCl, 150 mM L-arginine, 100 mM KCl, 5% (w/v) glycerol, pH 8) followed by RIMAC and SEC, or purified as the intact SUMO fusion protein by SEC on the same day using a Superdex 200 16/600 column equilibrated in 20 mM HEPES, 150 mM NaCl, pH 8. Final protein was concentrated, aliquoted and stored at -80 *°*C.

#### Expression and purification of mcH3H

A single colony was used to inoculate a starter culture of Terrific Broth supplemented with kanamycin (50 µg mL^-1^) then grown overnight at 37°C at 180 rpm. An 18 L culture of Terrific Broth was then inoculated and grown at 37 *°*C at 180 rpm until an OD_600_ of 0.8 was reached. The cultures were cooled to 15 *°*C to induced expression. Cells were expressed overnight and harvested by centrifugation at 4800 x g for 15 min at 4 *°*C. Cell pellets were resuspended in buffer A (100 mM potassium phosphate, 400 mM NaCl, 300 mM L-arginine, 200 mM KCl, 4 mM TCEP, pH 7.4, supplemented with PMSF, lysozyme and DNase) and lysed by three passes through a high-pressure cell disruptor at 25 kPSI at 4 *°*C (Continuous flow cell disruptor, Constant Systems, UK). Insoluble debris was removed by centrifugation at 18000 x g for 20 min at 4 *°*C. The clarified lysate was loaded onto a 5 mL HisTrap HP column (Cytiva, UK) equilibrated in buffer A (50 mM potassium phosphate, 300 mM NaCl, 25 mM imidazole, 150 mM L-arginine, 100 mM KCl, 4 mM TCEP, 5% (w/v) glycerol, pH 7.4). The column was washed with Buffer A (25 CV) then 10 CV 5% buffer B (50 mM potassium phosphate, 300 mM NaCl, 500 mM imidazole, 150 mM L-arginine, 100 mM KCl, 4 mM TCEP, 5% (w/v) glycerol, pH 7.4). Protein was eluted by a step to 10% buffer B (5 CV) then a gradient to 500 mM imidazole (15 CV). Fractions containing mcH3H were pooled and concentrated for overnight 3C protease cleavage followed by RIMAC and SEC using a Superdex 200 16/600 column equilibrated in 20 mM HEPES, 150 mM NaCl, pH 8. Final protein was concentrated, aliquoted and stored at -80 *°*C.

### Expression and purification of Luz

#### N-terminal His tag

A single colony was used to inoculate a starter culture of 2xYT supplemented with kanamycin (50 µg mL^-1^) and then grown overnight at 37 *°*C at 180 rpm. A 12 L culture of Terrific Broth supplemented with kanamycin (50 µg mL^-1^) was subsequently inoculated and grown until an OD_600_ of 0.6 was reached. The cultures were cooled to 18 *°*C and expression was induced by addition of 0.5 mM IPTG. Cells were expressed overnight and harvested by centrifugation at 4800 x g at 4 *°*C for 15 min. Cell pellets were resuspended in lysis buffer (100 mM sodium phosphate, 100 mM NaCl, pH8 supplemented with PMSF, lysozyme and DNase) and lysed by three passes through a high-pressure cell disruptor at 25 kPSI at 4 *°*C (Continuous flow cell disruptor, Constant Systems, UK). Insoluble debris was removed by centrifugation at 12000 x g for 20 min at 4 *°*C. Membrane fractions were collected by ultracentrifugation at 175,000 x g (SW 32 Ti, Optima L-90K Ultracentrifuge, Beckman Coulter, USA) for 1 hr at 4 *°*C. The membrane fraction was resuspended in solubilisation buffer containing 50 mM HEPES, 500 mM NaCl, 20% glycerol, 3.25% CHAPS (w/v), pH 8 overnight at 4 *°*C. The suspension was then diluted with the same buffer containing no detergent to reduce CHAPs content below 1% before batch binding to nickel resin for 1h at 4 *°*C. The sample was loaded into a gravity column then washed with >10 CV of Buffer A (50 mM HEPES, 200 mM NaCl, 10 mM imidazole, 5 % (w/v) glycerol, 0.86% (w/v) CHAPS, pH 8). Contaminants were removed with 10 CV 5% Buffer B (50 mM HEPES, 200 mM NaCl, 500 mM imidazole, 5 % (w/v) glycerol, 0.86% (w/v) CHAPS, pH 8), whereas Luz was eluted with 5 CV 10% buffer B and 5 CV 50% buffer B. Fractions containing full-length and truncated Luz were pooled and concentrated, then 3C protease was added and dialysed overnight in buffer A containing 0.5 mM DTT. Cleaved protein was purified by RIMAC using a 1 mL HisTrap HP column (Cytiva, UK), pooled and concentrated. The protein was further purified by SEC using a Superdex 200 16/600 column equilibrated in a DDM containing buffer (20 mM HEPES, 150 mM NaCl, 0.02% (w/v) DDM, pH 8) before concentration and storage at -80 *°*C.

#### C-terminal His tag

The protocol was as above, except that CHAPS was replaced by DDM in the solubilisation and purification buffers. Specifically, membrane solubilisation was carried out in buffer containing 1% DDM, and the nickel purification buffers contained 0.02% DDM. Resolubilised membrane detergent content was diluted by half before addition to the nickel resin.

#### Spectral Emission

Luminescence emission spectra were recorded using a CLARIOstar Plus microplate reader (BMG Labtech, Germany) in 384 well microplates (white, Clear, non-binding, Greiner Bio-one, Germany). Reactions were performed in a final volume of 50 µL containing purified luciferase (60 nM) and 3-hydroxyhispidin (200 µM). Spectra were collected between 400 and 680 nm using 2 nm increments with an integration time of 0.5 s per wavelength at ambient temperature. Instrument gain was determined automatically prior to acquisition. Spectra were analysed using MARS software (BMG Labtech, Germany).

#### Kinetics

All kinetic measurements were performed at room temperature and air-saturated oxygen conditions unless otherwise stated.

#### Luciferase Kinetics

Steady-state kinetic measurements of purified luciferase were performed in 384 well microplates (white, Clear, non-binding, Greiner Bio-one, Germany) using a CLARIOstar Plus microplate reader (BMG Labtech, Germany) equipped with a reagent injector allowing sample injection and simultaneous data acquisition. Unless otherwise stated, reactions were performed in sodium phosphate buffer (100 or 200 sodium phosphate, 500 mM Na_2_SO_4_, 0.1% DDM, pH 8.0) in a final volume of 50 µL at 25 °C. Reactions were initiated by automated injection of 3-hydroxyhispidin into purified luciferase (200 nM) at 400 µL s^-1^, and luminescence was monitored at 540 ± 50 nm using the preconfigured Lumiphore setting generated from the emission spectrum of 3-hydroxyhispidin as previously described. Initial rates of the reaction were determined from the first 0.2-0.4 s of the reaction using the linear region of the progress curve and analysed using MARS software (BMG Labtech, Germany). Michaelis-Menten parameters were obtained by nonlinear regression in GraphPad Prism version 5.04 (GraphPad Software, USA). For pH dependency assays of the enzyme, the reaction was performed in 50 mM Tris-HCl, 150 mM NaCl at varying pH.

### H3H kinetics

#### Coupled Luciferase Assay

The reaction between H3H and hispidin was monitored using a coupled assay with excess Luz_tr_ v4 to ensure rapid conversion of 3-hydroxyhispidin. Reactions were performed in 384 well microplates (white, Clear, non-binding, Greiner Bio-one, Germany) in either 100 mM sodium phosphate, 500 mM Na_2_SO_4_, 0.1% DDM, pH 8.0 or 50 mM Tris-HCl, 100 mM NaCl, pH 8.0 using a BMG CLARIOStar Plus microplate reader (BMG Labtech, Germany) equipped with a reagent injector with simultaneous substrate injection and data acquisition capabilities. Luminescence output was monitored at 540 ± 50 nm. Initial assay development was performed using H3H (200 nM), Luz_Tr_ v4 (2 µM), FAD (2 µM) and NADPH (500 µM). For subsequent kinetic measurements, H3H and reporter concentrations were adjusted to ensure that luciferase activity remained in excess and did not limit the measured H3H rate, as described in the corresponding figure legends. Reactions were initiated by automated injection at 400 µL s^-1^ in a final volume of 50 µL at 25 °C. Initial rates were determined from the linear region of the progress curves during the first 3 s after substrate addition using MARS software (BMG Labtech, Germany). Steady-state kinetic parameters were obtained by nonlinear regression in GraphPad Prism version 5.04 (GraphPad Software, USA). For pH dependence experiments, the assay was performed in 50 mM Tris HCl, 150 mM NaCl over the indicated pH range.

#### Solid-phase extraction mass spectrometry

Direct monitoring of hispidin, 3-hydroxyhispidin and other cofactors was achieved using an Agilent RapidFire RF365 high throughput sampling robot connected to an Agilent 6550 accurate mass iFunnel quadrupole time of flight (QTOF) mass spectrometer. Assays were performed in a deep-well 2 mL 96 well plate (Greiner Bio-one, Germany) in 50 mM Tris-HCl, 150 mM NaCl, pH 8.0. For extended turnover experiments, reactions were typically initiated by addition of 10 µL enzyme solution containing H3H (50 µM) and FAD (100 µM) to 990 µL assay mixture, giving final concentrations of 0.5 µM H3H and 1 µM FAD. For kinetic measurements, H3H, FAD, NADPH and substrate concentrations were varied as specified in the corresponding figure legends. Reactions were rapidly mixed by pipetting before analysis. Samples were taken at the indicated time points and loaded under vacuum onto a C18 or Graphitic SPE cartridge at a flow rate of 1.5 mL min^-1^. The SPE cartridge was then washed for 5.5 sec with LCMS water containing 0.1% (v/v) formic acid at a flow rate of 1.5 mL min^-1^ to remove non-volatile buffer salts. Analytes were then eluted from the cartridge in an organic solvent (85% (v/v) acetonitrile, 15% (v/v) LCMS water containing 0.1% (v/v) formic acid) at a flow rate of 1.25 mL min^-1^ for 5.5 sec. The mass spectrometer was operated in the positive ion mode with a drying gas temperature (280 °C), drying gas flow rate (13 L min^-1^), nebulizer pressure (40 psig), sheath gas temperature (350 °C), sheath gas flow rate (12 L min^-1^), capillary voltage (4000 V), nozzle voltage (1000 V).

#### Preparative-scale H3H reaction

Preparative scale turnover reactions were performed using purified mcH3H in 50 mM NH_4_HCO_3_ (pH 8.0) prepared in D_2_O or H_2_O. Reaction mixtures contained 3-hydroxyhispidin (2.3 mg at 2.5 mM final concentration), NADPH (3 eq., 7.5 mM final concentration), FAD (0.1 eq., 0.25 mM final concentration), and mcH3H (10 µM). Reactions were incubated at room temperature and periodically oxygenated by bubbling oxygen through the reaction mixture every 20 min to maintain aerobic turnover. Reaction progress was monitored by ^1^H NMR or luminescence-based assays until complete depletion of 3-hydroxyhispidin was observed (typically after approximately 2 h). Following complete turnover, four reaction volumes of acetonitrile were added to quench the reaction and precipitate protein. Samples were centrifuged to remove precipitated protein, and the supernatant was concentrated before purification by semi-preparative HPLC. Fractions containing products of interest were identified by LCMS, pooled, concentrated and characterised by ^1^H NMR spectroscopy, 2D NMR experiments and LCMS.

LCMS analysis was conducted using a Xevo G2-XS mass spectrometer (Waters) coupled to an Acquity-UPLC system (Waters), equipped with BEH C18 column (1.7 µM), samples were loaded onto the column then eluted using a water/acetonitrile gradient of 99/1% - 5/95% (v/v) with 0.1% (v/v) formic acid. Accurate mass analysis by loop injection MS was performed on an ACQUITY I-Class PLUS UPLC System (Waters, Milford, MA, USA) coupled to an ACQUITY RDa mass spectrometer (Waters, Milford, MA, USA) equipped with an ESI probe, in positive/negative ion mode using a 50% methanol and 50% H_2_O + 0.1% formic acid (v/v) eluent.

#### ^18^O_2_ and H_2_^18^O labelling experiments

Isotope-labelling experiments were performed under controlled ^16^O_2_ or ^18^O_2_ atmospheres using a Schlenk-line setup as previously described (43–45). Reaction components were transferred into an anaerobic chamber (Belle Technology, <2 ppm O_2_) and equilibrated to remove residual O_2_ prior to preparation of the reaction mixtures. Reactions contained mcH3H (0.5 µM) and either hispidin (50 µM) with NADPH (50 or 150 µM), or 3-hydroxyhispidin (50 µM) with NADPH (100 µM), in 50 mM Tris, 150 mM NaCl, pH 8.0. Prepared samples were transferred to the sealed Schlenk line apparatus, removed from the anaerobic chamber and the system was repeatedly evacuated and purged with Ar before exposure to ^18^O_2_ [Merck, 97 atom % ^18^O]. Reactions were incubated under ^18^O_2_ for 2 h, the sealed apparatus returned to the anaerobic chamber, before quenching with formic acid to a final concentration of 0.1% (v/v) and analysis by LCMS. Corresponding reactions performed under ^16^O_2_ from air were used as unlabelled controls. For H ^18^O labelling experiments, mcH3H was buffer exchanged using Zeba™ Spin Desalting Columns (ThermoFisher, UK) and reaction mixtures were prepared under otherwise equivalent conditions using H ^18^O [Rotem, > 98 atom % ^18^O] as solvent. Reactions contained either hispidin (50 µM) with NADPH (50 or 150 µM), or 3-hydroxyhispidin (50 µM) with NADPH (100 µM), and mcH3H (0.5 µM). Reactions were incubated for 2 h before quenching with 0.1% (v/v) formic acid and analysis by LCMS. Isotope incorporation was assessed from the corresponding +2 Da mass shifts relative to reactions performed with ^16^O_2_ or H ^16^O.

#### NMR

Purified mcH3H or SUMO nnH3H was buffer exchanged into NMR buffer (25 mM Tris-d_11_ in H_2_O, pH 8.0) using Micro Bio Spin columns (Bio Rad, USA) and if needed concentrated using Amicon Ultra centrifugal filters (10 kDa MWCO, Merck, Germany). Stock solutions of hispidin and 3-hydroxyhispidin (50 mM) were prepared in DMSO-d_6_. All other cofactor solution including FAD (1 mM) and NADPH (100 mM) were prepared in H_2_O. Reaction mixtures typically contained 250 µM hispidin or 3-hydroxyhispidin, 10 to 50 µM FAD, 1 to 3 equivalents of NADPH relative to substrate, and 0.5 to 3 µM H3H in NMR buffer containing 10% D_2_O (v/v). Exact enzyme, cofactor and substrate concentrations varied between experiments and are specified in the corresponding figure legends. Reaction mixtures were prepared immediately prior to analysis, transferred to 3 mm NMR tubes (Norell, USA), and monitored at 298 K. Unless otherwise stated, reactions were performed under ambient oxygen, with periodic oxygenation for preparative scale reactions.

^1^H NMR spectra were acquired on a Bruker AVIII 700 MHz spectrometer equipped with a 5 mm inverse TCI cryoprobe. Time course experiments were recorded using the PROJECT Carr Purcell Meiboom Gill (PROJECT CPMG) pulse sequence with water presaturation (53). Typical acquisition parameters were a total echo time of 40 ms and a relaxation delay of 2 s. For structural characterisation of isolated compounds, standard one-dimensional ^1^H NMR experiments and two-dimensional ^1^H,^1^H-COSY, ^1^H,^13^C-HSQC and ^1^H,^13^C-HMBC experiments were acquired. Spectra were processed using TopSpin 3.6.1 (Bruker, Germany) and MestReNova 14.1 (Mestrelab, Spain).

#### Organic Synthesis

##### Synthesis of (3*E*)-4-(3,4-dihydroxyphenyl)-3-buten-2-one

The synthetic procedure was adapted from the literature (14). To a solution of 3,4-(methylenedioxy)benzylideneacetone (100 mg, 0.526 mmol, 1.0 equiv.) in dry CH_2_Cl_2_ (4 mL), 1 M BBr_3_ in CH_2_Cl_2_ (2.6 mL, 2.63 mmol, 5.0 equiv.) was added at room temperature under Argon atmosphere. The reaction mixture was stirred overnight at room temperature. The reaction was quenched with 100 mM sodium phosphate buffer (pH 7.0, 20 mL) at 0 °C. The layers were separated and the aqueous layer was extracted with EtOAc (2 × 20 mL). The organic layers were pooled, dried over MgSO_4_, filtered, then concentrated *in vacuo*. The crude material was purified by reversed-phase flash column chromatography (5–100% acetonitrile in H_2_O (+ 0.1% formic acid) over 25 CV) to yield the final product as a yellow solid (24 mg, 26% yield).^1^H NMR (500 MHz, MeOD) δ 7.52 (d, *J* = 16.1 Hz, ^1^H), 7.08 (d, *J* = 2.1 Hz, 1H), 6.99 (dd, *J* = 8.2, 2.1 Hz, 1H), 6.79 (d, *J* = 8.2 Hz, 1H), 6.55 (d, *J* = 16.1 Hz, 1H), 2.33 (s, 3H). ^13^C NMR (126 MHz, D_2_O) δ 204.4, 147.4, 146.7, 144.3, 126.9, 124.1, 123.2, 116.2, 115.2, 48.8. HRMS (ESI): *m/z* [M + H]^+^ calculated for C_10_H_11_O_3_: 179.0708, found: 179.0701.

## Supporting information

This article contains supporting information.

## Conflicts of interest

The authors declare that they have no conflicts of interest with the contents of this article.

## Acknowledgments

This research was funded by the Wellcome Trust (grant no.: 227298/Z/23/Z) and the John fell Fund Oxford. P.R. thanks C.J.S. for continuous support and use of instruments. We thank A. Oluwole and the Robinson Group (Oxford) for ultracentrifuge access.

## Author contributions

J.F.B., P.R. conceptualization; J.F.B., A.T., J.L., H.W,M., P.R. methodology; J.F.B., J.L., A.T., H.W.M, P.R. validation; J.F.B., I.P., I.X.R.C., C.W., A.T., J.L., P.R. investigation; J.F.B. writing-original draft; J.F.B., J.L.,

P.R. writing-review & editing; J.F.B., J.L., P.R. visualisation; P.R. supervision; P.R. funding acquisition.

## Funding and additional information

P.R. thanks the Wellcome Trust (grant no.: 227298/Z/23/Z) for funding this research. J.L. acknowledges funding from the Deutsche Forschungsgemeinschaft (Walter Benjamin Fellowship, project number: 564012687).

## Abbreviations

nnH3H,: hispidin 3-hyroxylase from *Neonothopanus nambi*;
mcH3H: hispidin 3-hyroxylase from *Mycena chlorophos*;
Luz,: Luciferase from *Neonothopanus nambi*;
LuzTr,: (38-267) truncated luciferase from *Neonothopanus nambi*;
LuzTr v4,: truncated Luciferase from *Neonothopanus nambi* with I63T, T99P, T192S and A199P mutations;
SPE-MS,: solid phase extraction coupled to mass spectrometry;
CPH,: caffeylpyruvate hydrolase;
HispS,: hispidin synthase.

## Notes

### Competing Interest Statement

The authors have declared no competing interest.

### Summary of Updates

The manuscript has been substantially revised and expanded. Additional experiments validate the coupled H3H-luciferase assay and further assess the rapid luciferase measurements. Key kinetic measurements were repeated and reanalysed, including direct SPE-MS analysis of H3H activity. Additional NMR controls were performed to distinguish H3H-dependent secondary turnover of 3-hydroxyhispidin from non-enzymatic degradation. The isotope-labelling experiments and proposed mechanism of secondary turnover have been revised and interpreted more cautiously. Figures, Supporting Information (SI), experimental methods, and figure cross-references have also been updated throughout.

