## Supplementary Information for "Biochemical characterisation of fungal bioluminescence enzymes reveals substrate inhibition and secondary turnover of 3-hydroxyhispidin by H3H"

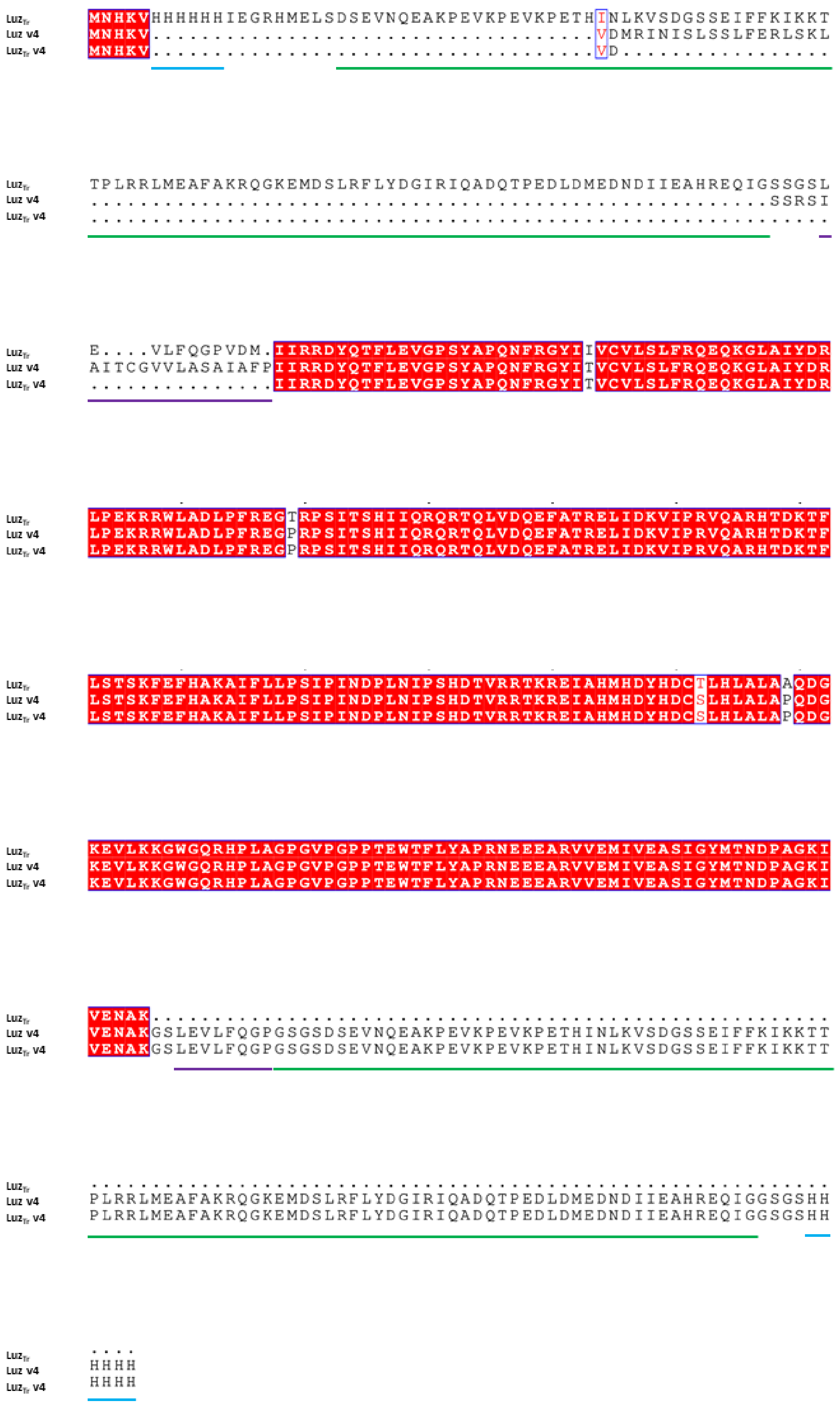


**Figure S1: Amino acid sequences of luciferase constructs used in this study.** $Luz_{Tr}$, Luz v4 and $Luz_{Tr}$ v4 construct alignments were performed using ESPript 3.2. (55), $Luz_{Tr}$ is N-terminally His-SUMO tagged, Luz v4 and $Luz_{Tr}$ v4 constructs are C-terminally SUMO-His tagged. All constructs contain a 3C cleavage sequence (purple) for tag removal. Conserved residues are highlighted in red. SUMO tag (green) and His tags (blue).

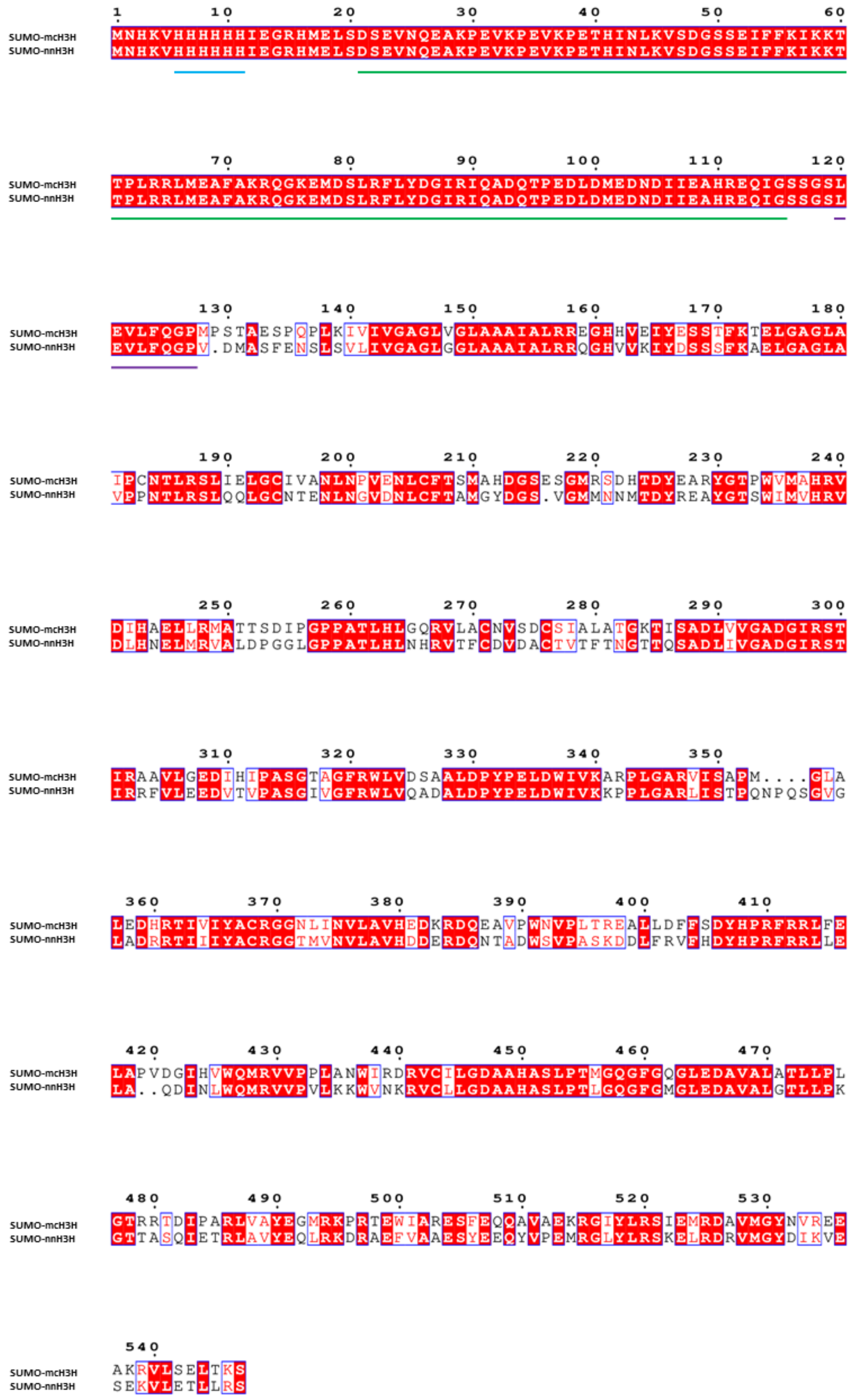


**Figure S2: Amino acid sequences of Hispidin-3-hydroxylase constructs used in this study**. mcH3H and nnH3H construct alignments were performed using ESPript 3.2. (55), both constructs contain an N-terminal His-SUMO tag and a 3C cleavage sequence for tag removal (purple). Conserved residues are highlighted in red. SUMO tag (green) and His tags (blue).

**Expression Construct Sequences**

**SUMO nnH3H**

MNHKVHHHHHHIEGRHMELSDSEVNQEAKPEVKPEVKPETHINLKVSDGSSEIFFKIKKTTPLRRLMEAFAKRQGK
EMDSLRFLYDGIRIQADQTPEDLDMEDNDIIEAHREQIGSSGSLEVLFQGPVDMASFENSLSVLIVGAGLGGLAAAI
ALRRQGHVVKIYDSSSFKAELGAGLAVPPNTLRSLQQLGCNTENLNGVDNLCFTAMGYDGSVGMMNNMTDYRE
AYGTSWIMVHRVDLHNELMRVALDPGGLGPPATLHLNHRVTFCDVDACTVTFTNGTTQSADLIVGADGIRSTIRR
FVLEEDVTVPASGIVGFRWLVQADALDPYPELDWIVKKPPLGARLISTPQNPQSGVGLADRRTIIIYACRGGTMVN
VLAVHDDERDQNTADWSVPASKDDLFRVFHDYHPRFRRLLELAQDINLWQMRVVPVLKKWVNKRVCLLGDAAH
ASLPTLGQGFGMGLEDAVALGTLLPKGTTASQIETRLAVYEQLRKDRAEFVAAESYEEQYVPEMRGLYLRSKELRDR
VMGYDIKVESEKVLETLLRSSNSA*

**SUMO mcH3H**

MNHKVHHHHHHIEGRHMELSDSEVNQEAKPEVKPEVKPETHINLKVSDGSSEIFFKIKKTTPLRRLMEAFAKRQGK
EMDSLRFLYDGIRIQADQTPEDLDMEDNDIIEAHREQIGSSGSLEVLFQGPMPSTAESPQPLKIVIVGAGLVGLAAAI
ALRREGHHVEIYESSTFKTELGAGLAIPCNTLRSLIELGCIVANLNPVENLCFTSMAHDGSESGMRSDHTDYEARYGT
PWVMAHRVDIHAELLRMATTSDIPGPPATLHLGQRVLACNVSDCSIALATGKTISADLVVGADGIRSTIRAAVLGE
DIHIPASGTAGFRWLVDSAALDPYPELDWIVKARPLGARVISAPMGLALEDHRTIVIYACRGGNLINVLAVHEDKRD
QEAVPWNVPLTREALLDFFSDYHPRFRRLFELAPVDGIHVWQMRVVPPLANWIRDRVCILGDAAHASLPTMGQG
FGQGLEDAVALATLLPLGTRRTDIPARLVAYEGMRKPRTEWIARESFEQQAVAEKRGIYLRSIEMRDAVMGYNVRE
EAKRVLSELTKSDC*

**N-terminal SUMO (38-267) Luz$_{Tr}$**

MNHKVHHHHHHIEGRHMELSDSEVNQEAKPEVKPEVKPETHINLKVSDGSSEIFFKIKKTTPLRRLMEAFAKRQGK
EMDSLRFLYDGIRIQADQTPEDLDMEDNDIIEAHREQIGSSGSLEVLFQGPVDMIIRRDYQTFLEVGPSYAPQNFRG
YIIVCVLSLFRQEQKGLAIYDRLPEKRRWLADLPFREGTRPSITSHIIQRQRTQLVDQEFATRELIDKVIPRVQARHTDK
TFLSTSKFEFHAKAIFLLPSIPINDPLNIPSHDTVRRTKREIAHMHDYHDCTLHLALAAQDGKEVLKKGWGQRHPLA
GPGVPGPPTEWTFLYAPRNEEEARVVEMIVEASIGYMTNDPAGKIVENAK

**C-terminal SUMO (38-267) Luz$_{Tr}$ v4**

MNHKVVDIIRRDYQTFLEVGPSYAPQNFRGYITVCVLSLFRQEQKGLAIYDRLPEKRRWLADLPFREGPRPSITSHII
QRQRTQLVDQEFATRELIDKVIPRVQARHTDKTFLSTSKFEFHAKAIFLLPSIPINDPLNIPSHDTVRRTKREIAHMHD
YHDCSLHLALAPQDGKEVLKKGWGQRHPLAGPGVPGPPTEWTFLYAPRNEEEARVVEMIVEASIGYMTNDPAGK
IVENAKGSLEVLFQGPGSGSDSEVNQEAKPEVKPEVKPETHINLKVSDGSSEIFFKIKKTTPLRRLMEAFAKRQGKEM
DSLRFLYDGIRIQADQTPEDLDMEDNDIIEAHREQIGGSGSHHHHHH*

**C-terminal SUMO Luz v4**

MNHKVVDMRINISLSSLFERLSKLSSRSIAITCGVVLASAIAFPIIRRDYQTFLEVGPSYAPQNFRGYITVCVLSLFRQE
QKGLAIYDRLPEKRRWLADLPFREGPRPSITSHIIQRQRTQLVDQEFATRELIDKVIPRVQARHTDKTFLSTSKFEFHA
KAIFLLPSIPINDPLNIPSHDTVRRTKREIAHMHDYHDCSLHLALAPQDGKEVLKKGWGQRHPLAGPGVPGPPTEW
TFLYAPRNEEEARVVEMIVEASIGYMTNDPAGKIVENAKGSLEVLFQGPGSGSDSEVNQEAKPEVKPEVKPETHINL
KVSDGSSEIFFKIKKTTPLRRLMEAFAKRQGKEMDSLRFLYDGIRIQADQTPEDLDMEDNDIIEAHREQIGGSGSHH
HHHH*

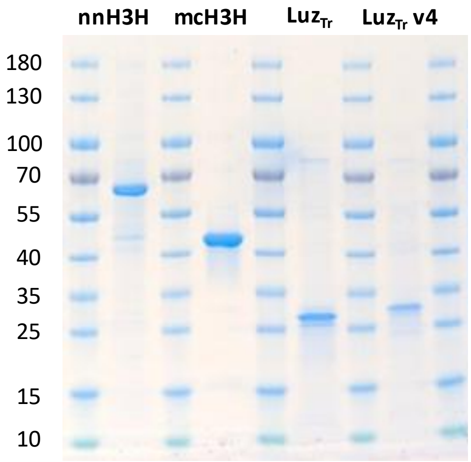


**Figure S3: SDS-PAGE gel analysis of purified proteins**. Coomassie stained SDS PAGE gel of the final purified recombinant proteins used in this study. Molecular masses of the protein standards are indicated in kDa. The expected molecular masses of the purified proteins are 61.3 kDa for His$_6$-SUMO-nnH3H, 46.3 kDa for mcH3H, 26.9 kDa for Luz$_{Tr}$, and 28.1 kDa for Luz$_{Tr}$ v4. Luz$_{Tr}$, residues 38 - 267 of luciferase; nnH3H, *Neonothopanus nambi* hispidin-3-hydroxylase; mcH3H, *Mycena chlorophos* hispidin-3-hydroxylase.

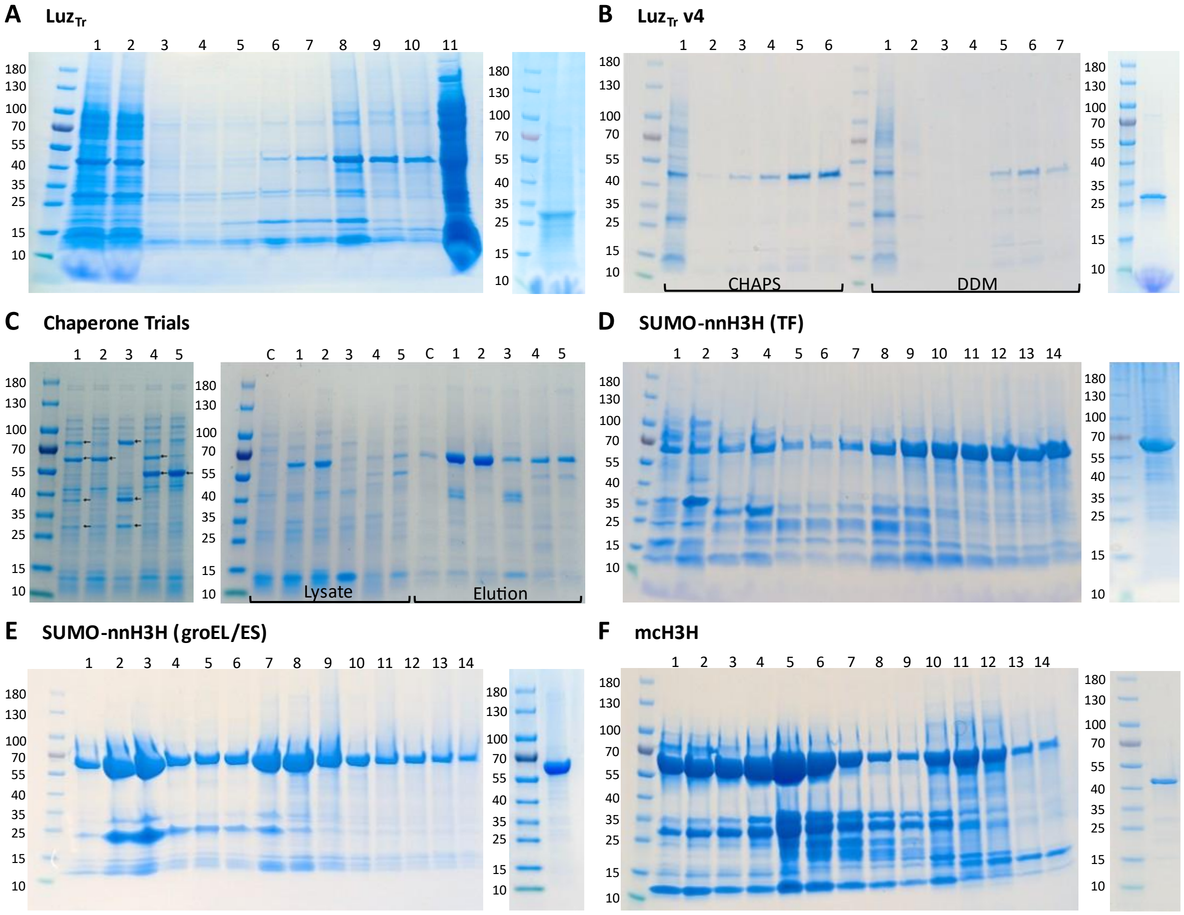


**Figure S4: SDS-PAGE gel analysis of SUMO nnH3H, mcH3H and Luz IMAC and SEC purifications.** Coomassie stained SDS PAGE analysis of recombinant Luz and H3H expression and purification. (**A**) Gravity column purification of N-terminally His$_6$ SUMO tagged Luz$_{Tr}$ (41.4 kDa) using CHAPS for membrane solubilisation (left) followed by 3C protease cleavage, reverse IMAC and SEC yielding purified Luz$_{Tr}$ (26.9 kDa, right). Left: ladder, 1/2. Flowthrough, 3/4. Buffer A wash, 5-7. 10% bufferB elution, 8/9. 50% buffer B elution, 10. 100% buffer B elution, 11. Soluble lysate. Right: Final purified Luz$_{Tr}$. (**B**) Gravity column purification of C-terminally SUMO Luz$_{Tr}$ v4 (40.7 kDa) using CHAPS and DDM for membrane solubilisation (left) followed by 3C protease cleavage, reverse IMAC and SEC yielding purified Luz$_{Tr}$ v4 (28.1 kDa, right). Left: Ladder, 1. Flowthrough, 2. Buffer A wash, 3/4. 10% buffer B elution, 5/6. 50% buffer B elution. Right: Final purified Luz$_{Tr}$ v4. Luz purifications used a 16.600 200 pg SEC column, equilibrated in 20 mM Hepes 150 mM NaCl 0.02% (w/v) DDM pH 8. (**C**) Chaperone co-expression screen for SUMO nnH3H. Left: Expression of chaperone sets in C43 cells: 1. DnaK, DnaJ, GrpE, GroES and GroEL, 2. GroES and GroEL, 3. DnaK, DnaJ and GrpE, 4. GroES, GroEL and trigger factor, 5. Trigger factor. Right: Co-expression of SUMO nnH3H with chaperone sets 1 to 5 and a chaperone negative control with both lysate and elution fractions shown. Expressed chaperones are indicated with arrows. Note: The best expression was observed with GroES and GroEL (2) then Trigger factor (5) chaperon co-expression, however SUMO nnH3H overlaps with GroEL chaperone. Expression was performed in TB, with chaperone induction using arabinose (0.1 mg ml$^{-1}$) at OD$_{600}$ = 0.4 at 37 °C for ~20 min until OD$_{600}$ = 0.6 where cells were cold shocked and induced with IPTG (0.1 mM) then expressed O/N at 15°C. (**D**) IMAC purification of SUMO nnH3H (61.3 kDa) with trigger factor chaperone (56 kDa) co-expression followed by SEC. Right: Ladder, 1/2. 5% buffer B Wash, 3/4. High salt wash using 2 M NaCl, 5-7. 10% buffer B wash, 8-14 Gradient elution from 10% - 100% buffer B. Right: Final purified SUMO nnH3H after SEC. (**E**) IMAC purification of SUMO nnH3H (61.3 kDa) with groEL/ES chaperones (60/10 kDa) co-expression followed by SEC. Right: Ladder, 1/2. 5% buffer B Wash, 3/4. High salt wash using 2 M NaCl, 5-7. 10% buffer B wash, 8-14 Gradient elution from 10% - 100% buffer

B. Right: Final purified SUMO nnH3H after SEC. nnH3H purifications used a 16.600 200 pg SEC column, equilibrated in 20 mM Hepes 150 mM NaCl pH 8. (**F**) IMAC purification of SUMO mcH3H (60.8 kDa) followed by 3C protease cleavage, reverse IMAC and SEC yielding mcH3H (46.3 kDa). Right: ladder, 1-4. 5% buffer B elution, 5-9. 10% buffer B elution, 10-14. gradient elution from 10% - 100% buffer B. Left: Final purified SEC product after 3C cleavage and reverse IMAC using a 26.600 200 pg SEC column equilibrated in 20 mM Hepes 150 mM NaCl pH 8. $Luz_{Tr}$, residues 38 - 267 of *Neonothopanus nambi* luciferase; $Luz_{Tr}$ v4, engineered LuzTr variant containing I63T, T99P, T192S and A199P substitutions; nnH3H, *N. nambi* hispidin-3-hydroxylase; mcH3H, *Mycena chlorophos* hispidin-3-hydroxylase; IMAC, immobilised metal affinity chromatography; SEC, size exclusion chromatography.

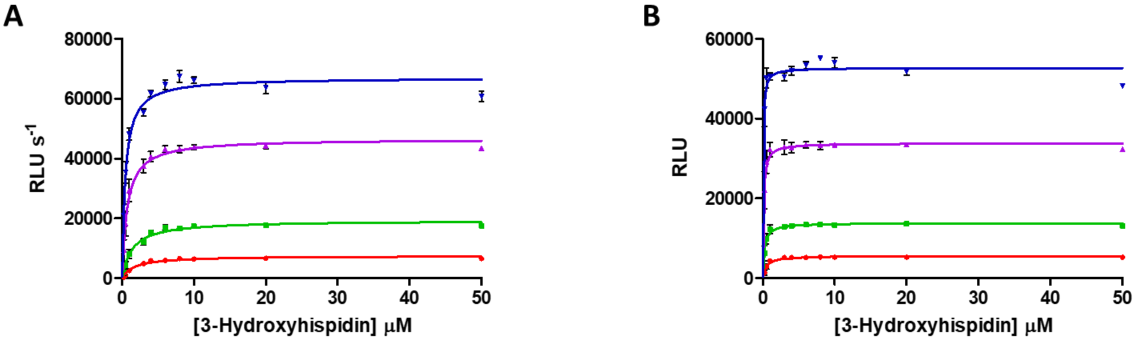


C

| $Luz_{Tr}$ v4 (nM) | $K_M$ (µM) | |
|---|---|---|
| | Initial Rate | Maxima |
| 200 | 0.453 ± 0.064 | 0.050 ± 0.015 |
| 100 | 0.714 ± 0.076 | 0.108 ± 0.015 |
| 40 | 1.43 ± 0.210 | 0.230 ± 0.030 |
| 20 | 1.71 ± 0.345 | 0.461 ± 0.084 |

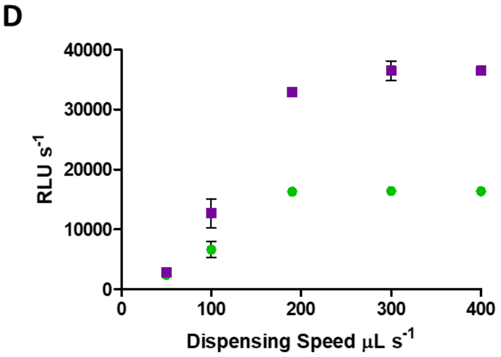


**Figure S5: $Luz_{Tr}$ v4 rate analysis and dispensing controls.** (**A-C**) $Luz_{Tr}$ v4 (blue: 200, purple: 100, green: 40, red: 20 nM) and 3-hydroxyhispidin (50 - 0.25 µM), Michaelis-Menten kinetic fit from (A) the initial rate of RLU change (ΔRLU $s^{-1}$) t = 0 – 0.45 s and (**B**) maximal RLU during plateau determined as an average between t=4 and t=10 s. (**C**) $K_M$ parameters from analysis methods **A**/**B**. (**D**) Dispensing controls for initial rate analysis. Reaction of $Luz_{Tr}$ v4 (Purple: 100, Green: 50 nM) and 3-hydroxyhispidin (Purple: 10, Green 50 µM) initiated by substrate addition at varying speeds (50 – 400 µL $s^{-1}$). Luminescence-based plate reader assays were performed in sodium phosphate buffer (100 mM $NaH_2PO_4$/$Na_2HPO_4$, 500 mM $Na_2SO_4$, 0.1% DDM, pH 8.0). Results were collected in triplicate (n=3; mean ± SD). $Luz_{Tr}$, (38-267) luciferase; RLU, relative luminescent units.

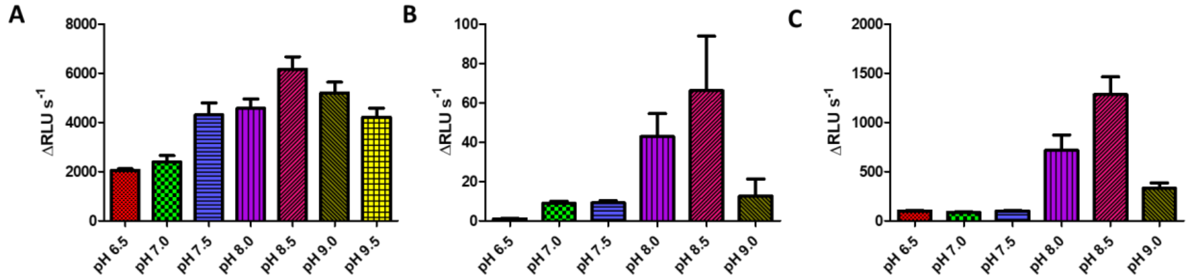


**Figure S6: pH dependency on the activity of purified FBP enzymes Luz$_{Tr}$ v4, SUMO nnH3H and mcH3H.** (**A**) The initial rate of reaction between Luz$_{Tr}$ v4 (60 nM) and 3-hydroxyhispidin (50 µM) was determined from the change in luminescence over time in sodium phosphate buffer (200 mM $NaH_2PO_4$/$Na_2HPO_4$, 500 mM $Na_2SO_4$, 0.1% DDM). (**B**) The initial rate of the reaction between SUMO nnH3H (200 nM) and hispidin (5 µM) with FAD (2 µM) and NADPH (250 µM) was determined using a coupled luminescence-based assay with Luz$_{Tr}$ v4 (2 µM) in Tris buffer (50 mM Tris, 100 mM NaCl). (**C**) The initial rate of reaction between mcH3H (20 nM) and hispidin (3.2 µM) with NADPH (250 µM) was determined from the change in luminescence over time via a coupled luminescence-based assay with Luz$_{Tr}$ v4 (200 nM) in Tris buffer (50 mM Tris 100 mM NaCl). Results were collected in triplicate (n=3; mean ± SD). FBP, fungal bioluminescence pathway; Luz$_{Tr}$, (38-267) luciferase; nnH3H, *Neonothopanus nambi* hispidin-3-hydroxylase; mcH3H, *Mycena chlorophos* hispidin-3-hydroxylase; RLU, relative luminescent units.

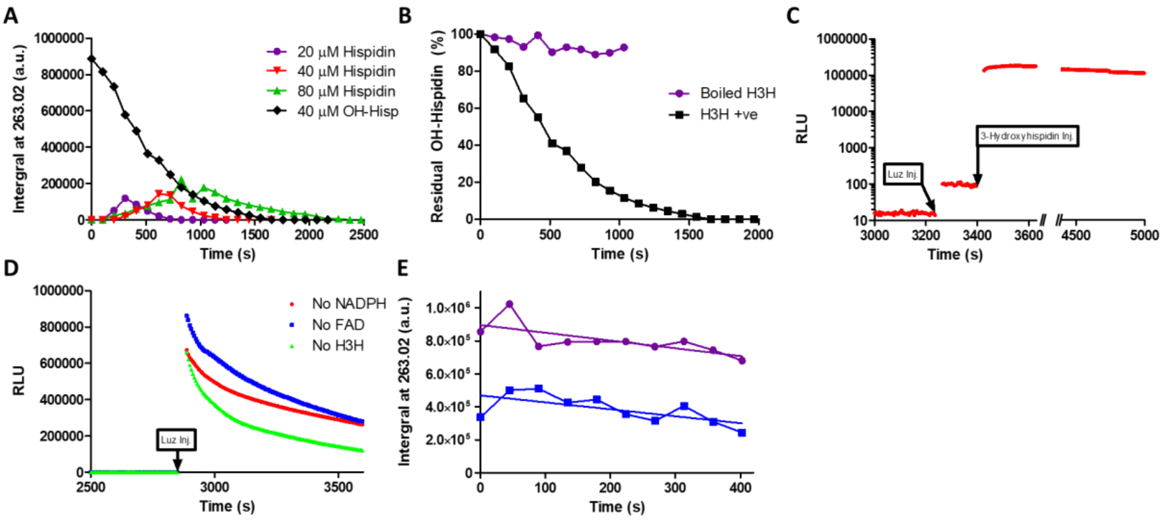


**Figure S7: Control experiments support H3H catalysed turnover of 3-hydroxyhispidin.** (**A**) SPE-MS analysis of 3-hydroxyhispidin accumulation and subsequent depletion following turnover of hispidin (20, 40 or 80 µM) or direct incubation of 3-hydroxyhispidin (OH-Hisp, 40 µM) with SUMO nnH3H (500 nM) in the presence of NADPH (2 mM) and FAD (10 µM). Ion intensity at m/z = 263.0281 ($[M+H]^+$) was used to monitor 3-hydroxyhispidin throughout the reaction. (**B**) Control experiment demonstrating that 3-hydroxyhispidin turnover requires active enzyme. 3-hydroxyhispidin (40 µM) was incubated with native or heat denatured SUMO nnH3H (500 nM) in the presence of NADPH (2 mM) and FAD (10 µM). (**C**) Orthogonal coupled luciferase assay confirming 3-hydroxyhispidin turnover following incubation with H3H. SUMO nnH3H (500 nM) was preincubated with 3-hydroxyhispidin (40 µM), NADPH (2 mM) and FAD (5 µM) for >40 min before addition of $Luz_{Tr}$ v4 (1 µM) to detect residual 3-hydroxyhispidin. Subsequent addition of fresh 3-hydroxyhispidin (40 µM) confirmed that the absence of an initial luminescence response was not due to inhibition or loss of activity of $Luz_{Tr}$ v4. (**D**) Component omission controls demonstrating that 3-hydroxyhispidin turnover requires H3H, NADPH and FAD. SUMO nnH3H (500 nM) was preincubated with 3-hydroxyhispidin (40 µM) for >40 min in the absence of NADPH (red), FAD (blue) or enzyme (green) before addition of $Luz_{Tr}$ v4 (1 µM) to detect residual 3-hydroxyhispidin. Absence of any individual component restored the luminescence response of $Luz_{Tr}$ v4, consistent with presence of 3-hydroxyhispidin (**E**) SPE-MS catalase control demonstrating that 3-hydroxyhispidin turnover is not driven by hydrogen peroxide generated during reaction. Consumption of 3-hydroxyhispidin (50 µM) by SUMO nnH3H (200 nM) was monitored by SPE-MS in the presence or absence of catalase (200 nM), with NADPH (100 µM) and FAD (5 µM). Ion intensity at m/z = 263.0281 ($[M+H]^+$) was used to monitor 3-hydroxyhispidin throughout the reaction. All assays were performed in 50 mM Tris-HCl, 150 mM NaCl, pH 8.0. nnH3H, *Neonothopanus nambi* hispidin-3-hydroxylase; $Luz_{Tr}$, (38-267) luciferase; RLU, relative luminescent units; SPE-MS, solid phase extraction coupled to mass spectrometry; OH-Hisp, 3-hydroxyhispidin; a.u., arbitrary units.

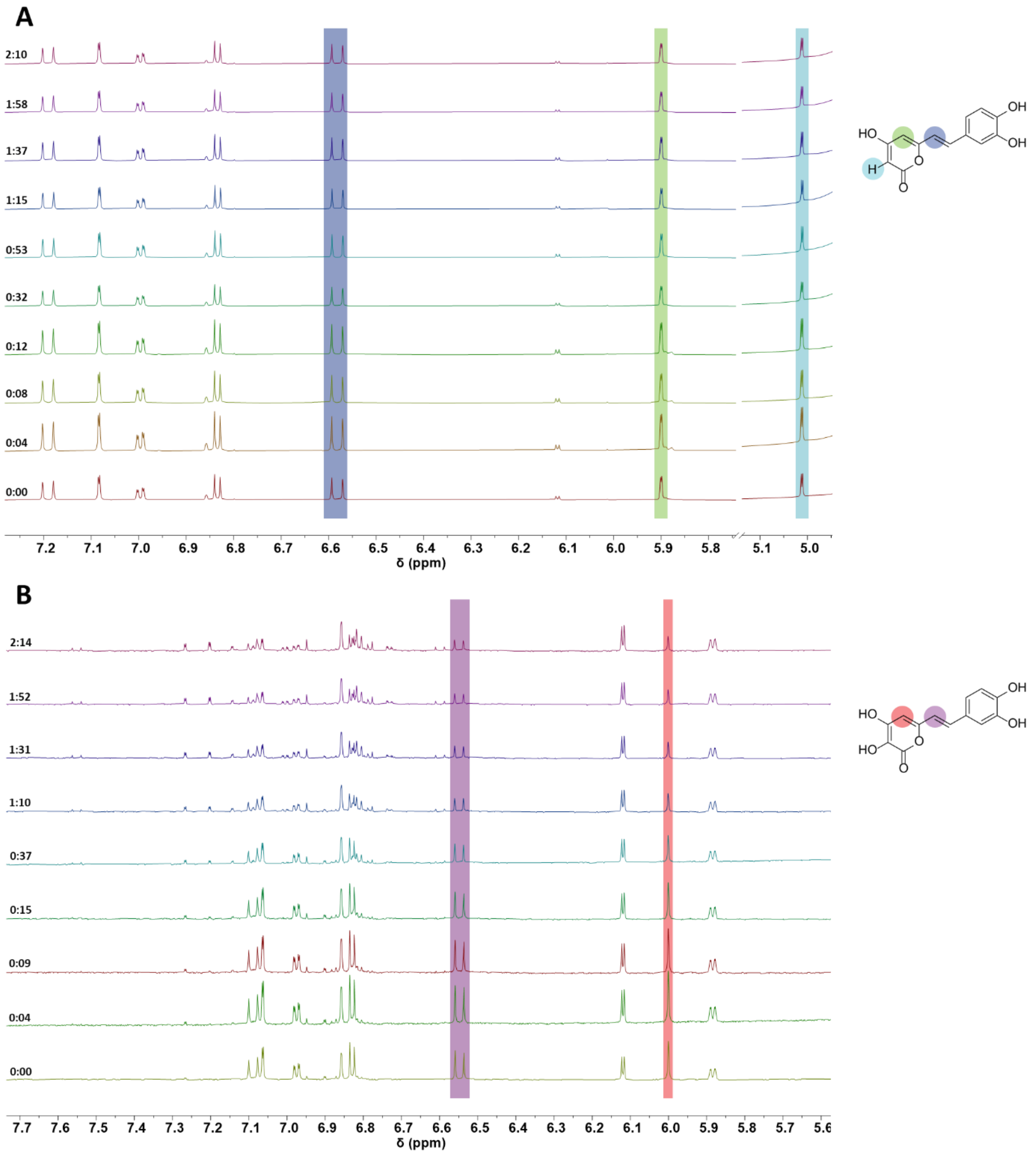


**Figure S8: Non-enzymatic stability of hispidin and 3-hydroxyhispidin under assay conditions.** Substrate stability was monitored over 3 h in the absence of enzyme. Reactions contained substrate (250 µM), NADPH (250 µM) and FAD (500 nM). (**A**) Hispidin. (**B**) 3-hydroxyhispidin. Note: hispidin remained stable over the 3 h time course, while 3-hydroxyhispidin showed minor non-enzymatic decomposition. Spectra were acquired in 25 mM Tris-$d_{11}$ (pH 8.0, 10% $D_2O$) using a Bruker AVIII 700 MHz spectrometer with water presaturation. Resonances corresponding to the substrate protons used to monitor substrate depletion are highlighted. Time shown in h:mm.

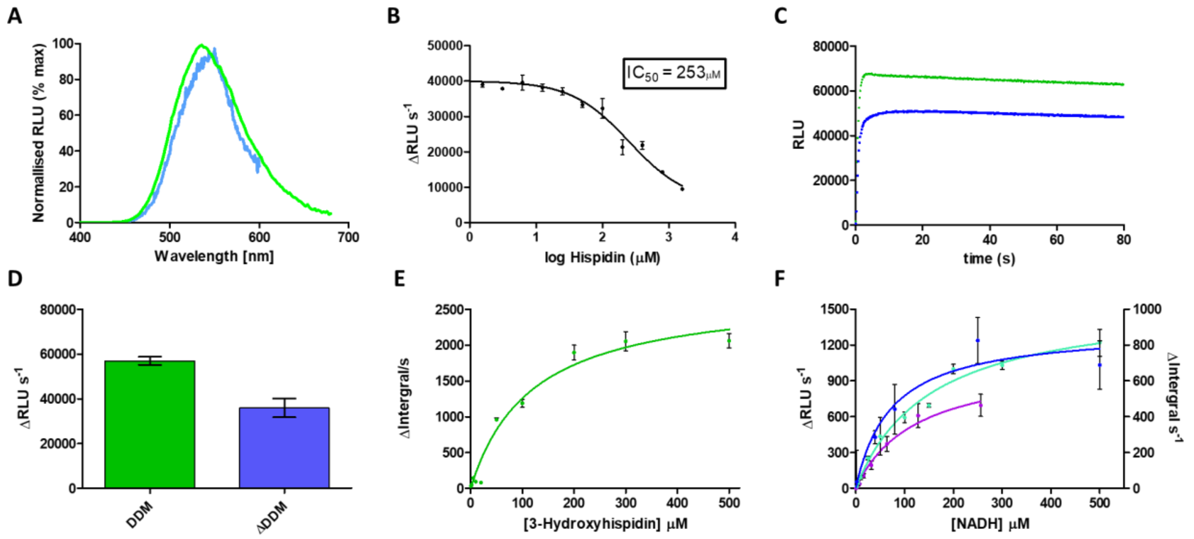


**Figure S9: Control experiments characterising luciferase and H3H assay conditions.** (**A**) Emission spectra of the reaction between 3-hydroxyhispidin with $Luz_{Tr}$ (450 – 600 nm, blue) or $Luz_{Tr}$ v4 (400 – 680 nm, green) normalised to the maximum RLU of each spectrum. (**B**) Competitive inhibition of $Luz_{Tr}$ v4 (200 nM) by Hispidin. Initial rates of 3-hydroxyhispidin (2 µM) turnover by $Luz_{Tr}$ v4 was determined from the change in RLU over time across a range of hispidin concentrations (1600, 800, 400, 200, 100, 50, 25, 12.5, 6.25, 3.125, 1.5625, 0.78125, 0 µM). Assays were performed in sodium phosphate buffer (200 mM $NaH_2PO_4$/$Na_2HPO_4$, 500 mM $Na_2SO_4$, 0.1% DDM, pH 8.0) and data were fit to three parameter inhibitor versus response model in Prism. (**C**) Progress curves for the reaction between $Luz_{Tr}$ v4 (100 nM) and 3-hydroxyhispidin (250 µM) in sodium phosphate buffer (200 mM $NaH_2PO_4$/$Na_2HPO_4$, 500 mM $Na_2SO_4$, 0.1% DDM, pH 8.0, green) or SPE-MS buffer (50 mM Tris-HCl, 150 mM NaCl, pH 8.0, blue). (**D**) Impact of detergent on luciferase activity. Initial rate analysis of progress curves shown in (C), calculated from the change in RLU over time. (**E**) Steady-state kinetic analysis of purified $Luz_{Tr}$ v4 by SPE-MS. Initial rates were determined from depletion of 3-hydroxyhispidin monitored by the ion intensity at m/z 263.0281 ($[M+H]^+$). Luminescence-based plate reader assays were performed in sodium phosphate buffer (200 mM $NaH_2PO_4$/$Na_2HPO_4$, 500 mM $Na_2SO_4$, 0.1% DDM, pH 8.0). (**F**) NADH utilisation by SUMO nnH3H and mcH3H measured using coupled luciferase and SPE-MS assays. Coupled assays contained SUMO nnH3H (200 nM; teal), LuzTr v4 (2 µM), FAD (2 µM) and hispidin (5 µM), or mcH3H (20 nM; purple), LuzTr v4 (200 nM) and hispidin (6.4 µM), with varying NADH concentrations. SPE-MS analysis contained SUMO nnH3H (200 nM; blue), hispidin (10 µM), FAD (5 µM) and varying NADH concentrations. $K_M$ kinetic constants calculated from nnH3H coupled, SPE-MS and mcH3H coupled assays at 53 ± 25.6, 73.0 ± 37.0, and 123 ± 23.6 µM respectively. Coupled luciferase and SPE-MS assays were performed in 50 mM Tris-HCl, 150 mM NaCl, pH 8.0. SPE-MS was performed on a C-18 cartridge. Results were collected in triplicate (n=3, mean ± SD), except mcH3H NADH data (n=9, mean ± SD). $Luz_{Tr}$, (38-267) luciferase; RLU, relative luminescent units; nnH3H*, Neonothopanus nambi hispidin*-3-hydroxylase; mcH3H, *Mycena chlorophos* hispidin-3-hydroxylase; SPE-MS, solid phase extraction coupled to mass spectrometry.

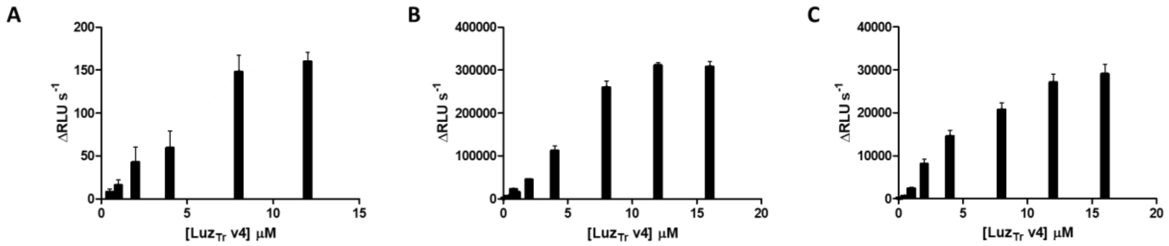


**Figure S10: H3H-Luz coupled assay validation.** Luciferase was screened against reported activity of coupled H3H protein at fixed concentrations to demonstrate successful coupling. (**A**) SUMO nnH3H (50 nM), NADPH (500 µM), hispidin (10 µM), FAD (0.5 µM). (**B**) mcH3H (20 nM), NADPH (100 µM), hispidin (6.4 µM). (**C**) mcH3H (2 nM), NADPH (500 µM), hispidin (6.4 µM). Coupled luciferase assays were performed in 50 mM Tris-HCl, 150 mM NaCl, pH 8.0. Results were collected in triplicate (n=3, mean ± SD). $Luz_{Tr}$, (38-267) luciferase; RLU, relative luminescent units; nnH3H, *Neonothopanus nambi* hispidin-3-hydroxylase; mcH3H, *Mycena chlorophos* hispidin-3-hydroxylase.

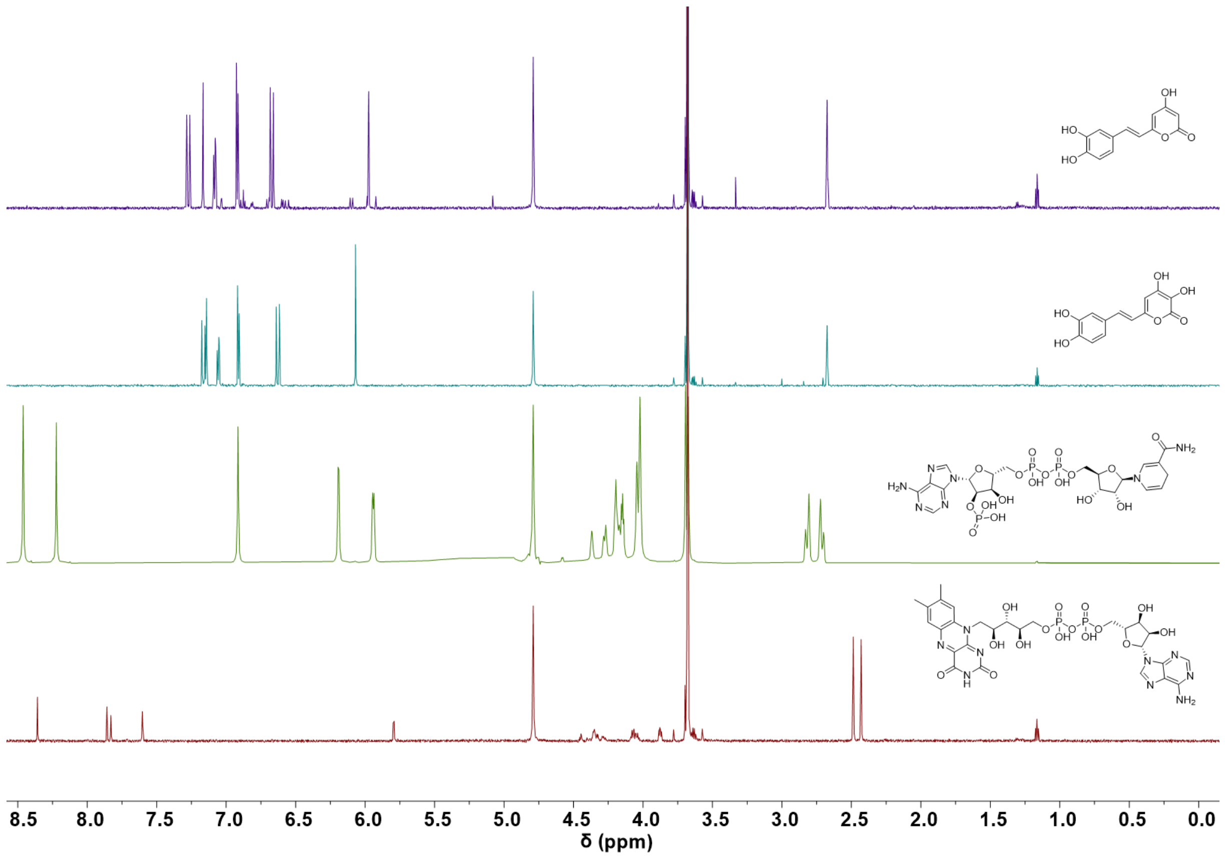


**Figure S11: $^1$H NMR spectra of fungal bioluminescence pathway substrates and cofactors**. Spectra of 50 µM FAD in $D_2O$ (red), 2 mM NADPH in $D_2O$ (green), 250 µM 3-hydroxyhispidin (turquoise) and 250 µM hispidin (purple) in 25 mM Tris-$d_{11}$ pH 7.5 containing 10% (v/v) $D_2O$. Stock solutions of hispidin and 3-hydroxyhispidin were initially dissolved in DMSO-$d_6$. Spectra were acquired using a Bruker AVIII 700 MHz spectrometer with water presaturation.

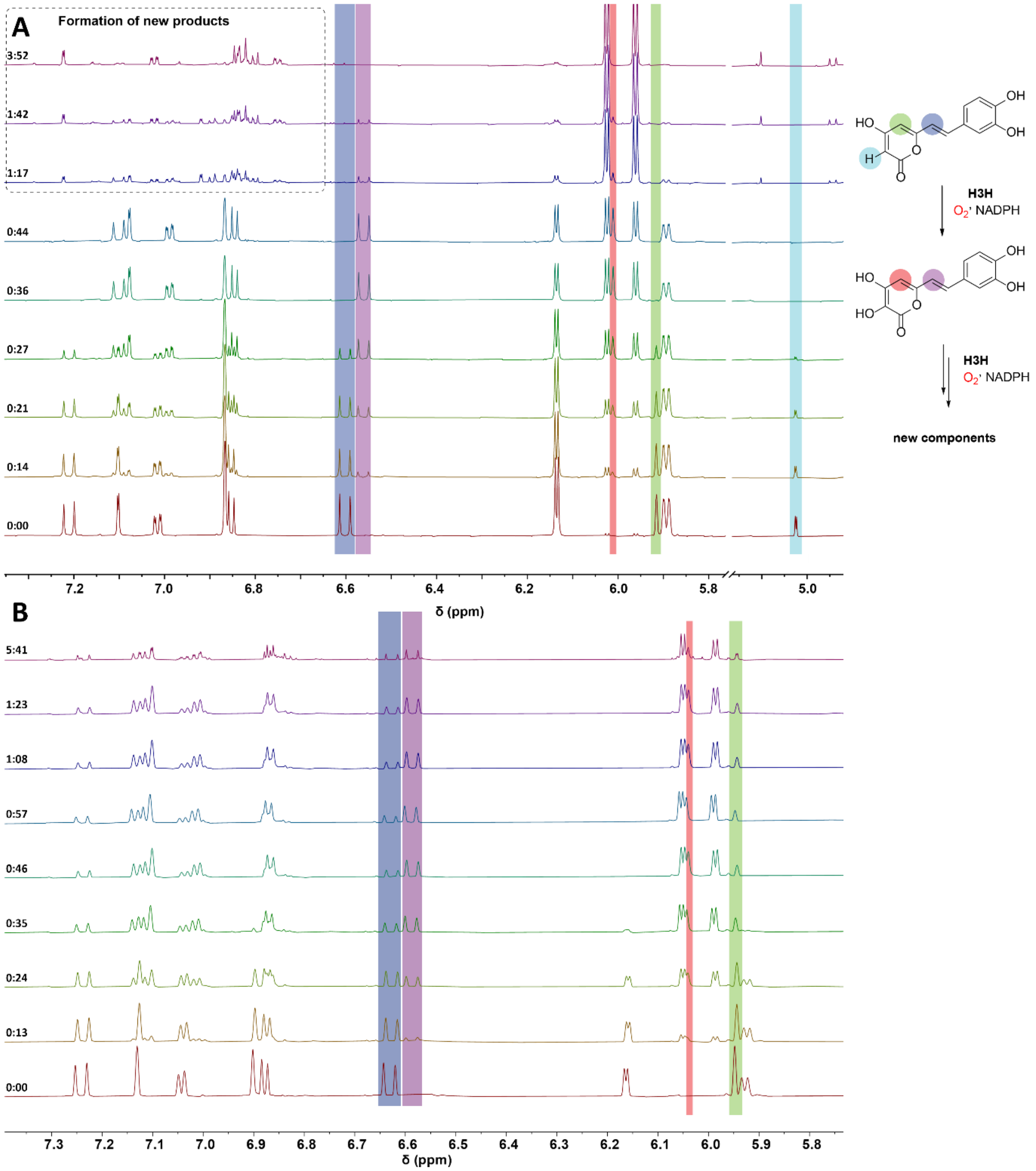


**Figure S12: [1]H NMR time course analysis of mcH3H-catalysed hispidin turnover at different hispidin:NADPH ratios.** Turnover of hispidin (250 µM) by mcH3H (375 nM) was monitored over time in the presence of FAD (500 nM) and either (**A**) NADPH (750 µM; 1:3 hispidin:NADPH) or (**B**) NADPH (250 µM; 1:1 hispidin:NADPH). Spectra prior to enzyme addition are shown at the bottom, with reaction time progressing upwards. Under excess NADPH conditions (**A**), initial formation of 3-hydroxyhispidin was followed by its depletion and formation of additional products, whereas near stoichiometric NADPH (**B**) resulted in accumulation of 3-hydroxyhispidin with limited subsequent turnover. Reactions were performed in 25 mM Tris-$d_{11}$, pH 8.0, containing 10% (v/v) $D_2O$. Spectra were acquired using a Bruker AVIII 700 MHz spectrometer with water presaturation. Resonances corresponding to hispidin and 3-hydroxyhispidin are highlighted according to the proton assignments shown. Time is shown as h:mm.

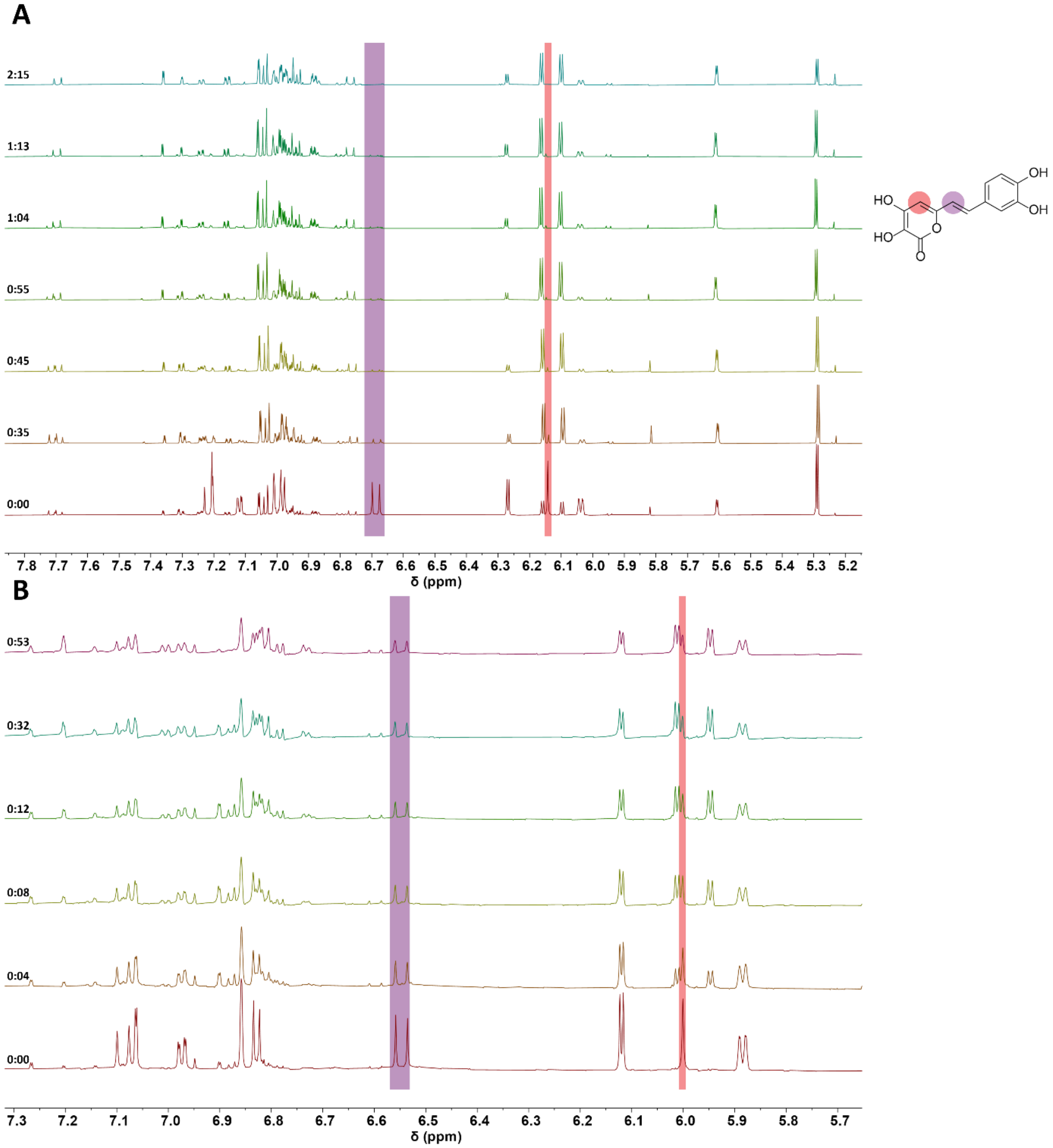


**Figure S13: $^1$H NMR time course analysis of mcH3H-catalysed 3-hydroxyhispidin turnover with and without NADPH recycling.** Turnover of 3-hydroxyhispidin by mcH3H was monitored over time (**A**) in the presence of an NADPH recycling system containing mcH3H (2.5 µM), 3-hydroxyhispidin (2 mM), NADPH (2 mM), FAD (15 µM), glucose (4 mM) and glucose dehydrogenase (1.6 ng $mL^{-1}$), or (**B**) without NADPH recycling using mcH3H (0.5 µM), 3-hydroxyhispidin (250 µM), NADPH (500 µM) and FAD (0.5 µM). Spectra prior to enzyme addition are shown at the bottom, with reaction time progressing upwards. Progressive depletion of 3-hydroxyhispidin and formation of additional products were observed under both conditions. Reactions were performed in 25 mM Tris-$d_{11}$, pH 8.0, containing 10% (v/v) $D_2O$. Spectra were acquired using a Bruker AVIII 700 MHz spectrometer with water presaturation. Resonances corresponding to 3-hydroxyhispidin are highlighted according to the proton assignments shown. Time is shown as h:mm.

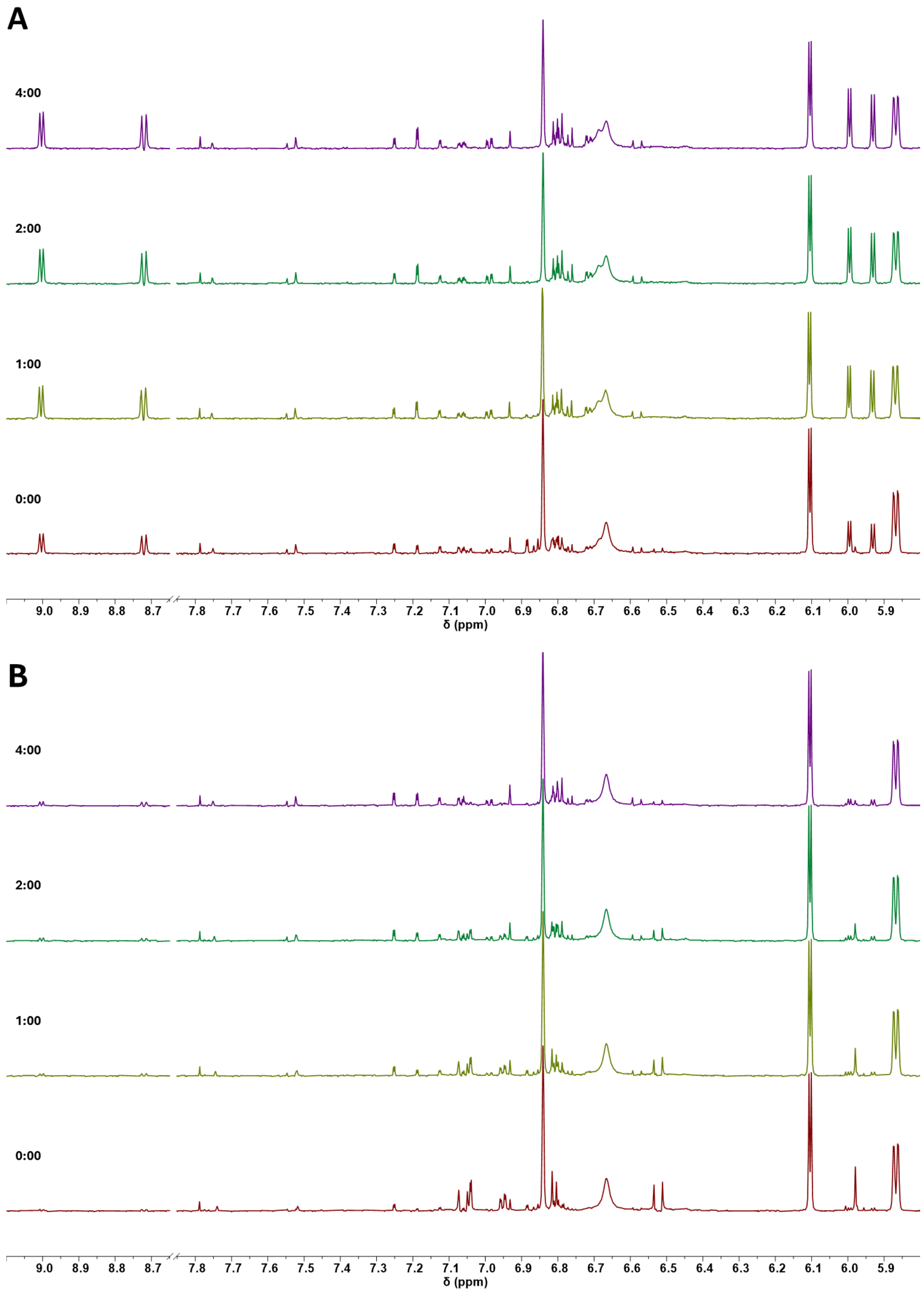


**Figure S14: Time-resolved $^{1}$H NMR control experiments of mcH3H-catalysed secondary 3-hydroxyhispidin turnover under different reaction conditions.** Stacked $^{1}$H NMR spectra recorded at 0 min, 1 h, 2 h and 4 h reaction with NADPH (750 µM) and FAD (2.5 µM) after addition of 3-hydroxyhispidin (250 µM) containing (**A**) mcH3H (0.25 µM), (**B**) heat-inactivated mcH3H (1.0 µM). The

1 h time point was recorded after incubating the sample in an Eppendorf tube exposed to air for 1 h. The sample was then transferred to an NMR tube, where subsequent time points were recorded under oxygen-limited conditions. Spectra were acquired in 25 mM Tris-$d_{11}$, 100 mM NaCl, pD 8.0 in $H_2O$/ $D_2O$ (9:1) using a Bruker AVIII 700 MHz spectrometer with water presaturation. Time is shown as h:mm.

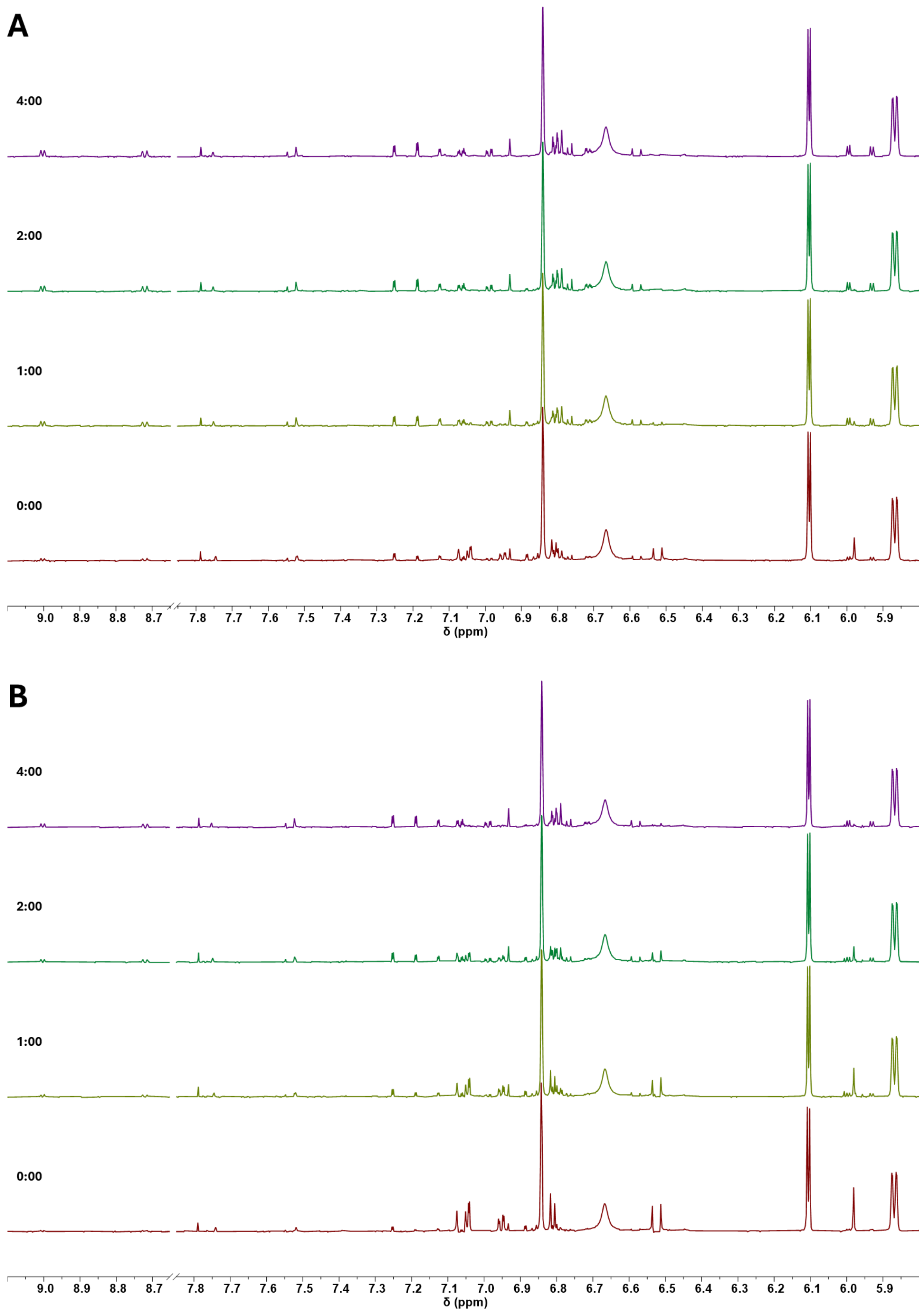


**Figure S15: Time-resolved $^1$H NMR control experiments of nnH3H-catalysed secondary 3-hydroxyhispidin turnover under different reaction conditions.** Stacked $^1$H NMR spectra recorded at 0 min, 1 h, 2 h and 4 h reaction with NADPH (750 µM) and FAD (2.5 µM) after addition of 3-

hydroxyhispidin (250 µM) containing (**A**) nnH3H (1.0 µM), (**B**) no enzyme. The 1 h time point was recorded after incubating the sample in an Eppendorf tube exposed to air for 1 h. The sample was then transferred to an NMR tube, where subsequent time points were recorded under oxygen-limited conditions. Spectra were acquired in 25 mM Tris-$d_{11}$, 100 mM NaCl, pD 8.0 in $H_2O$/ $D_2O$ (9:1) using a Bruker AVIII 700 MHz spectrometer with water presaturation. Time is shown as h:mm.

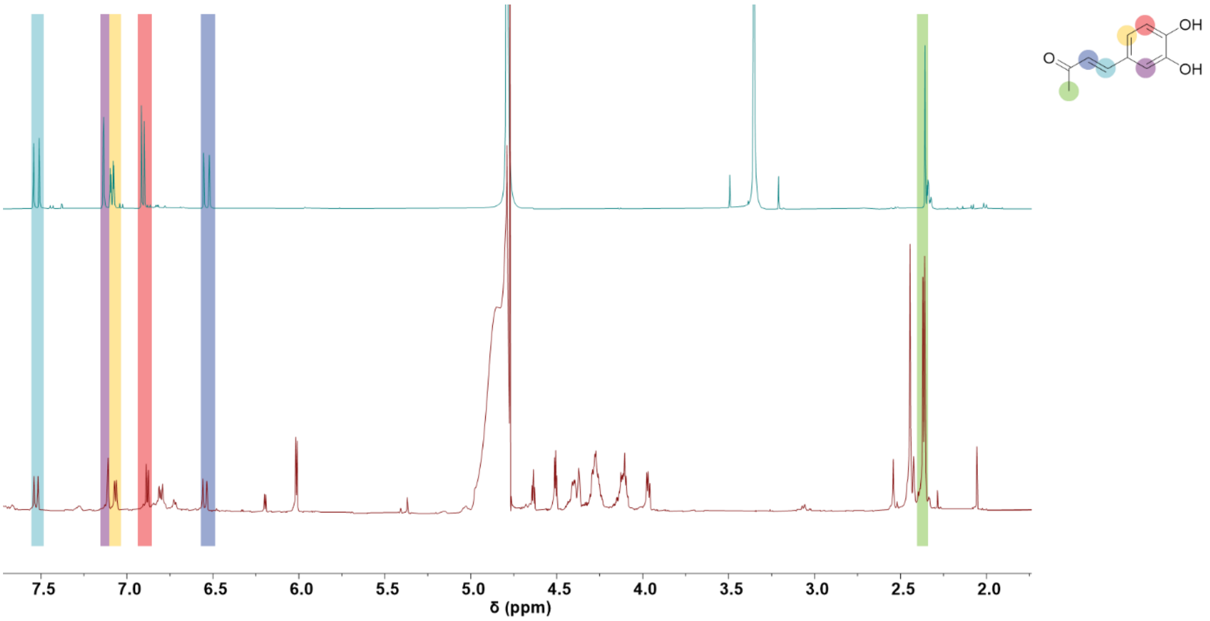


**Figure S16: [1]H NMR characterisation of compound 1 and comparison with a synthetic standard.** Compound **1** was isolated by HPLC following preparative mcH3H-catalysed turnover of 3-hydroxyhispidin and characterised by NMR and mass spectrometry. The $^{1}$H NMR spectrum of partially purified and isolated compound **1** (red) is shown alongside that of synthetic (3*E*)-4-(3,4 dihydroxyphenyl)-3-buten-2-one (blue), confirming the structural assignment. Synthetic procedure can be found in the experimental section. Spectra were acquired in $D_2O$ using a Bruker AVIII 700 MHz spectrometer with water presaturation. Resonances used for structural assignment are highlighted according to the proton positions indicated.

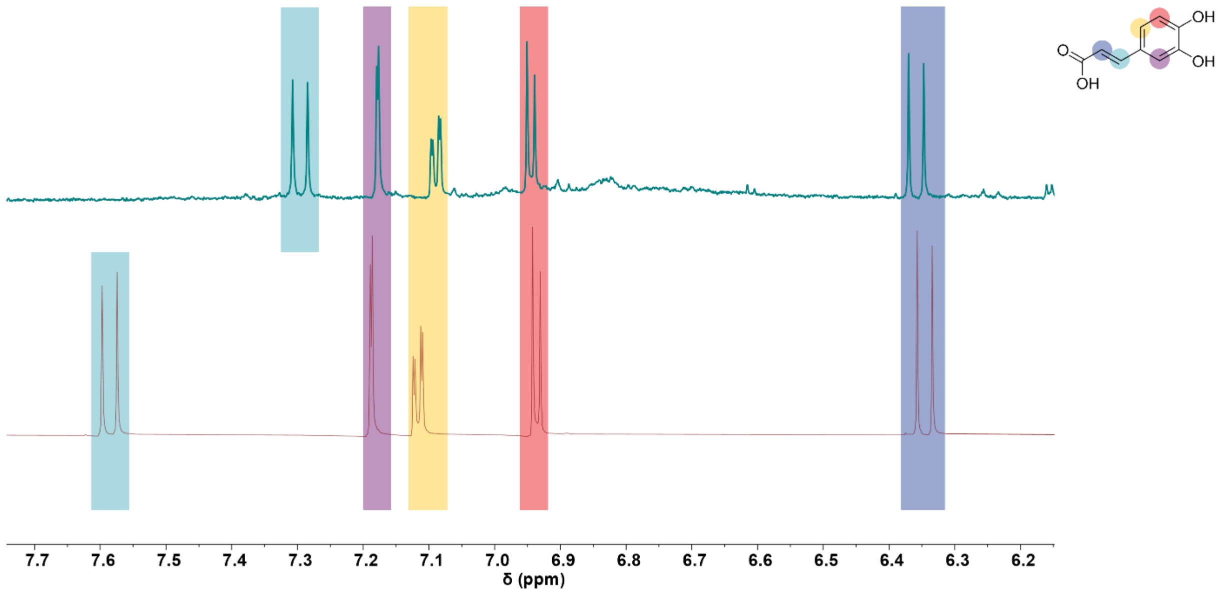


**Figure S17: [1]H NMR characterisation of compound 2 and comparison with authentic caffeic acid.** Compound **2** was isolated by HPLC following preparative mcH3H catalysed turnover of 3-hydroxyhispidin and characterised by NMR and LCMS. The $^1$H NMR spectrum of isolated compound **2** (blue) is shown alongside that of an authentic caffeic acid standard (red), supporting its assignment as caffeic acid. Minor differences in chemical shift are attributed to residual formic acid introduced during HPLC purification. Spectra were acquired in $D_2O$ using a Bruker AVIII 700 MHz spectrometer with water presaturation. Resonances used for structural assignment are highlighted according to the proton positions indicated.

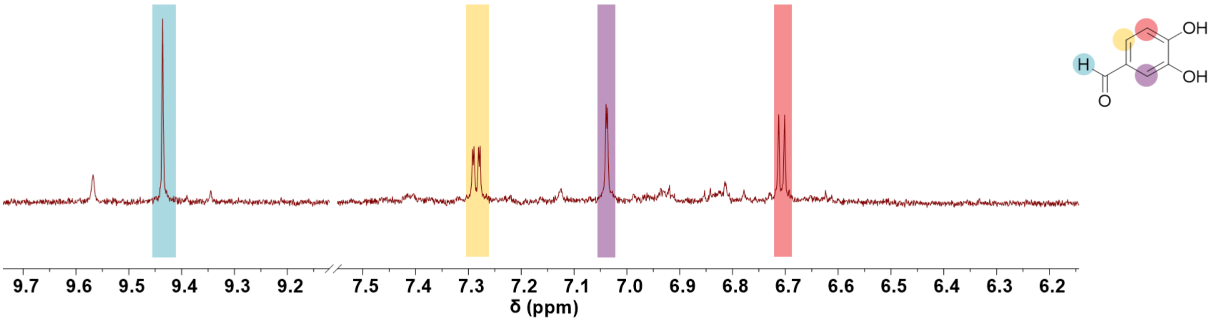


**Figure S18. $^{1}$H NMR characterisation of compound 3 isolated from secondary 3-hydroxyhispidin turnover.** Compound **3** was isolated by HPLC following preparative mcH3H-catalysed turnover of 3-hydroxyhispidin and characterised by $^{1}$H NMR, 2D NMR and high-resolution mass spectrometry, supporting its assignment as 3,4-dihydroxybenzaldehyde. The $^{1}$H NMR spectrum was acquired in $D_2O$ using a Bruker AVIII 700 MHz spectrometer. Resonances used for structural assignment are highlighted according to the proton positions indicated.

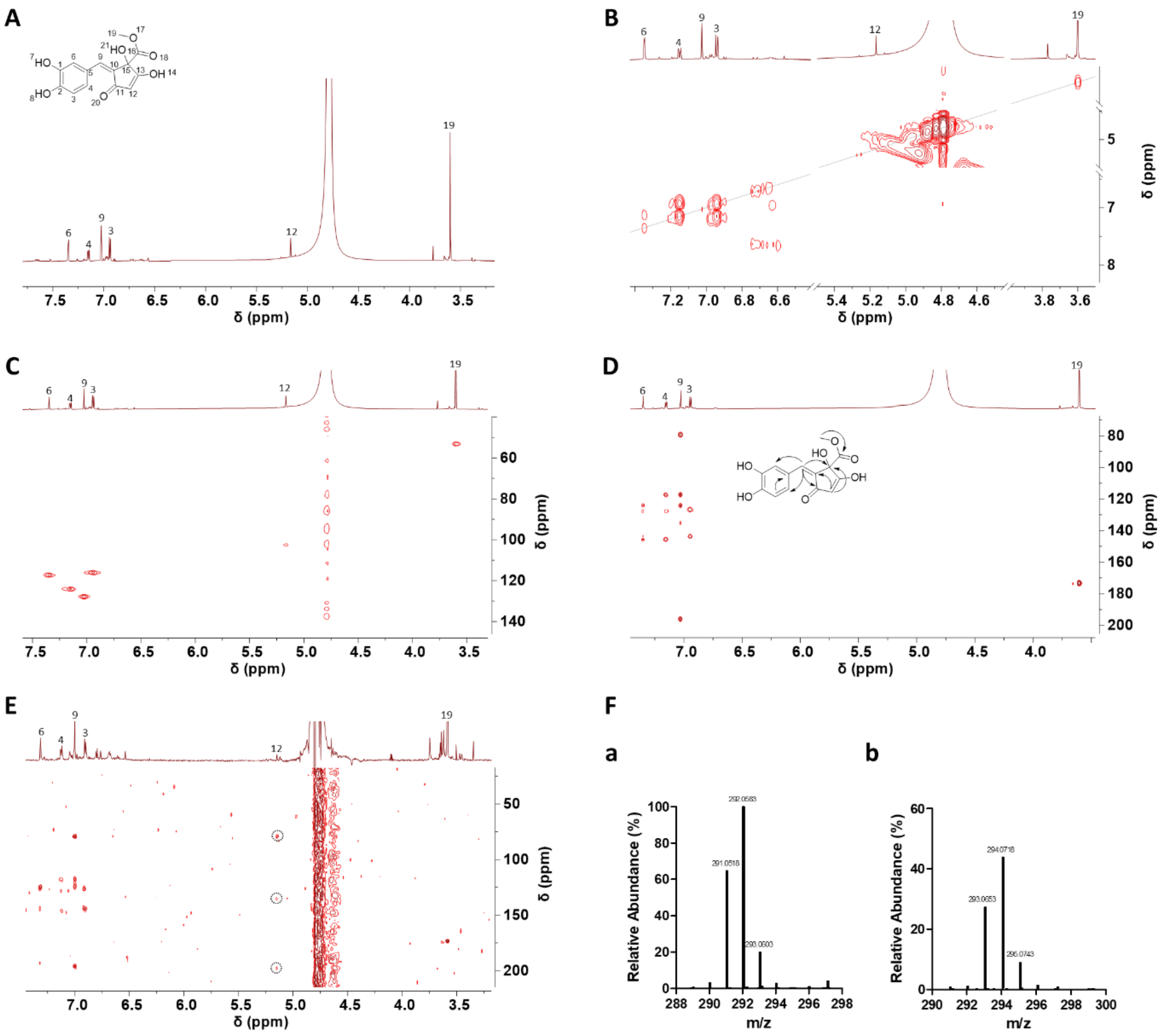


**Figure S19: NMR and high-resolution mass spectrometric characterisation of compound 4.** Compound **4** was characterised by (**A**) $^{1}H$ NMR, (**B**) $^{1}H,^{1}H$-COSY, (**C**) $^{1}H,^{13}C$-HSQC and (**D**) $^{1}H,^{13}C$-HMBC spectroscopy, with key correlations indicated obtained in $D_2O$. Note during the data collection the proton 12 exchanges to deuterium and therefore cross correlations are missing in the HMBC. (**E**) HMBC spectrum of a fraction containing compound **4** (and a second compound) in 90% $H_2O$/10% $D_2O$, enabling observation of correlations from the exchangeable proton 12 at $\delta_H$ 5.16. (**F**) High-resolution mass spectra of compound **4** acquired in (**a**) negative and (**b**) positive ionisation modes. The compound appears partially deuterated in the MS analysis. Spectra were acquired in $D_2O$ unless otherwise indicated. Atom assignments correspond to those shown in the structure. For full spectra please see Figure S27 and Figure S28.

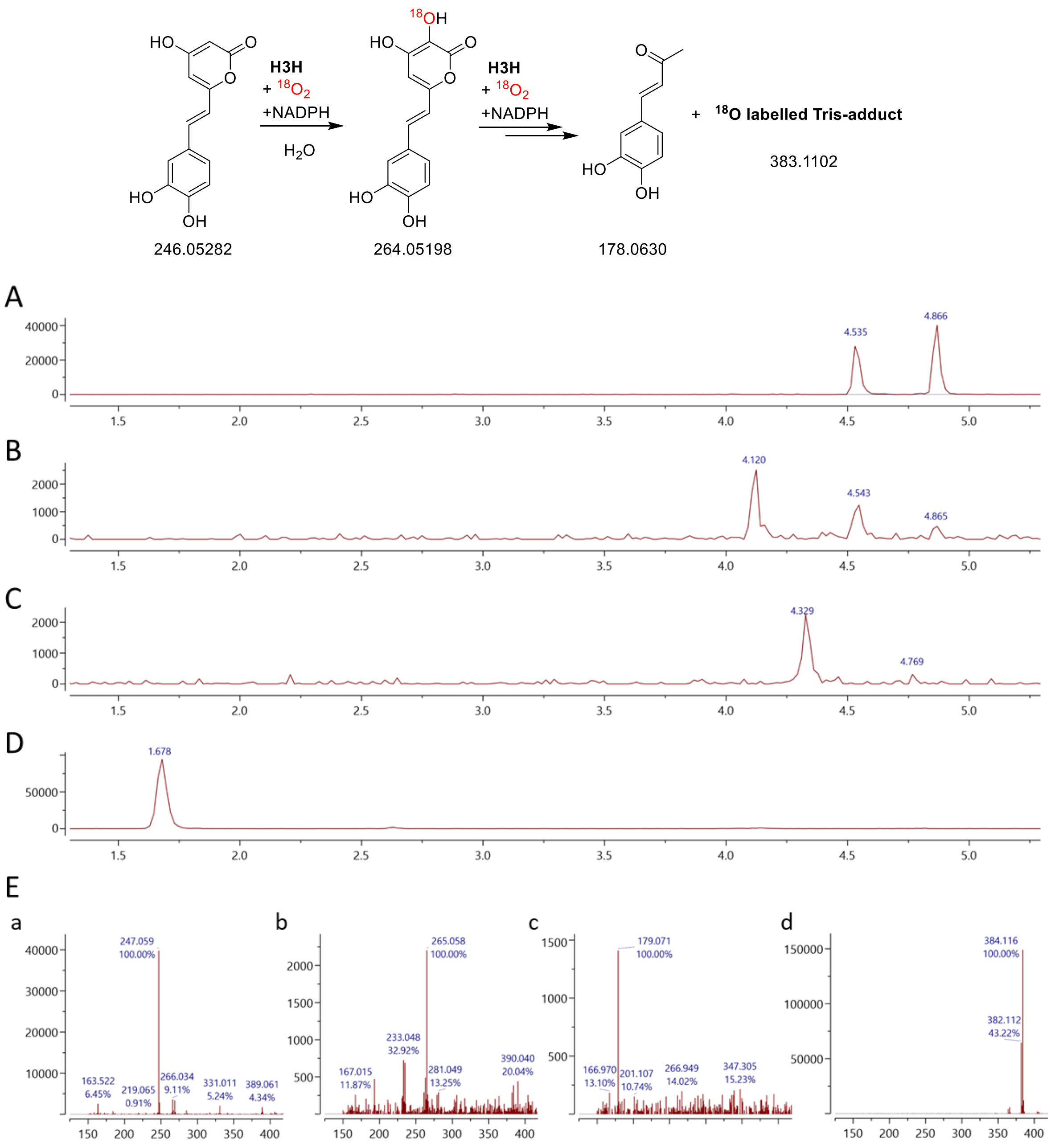


**Figure S20: $^{18}O_2$ labelling of mcH3H-catalysed hispidin turnover under stoichiometric NADPH conditions**. Hispidin (50 µM) and NADPH (50 µM; 1:1) were incubated with mcH3H (0.5 µM) in 50 mM Tris, 150 mM NaCl, pH 8.0 under anaerobic conditions before reaction initiation by exposure to $^{18}O_2$. Reactions were incubated for 2 h, quenched with 0.1% formic acid and analysed by LCMS. Extracted ion chromatograms show (**A**) m/z 247.1, hispidin; (**B**) m/z 265.1, $^{18}$O labelled 3-hydroxyhispidin; (**C**) m/z 179.1, compound **1**; (**D**) m/z 384.1, $^{18}$O labelled Tris-associated species. (**E**) Mass spectra corresponding to the indicated chromatographic peaks **(a)** 4.866**, (b)** 4.120, **(c)** 4.329**, (d)** 1.676 min. Extracted ion chromatograms were generated using a ± 0.25 m/z window. Scheme summarising the substrates and detected products, with observed $^{18}$O incorporation site indicated.

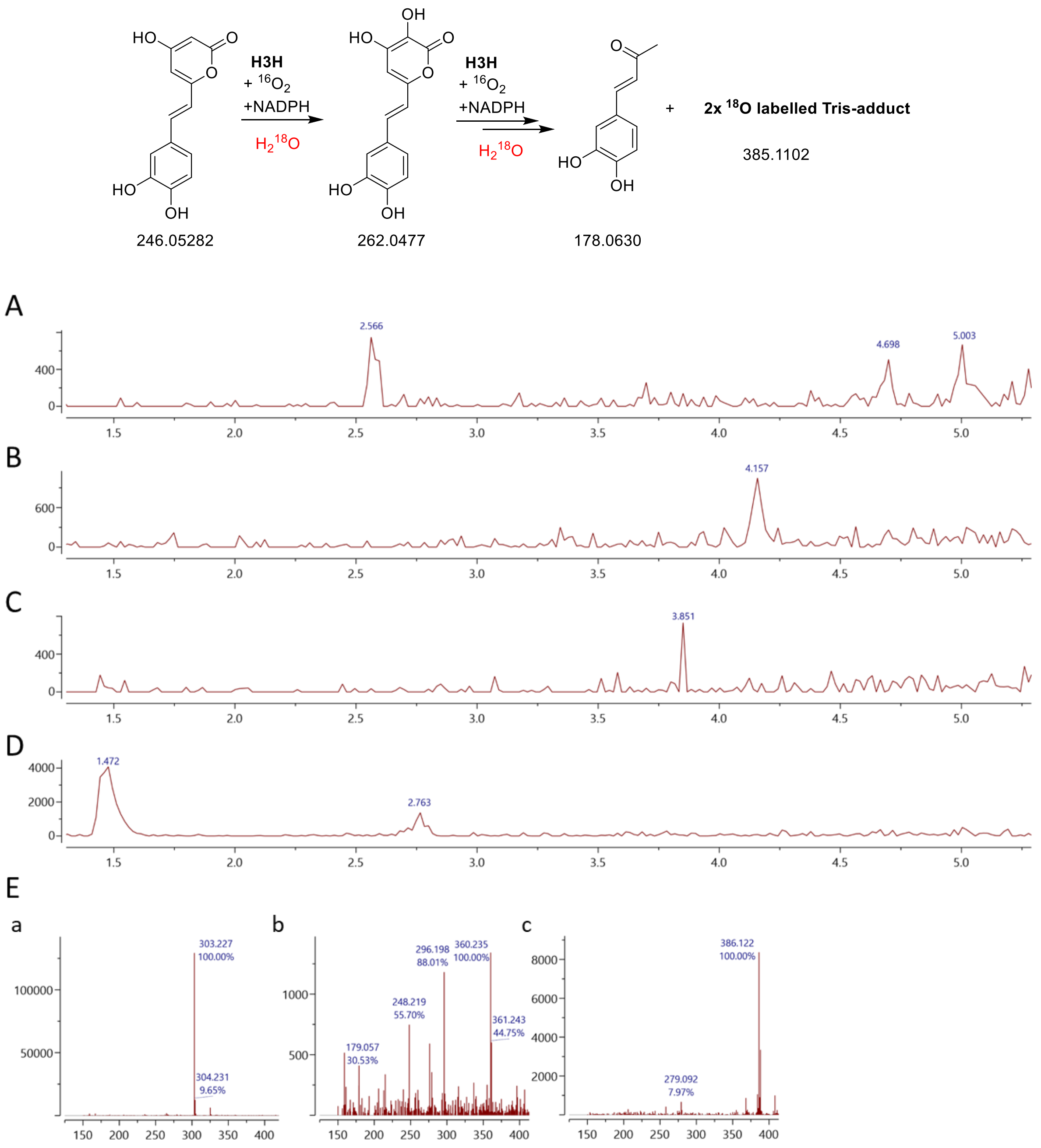


**Figure S21: $H_2{}^{18}O$ labelling of mcH3H-catalysed hispidin turnover under stoichiometric NADPH conditions**. Hispidin (50 µM) and NADPH (50 µM; 1:1) were incubated with mcH3H (0.5 µM) in 50 mM Tris, 150 mM NaCl, pH 8.0 prepared with $H_2{}^{18}O$. Reactions were incubated for 2 h, quenched with 0.1% formic acid and analysed by LCMS. Extracted ion chromatograms show (**A**) m/z 247.1, hispidin; (**B**) m/z 265.1, $^{18}O$ labelled 3-hydroxyhispidin; (**C**) m/z 179.1, compound **1**; (**D**) m/z 384.1, $^{18}O$ labelled Tris-associated species. (**E**) Mass spectra corresponding to the indicated chromatographic peaks **(a)** 4.157**, (b)** 3.851, **(c)** 1.472. Extracted ion chromatograms were generated using a ± 0.25 m/z window. Scheme summarising the substrates and detected products, with observed $^{18}O$ incorporation site indicated.

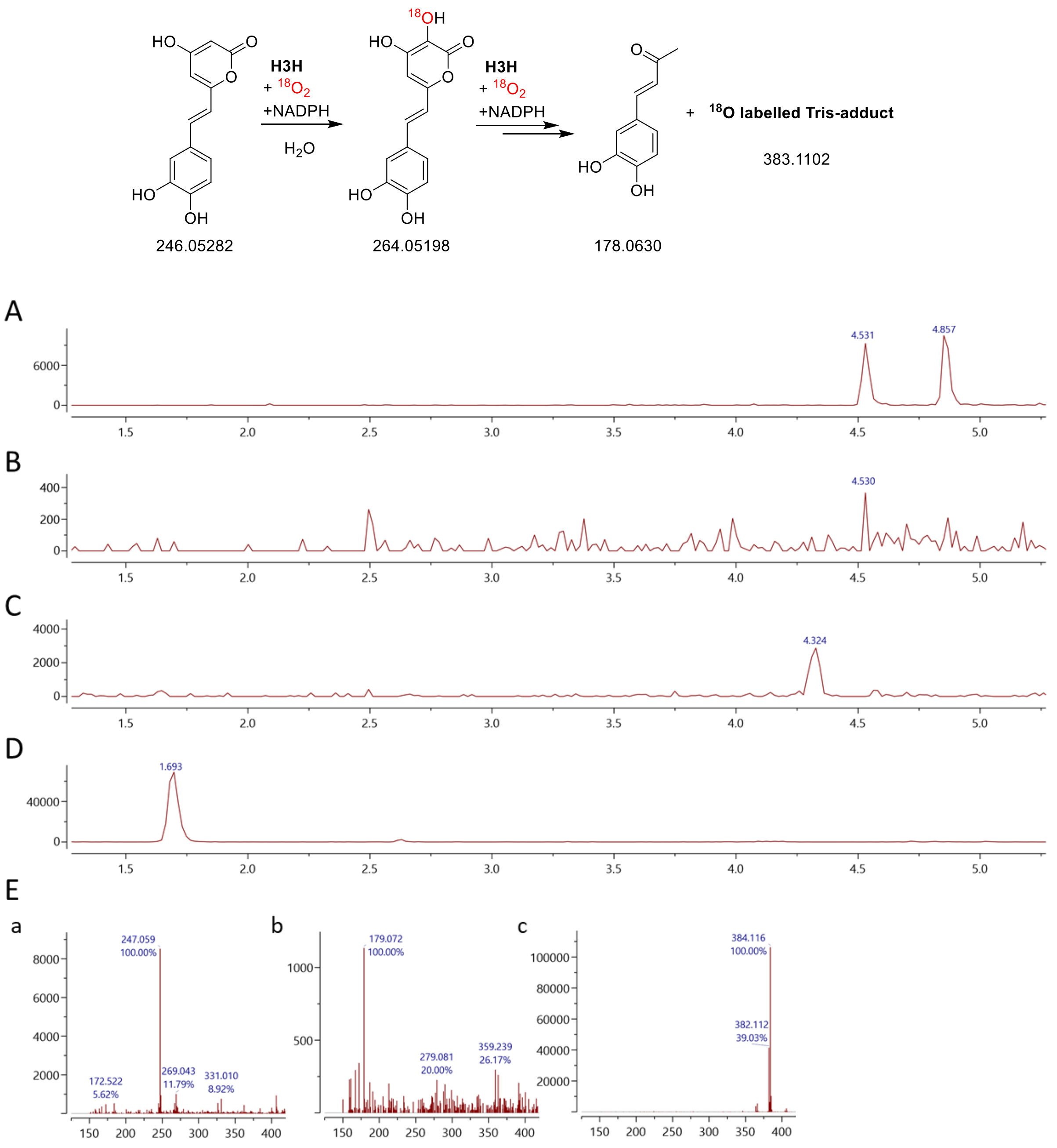


**Figure S22: $^{18}O_2$ labelling of mcH3H-catalysed hispidin turnover under excess NADPH conditions**. Hispidin (50 µM) and NADPH (150 µM; 1:3) were incubated with mcH3H (0.5 µM) in 50 mM Tris, 150 mM NaCl, pH 8.0 under anaerobic conditions before reaction initiation by exposure to $^{18}O_2$. Reactions were incubated for 2 h, quenched with 0.1% formic acid and analysed by LCMS. Extracted ion chromatograms show (**A**) m/z 247.1, hispidin; (**B**) m/z 265.1, $^{18}$O labelled 3-hydroxyhispidin (very minor); (**C**) m/z 179.1, compound **1**; (**D**) m/z 384.1, $^{18}$O labelled Tris associated species. (**E**) Mass spectra corresponding to the indicated chromatographic peaks **(a)** 4.857**, (b)** 4.324, **(c)** 1.693. Extracted ion chromatograms were generated using a ± 0.25 m/z window. Scheme summarising the substrates and detected products, with observed $^{18}$O incorporation site indicated.

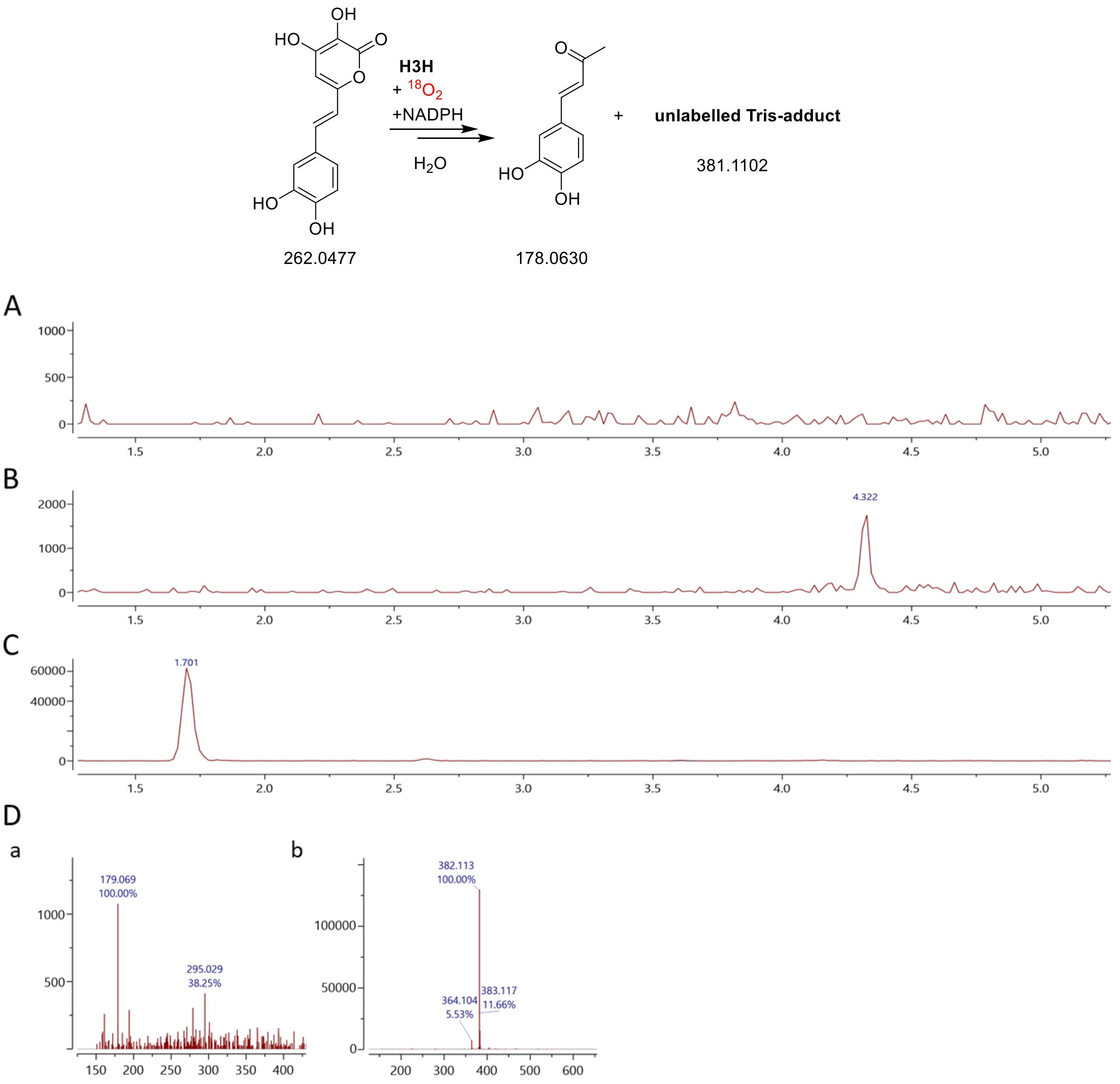


**Figure S23: $^{18}O_2$ labelling of mcH3H catalysed 3-hydroxyhispidin turnover under excess NADPH conditions**. 3-hydroxyhispidin (50 µM) and NADPH (100 µM; 1:2) were incubated with mcH3H (0.5 µM) in 50 mM Tris, 150 mM NaCl, pH 8.0 under anaerobic conditions before reaction initiation by exposure to $^{18}O_2$. Reactions were incubated for 2 h, quenched with 0.1% formic acid and analysed by LCMS. Extracted ion chromatograms show (**A**) m/z 263.1, 3-hydroxyhispidin; (**B**) m/z 179.1, compound **1**; (**C**) m/z 382.1, Tris-associated species. (**D**) Mass spectra corresponding to the indicated chromatographic peaks **(a)** 4.322**, (b)** 1.701. Extracted ion chromatograms were generated using a ± 0.25 m/z window. Scheme summarising the substrates and detected products, with observed $^{18}O$ incorporation site indicated.

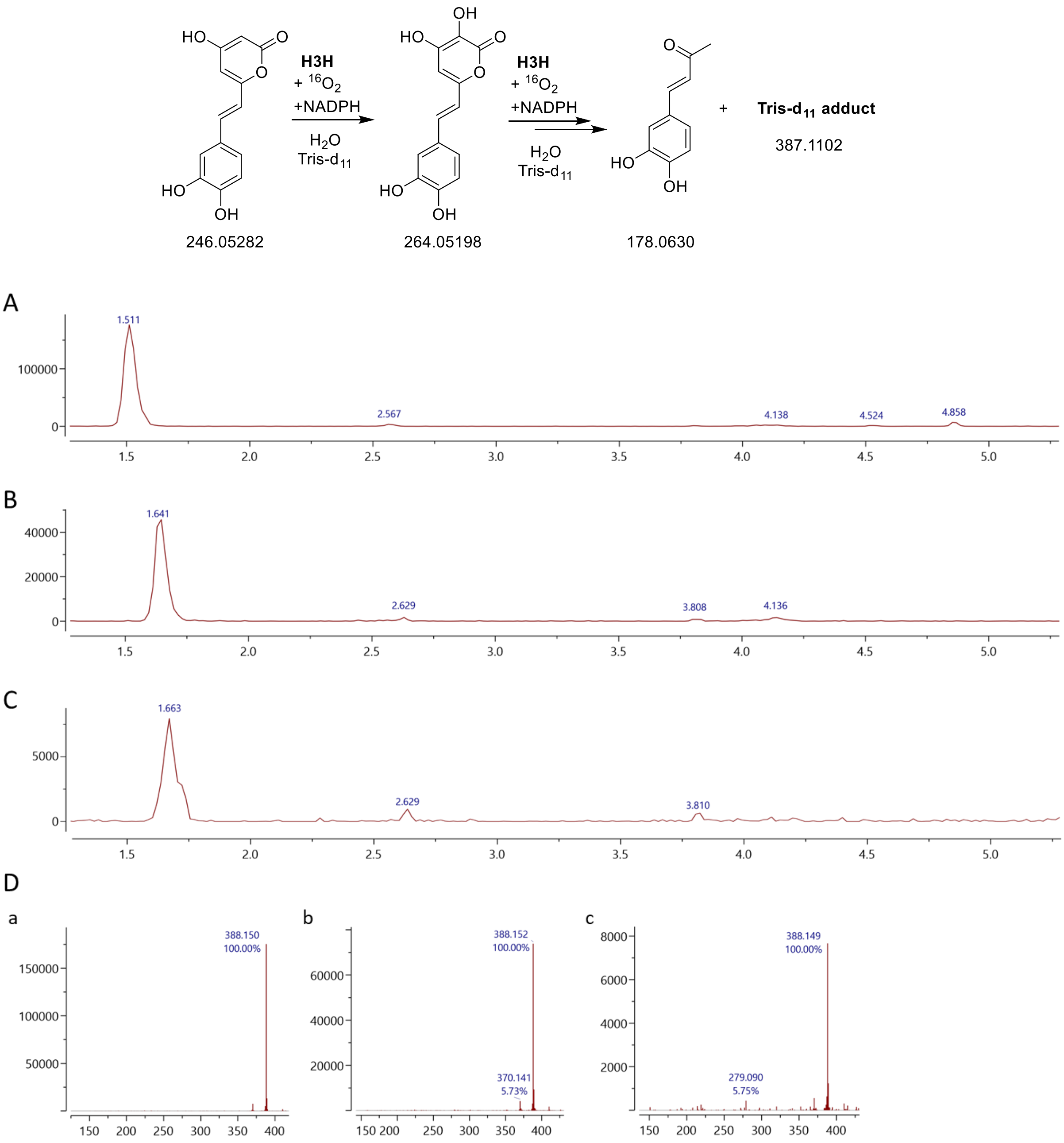


**Figure S24: Tris-d$_{11}$ labelling of mcH3H-catalysed hispidin and 3-hydroxyhispidin turnover under stoichiometric NADPH conditions**. Substrates and NADPH were incubated with mcH3H (0.5 µM) in 50 mM Tris-d$_{11}$, 150 mM NaCl, pH 8.0. The shift of the Tris-associated species from m/z 382.113 to m/z 388.150 is consistent with retention of the six non-exchangeable deuterium atoms of Tris-d$_{11}$, supporting incorporation of a Tris-derived component into this species. Reactions were incubated for 2 h, quenched with 0.1% formic acid and analysed by LCMS. Extracted ion chromatograms show m/z 388.1, Tris-d$_{11}$-associated species from reactions (**A**) hispidin (50 µM) and NADPH (50 µM; 1:1), (**B**) hispidin (50 µM) and NADPH (150 µM; 1:3) and (**C**) 3-hydroxyhispidin (50 µM) and NADPH (100 µM; 1:2). (**D**) Mass spectra corresponding to the indicated chromatographic peaks **(a)** 1.511**, (b)** 1.641, **(c)** 1.663 min. Extracted ion chromatograms were generated using a ± 0.25 m/z window. The scheme summarises the substrates and detected products, with retention of the six non-exchangeable deuterium atoms from Tris-d$_{11}$ indicated.

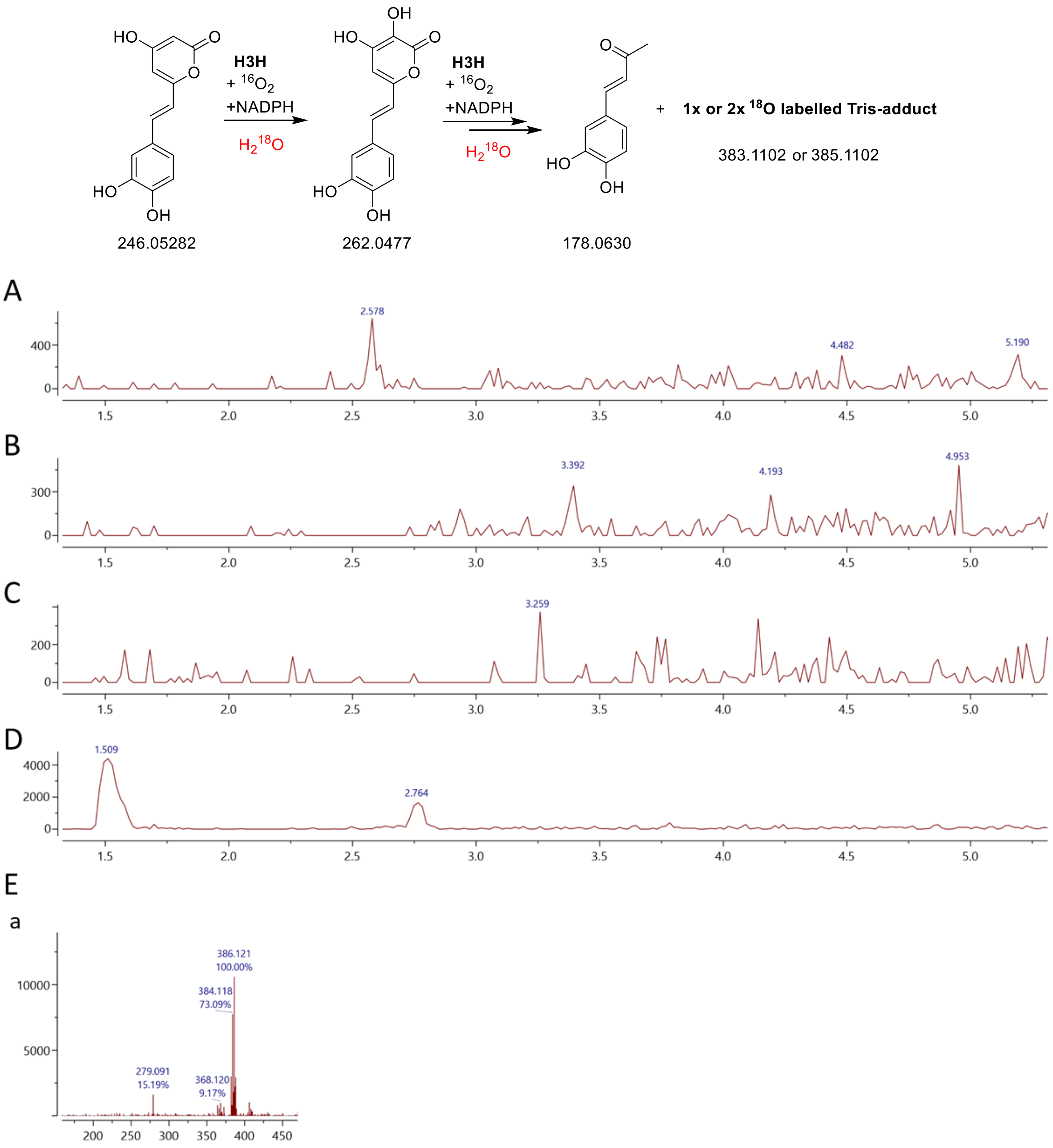


**Figure S25: $H_2{}^{18}O$ labelling of mcH3H-catalysed hispidin turnover under excess NADPH conditions**. Hispidin (50 µM) and NADPH (150 µM; 1:3) were incubated with mcH3H (0.5 µM) in 50 mM Tris, 150 mM NaCl, pH 8.0 prepared with $H_2{}^{18}O$. Reactions were incubated for 2 h, quenched with 0.1% formic acid and analysed by LCMS. Extracted ion chromatograms show (**A**) m/z 247.1, hispidin; (**B**) m/z 265.1, $^{18}O$ labelled 3-hydroxyhispidin; (**C**) m/z 179.1, compound **1**; (**D**) m/z 384.1, $^{18}O$ labelled Tris-associated species. (**E**) Mass spectra corresponding to the indicated chromatographic peaks **(a)** 1.509. Extracted ion chromatograms were generated using a ± 0.25 m/z window. Scheme summarising the substrates and detected products, with observed $^{18}O$ incorporation site indicated.

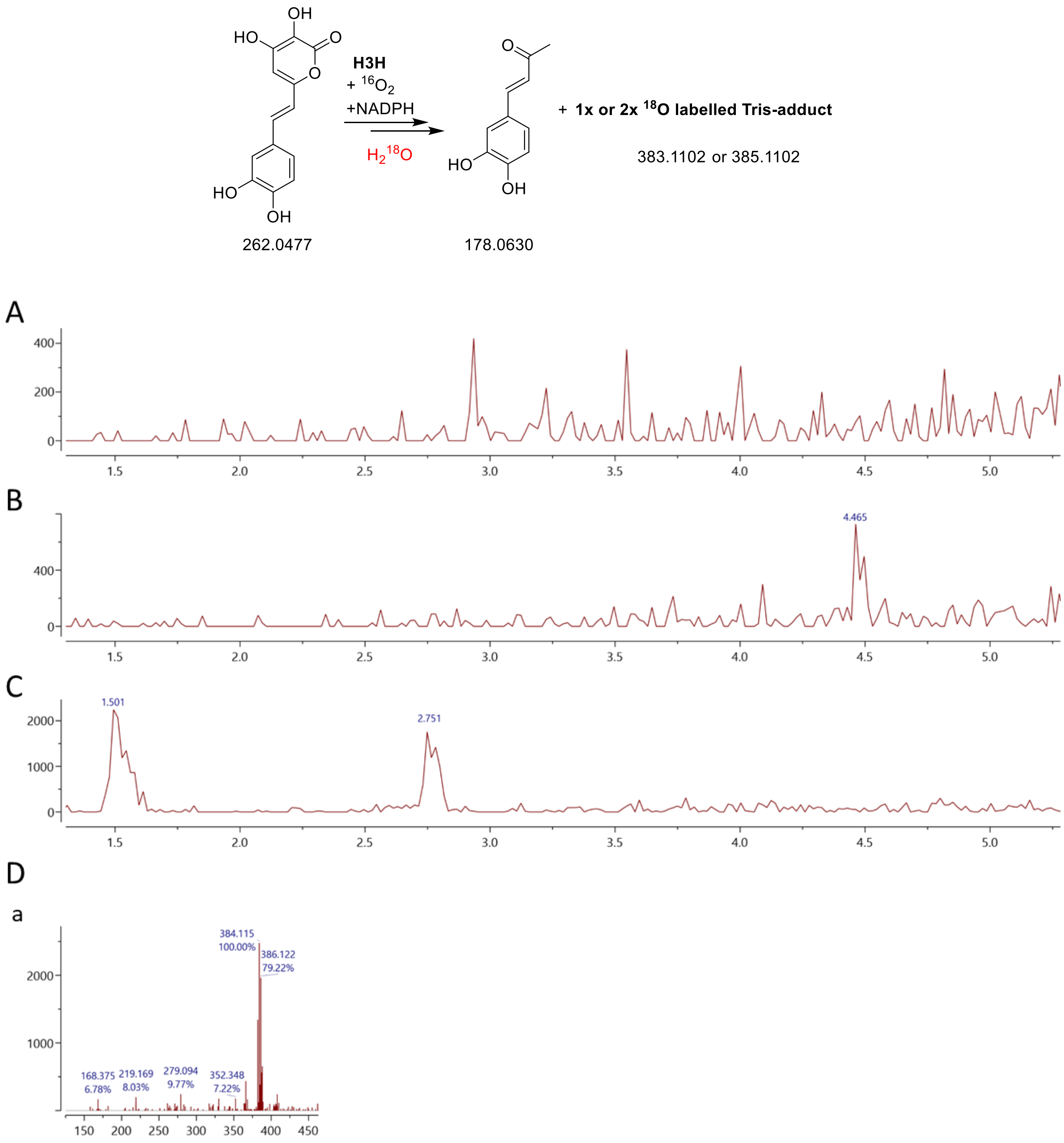


**Figure S26: $H_2{}^{18}O$ labelling of mcH3H-catalysed 3-hydroxyhispidin turnover under excess NADPH conditions**. 3-hydroxyhispidin (50 µM) and NADPH (100 µM; 1:2) were incubated with mcH3H (0.5 µM) in 50 mM Tris, 150 mM NaCl, pH 8.0 prepared with $H_2{}^{18}O$. Reactions were incubated for 2 h, quenched with 0.1% formic acid and analysed by LCMS. Extracted ion chromatograms show (**A**) m/z 247.1, hispidin (not present); (**B**) m/z 265.1, $^{18}O$ labelled 3-hydroxyhispidin (not present); (**C**) m/z 179.1, compound **1**; (**D**) m/z 384.1, $^{18}O$ labelled Tris-associated species. (**E**) Mass spectra corresponding to the indicated chromatographic peaks **(a)** 1.501. Extracted ion chromatograms were generated using a ± 0.25 m/z window. Scheme summarising the substrates and detected products, with observed $^{18}O$ incorporation site indicated.

**A**

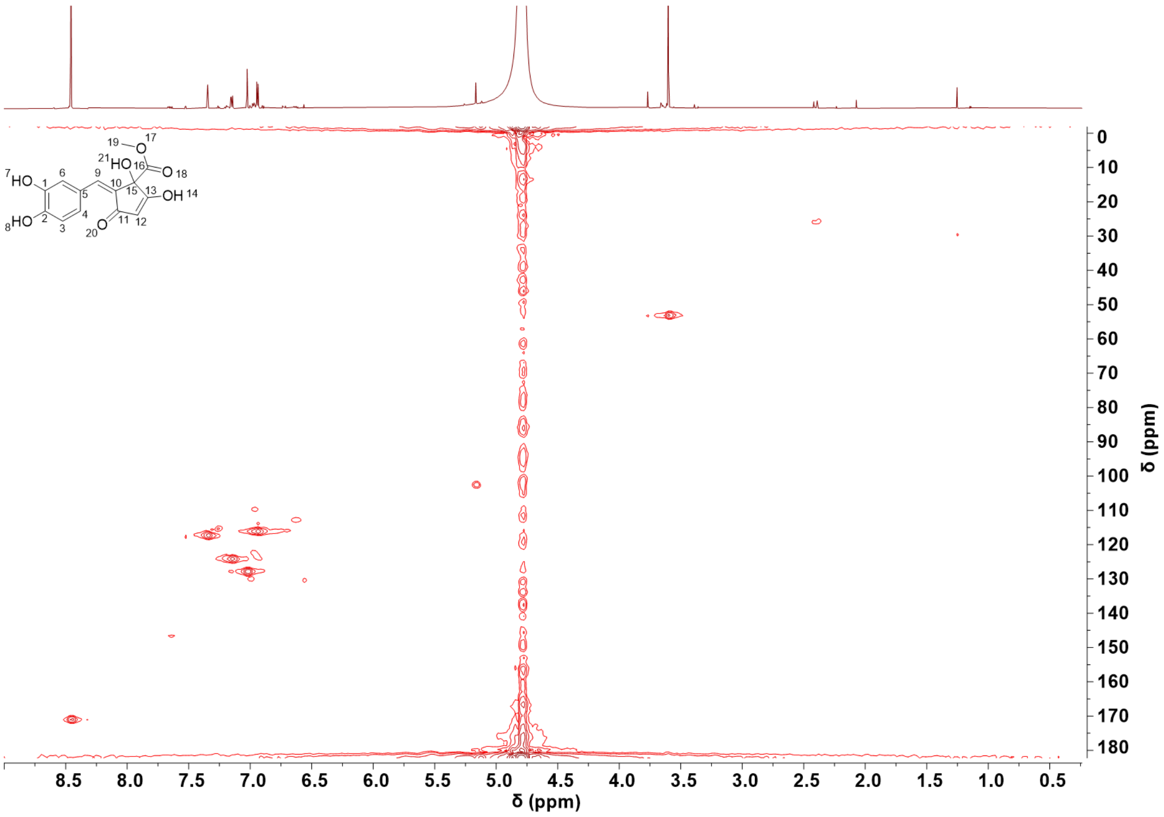


**B**

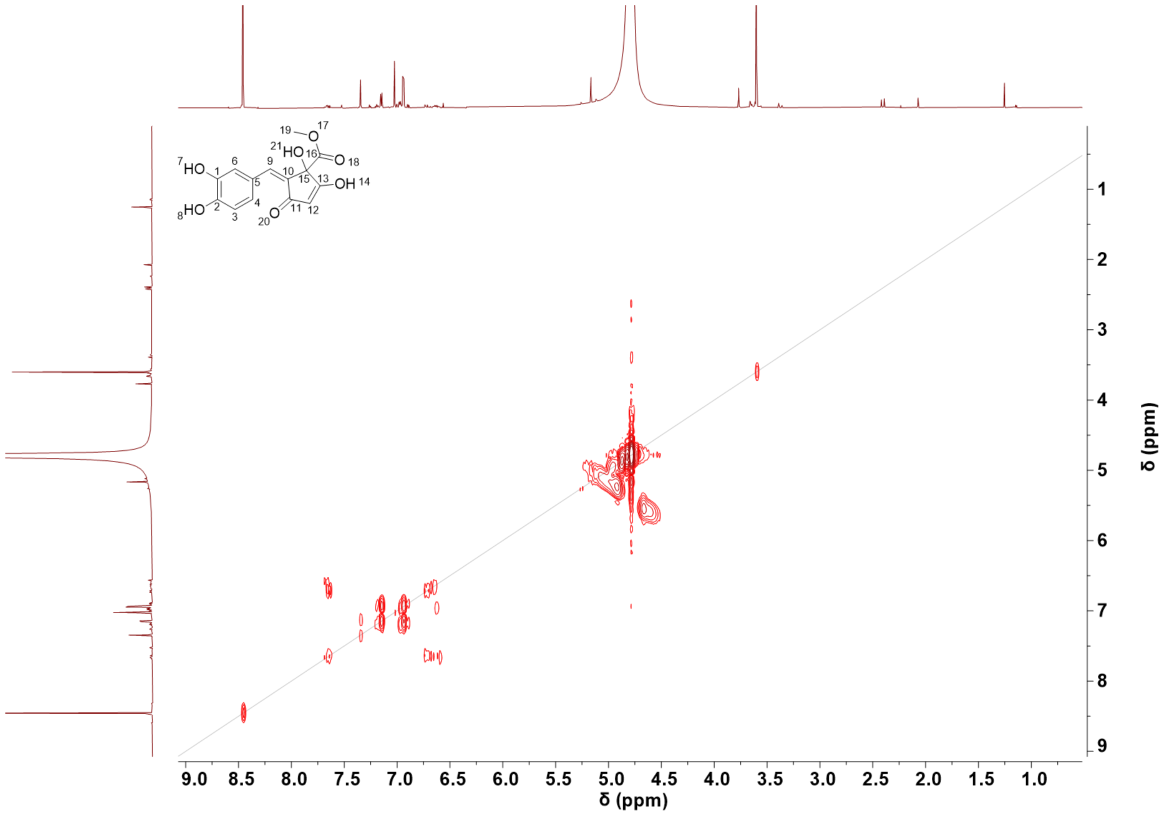


**Figure S27: HSQC and COSY NMR spectra of compound 4.** (**A**) HSQC and (**B**) COSY Spectra were acquired in $D_2O$ using a Bruker AVIII 700 MHz spectrometer with water presaturation.

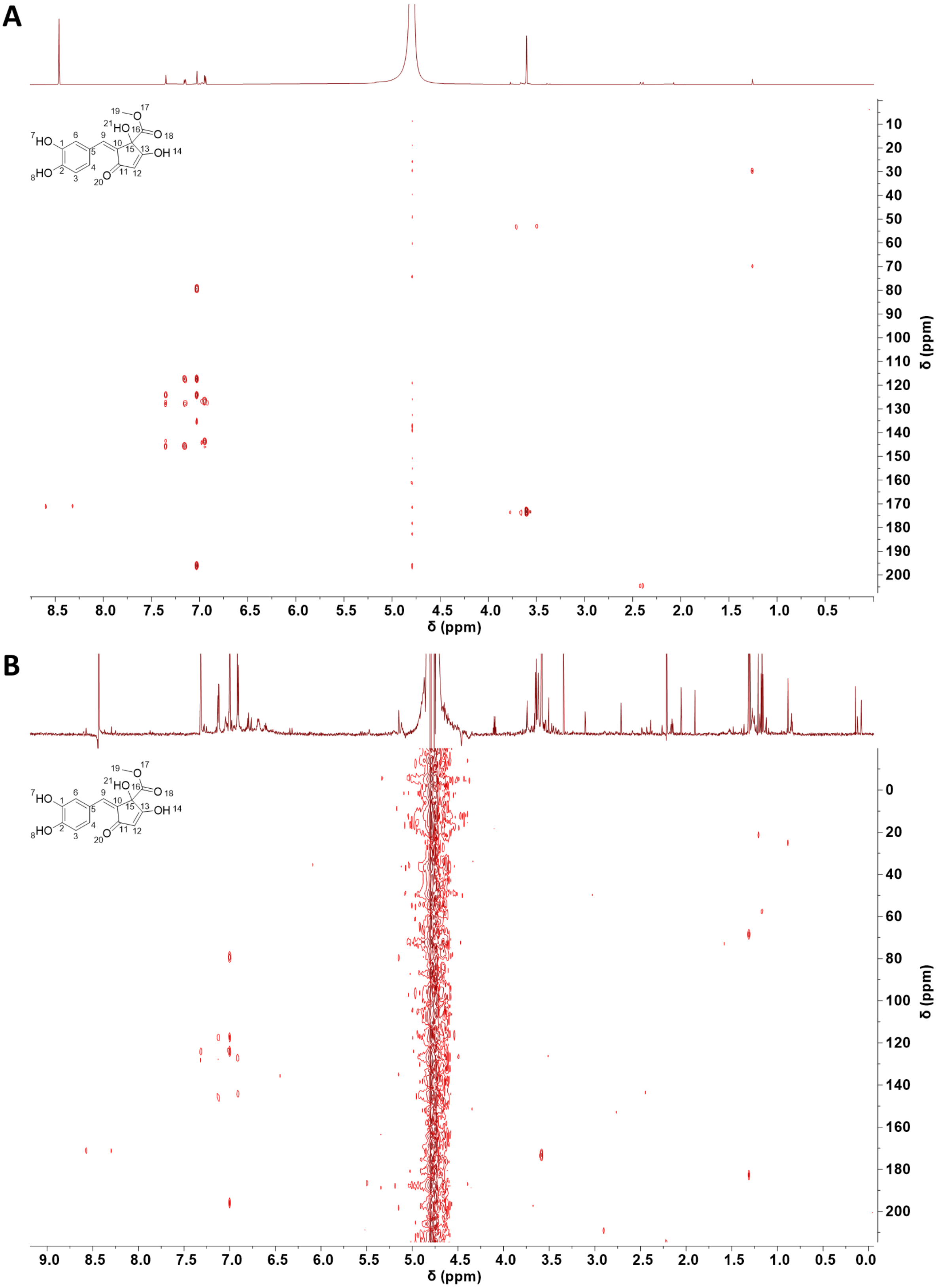


**Figure S28: HMBC NMR spectra of compound 4.** (**A**) HMBC of isolated **4** in $D_2O$, (**B**) HMBC of partially purified **4** in 90% $H_2O$ 10% $D_2O$ to limit exchange of singlet proton $\delta_H$ 5.16 (1H). Spectra were acquired in $D_2O$ using a Bruker AVIII 700 MHz spectrometer with water presaturation.

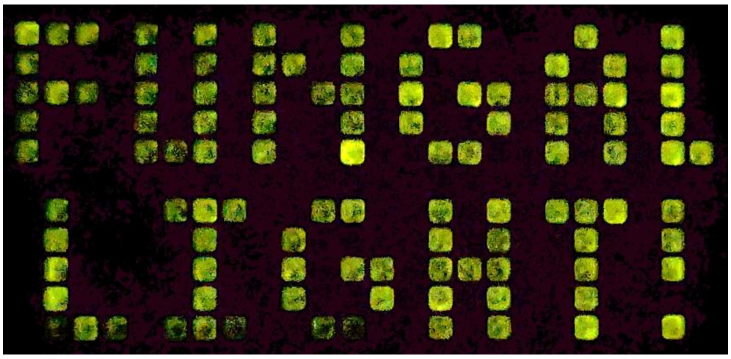

**Figure S29: 384-well format light production experiment**. Still image from timelapse video of reaction between 3-hydroxyhispidin (0.5 µM) with lysed *E. coli* (C43) cells expressing $Luz_{Tr}$ in a total volume of 100 µL prepared in 100 mM sodium phosphate, 100 mM NaCl, pH8.

**Table S1. Key NMR correlations for compound 4.** Assignments based on COSY, HSQC, HMBC in $D_2O$, HMBC repeated in 90% $H_2O$, 10% $D_2O$ due to exchange of **5**. Original isolated compound was exhausted during analysis therefore **5**'s cross peaks were determined from a partially purified fractions containing a mixture of compounds.  Chemical shifts in (δ) in ppm. s, singlet; d, doublet; dd, doublet of doublets. High-resolution mass spectrometry: Calculated for $C_{14}H_{12}O_7$ $[M+H]^+$ 293.0656, found 293.0653.

| Assignment | δH (ppm) | Multiplicity | Integration | J (Hz) | δC (ppm) | COSY | HMBC |
|---|---|---|---|---|---|---|---|
| **1** | 7.35 | d | 1H | 2.0 | 117.3 | 2 (weak) | 124.1, 127.8, 143.6, 145.6 |
| **2** | 7.15 | dd | 1H | 2.0, 8.3 | 124.1 | 4, 1 (weak) | 117.3, 127.8, 145.5 |
| **3** | 7.05 | s | 1H | - | 127.7 | - | 195.6, 135.4, 117.3, 124.1, 79.3 |
| **4** | 6.94 | d | 1H | 8.3 | 116.1 | 2 | 126.7, 143.6, 145.5 (weak) |
| **5** | 5.16 | s | 1H | - | 102.5 | - | 79.3, 135.4, 198.1 |
| **6** | 3.60 | s | 3H | - | 53.1 | - | 173.1 |

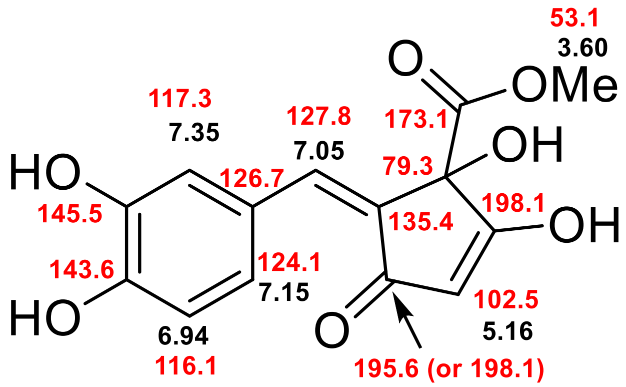

**Table S2. Primers used in this study**

| | Primer Sequences |
|---|---|
| **AOX1-3'** | 5'-GCAAATGGCATTCTGACATCC-3' |
| **AOX1-5'** | 5'GACTGGTTCCAATTGACAAGC-3' |
| **pCold-fwd** | 5'-ACGCCATATCGCCGAAAGG-3' |
| **pCold-rev** | 5'-GGCAGGGATCTTAGATTCTG-3' |
| **T7-Term** | 5'-GCTAGTTATTGCTCAGCGG-3' |
| **pCOLI_A_SUMO_fwd** | 5'-ATATATGAGCTCTCGGACTCAGAAGTCAATCAAGAAGCTAAG-3' |
| **Luzv4_Gibson_V_fwd** | 5'-CAAGCGCAATTGCATTTCCGATTATTCGTCGTGATTATCAGACCTTTCT-3' |
| **Luzv4_Gibson_V_rev** | 5'- AGGCTAATGTTAATACGCATGTCGACCACTTTGTGATTCATGG-3' |
| **Pcola_Gibson_1-37_fwd** | 5'- TGAATCACAAAGTGGTCGACATGCGTATTAACATTAGCCTG-3' |
| **Pcola_Gibson_1-37_rev** | 5'-TGATAATCACGACGAATAATCGGAAATGCAAT-3' |

The truncated luciferase sequence was cloned into a pCOLI_A_SUMO vector using SalI and NotI sites. This N-terminal SUMOlyated Luz constructs was then cloned into the custom pCold vectors. PCR with Q5 polymerase (NEB) was used to add a 5' SacI restriction site with T7 terminator primer at the 5' end of the SUMO. Subsequent SacI and NotI restriction digestion and ligation enabled production of the final N-terminal SUMO (38-267) $Luz_{Tr}$. Subsequent mcH3H and nnH3H were then cloned into this construct via SalI and NotI sites with no further modification. The C-terminal SUMO $Luz_{Tr}$ v4 construct was purchased using Thermofisher's GeneArt service with SUMO attached. The C-terminal SUMO full-length v4 construct was produced via Gibson assembly to reinstall AA 1-37 from our existing wt Luz construct, resulting in a variant without the first 2 mutations in the putative transmembrane domain.
